# Electrophoresis-Based Approach for Characterizing Dendrimer-Protein Interactions: A Proof-of-Concept Study

**DOI:** 10.1101/2023.08.31.555747

**Authors:** Simone A. Douglas-Green, Juan A. Aleman, Paula T. Hammond

## Abstract

Improving the clinical translation of nanomedicine requires better knowledge about how nanoparticles interact with biological environments. As researchers are recognizing the importance of understanding the protein corona and characterizing how nanocarriers respond in biological systems, new tools and techniques are needed to analyze nanocarrier-protein interactions, especially for smaller-size (<10nm) nanoparticles like polyamidoamine (PAMAM) dendrimers. Here we developed a streamlined, semi-quantitative approach to assess dendrimer-protein interactions using a non-denaturing electrophoresis technique combined with mass spectrometry. With this protocol, we detect fluorescently tagged dendrimers and proteins simultaneously, enabling us to analyze when dendrimers migrate with proteins. We found PAMAM dendrimers mostly interact with complement proteins, particularly C3 and C4a, which aligns with previously published data, verifying our approach can be used to isolate and identify dendrimer-protein interactions.

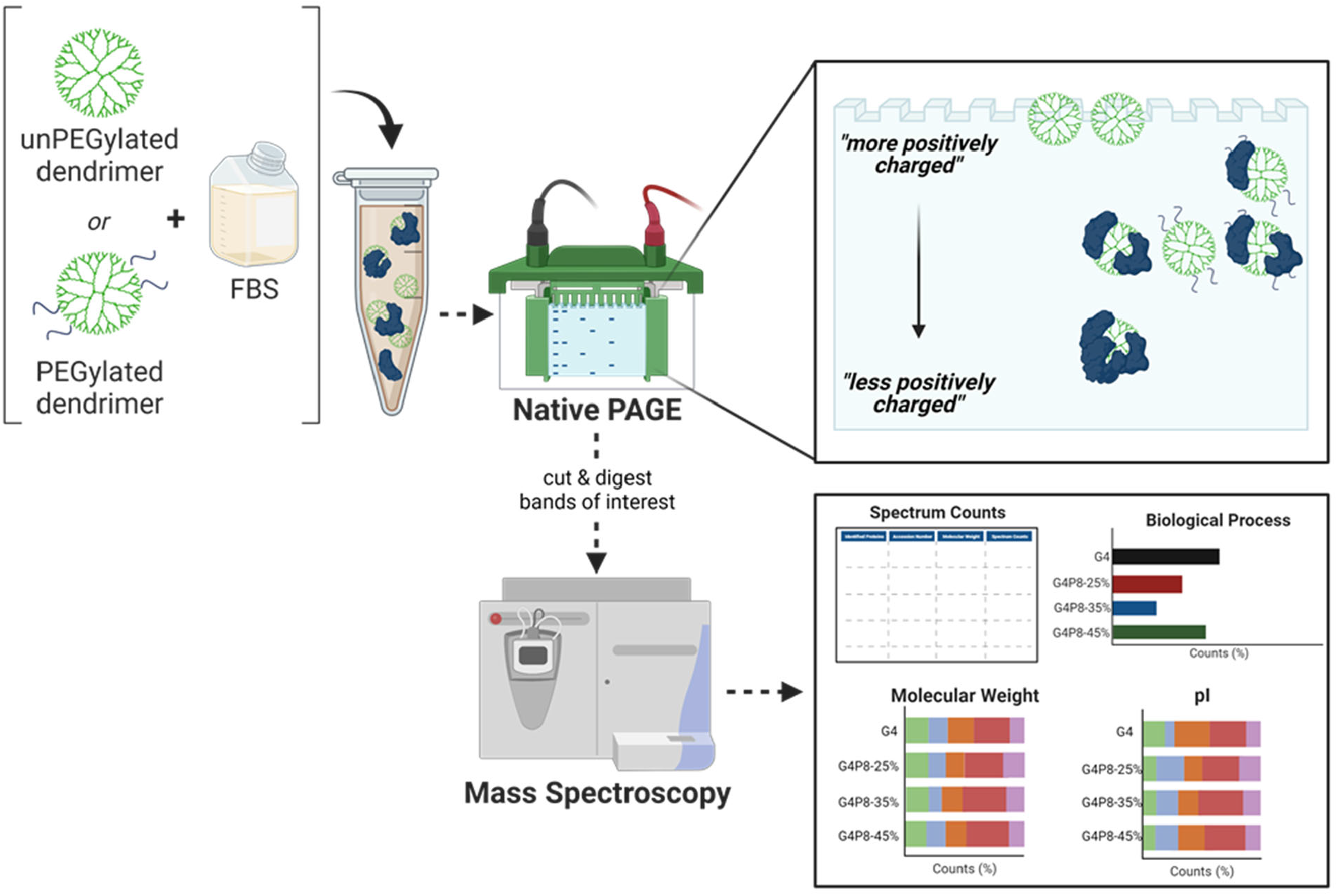

## Introduction

Polyamidoamine (PAMAM) dendrimers are small, hyperbranched macromolecules with repeating subunits of amides and a dense topological surface of cationic primary amines. PAMAM dendrimers are an emerging class of nanocarriers for drug delivery in a variety of therapeutic applications for brain^1,2^, cancer^3,4^, and gene therapy^3,5^, as well as detection diagnostics^6,7^. Our lab has identified PAMAM dendrimers as potential carriers that target cartilage to overcome delivery challenges for treating osteoarthritis; these carriers are small and positively charged, which facilitates diffusion through a dense negative charged matrix. Initial work is promising, as PAMAM dendrimers conjugated to insulin-like growth factor 1 (IGF-1) have enhanced therapeutic residence time in rat knee joints compared to free IGF-1^8^, and more recently, we characterized how the PEG corona on dendrimers interacts with ionizable surface groups during transport^9^. The next step is to understand how dendrimers interact with the biological environment by characterizing PAMAM dendrimer-protein interactions.

As nanocarriers are immersed in biological fluids, proteins, and other biomolecules adsorb to the nanocarrier’s surface forming a nanoparticle-protein complex. This complex, or protein corona, is considered the biological identity of the nanocarrier^10–13^. Protein coronas are determined by several factors including nanoparticle size^14–16^, material^17^, surface chemistry^15,16,18^, as well as biological environment^12,13,19^. This can potentially affect nanocarrier physiochemical properties, impacting their engineered targeting and therapeutic capabilities^20^. Biodistribution and pharmacokinetics of nanoparticles are affected by the protein corona, where adsorption of opsonin proteins to surfaces can trigger an immune response making nanoparticles recognizable by macrophages and vulnerable to phagocytosis^21–23^. Conversely, binding to dysopsonin proteins can reduce nanoparticle clearance, thereby improving their circulation time^11,23^. The protein corona can be perceived as a biological barrier to efficacious nanoparticle transport, targeting, and trafficking; identifying proteins adsorbed to nanoparticle surfaces can help overcome these challenges or provide knowledge that can be used to leverage their effects.

A challenging, but key step in identifying proteins adsorbed on nanoparticles is the complete separation of unbound proteins from proteins bound to nanoparticles. Most approaches for isolating nanoparticle-protein complexes for nanocarriers like polystyrene^14,24^, gold^16^, silicon dioxide^18^, iron oxide^22^, and liposomes^25^, include centrifuging and washing steps. After incubation, nanoparticles are centrifuged and the supernatant containing unbound protein is removed; the remaining pellet is washed to further remove proteins not bound to nanoparticles. This process is repeated until no protein is remaining in the supernatant. The remaining pellet, containing proteins bound to nanoparticles, is prepared for sodium dodecyl sulfate-polyacrylamide gel electrophoresis (SDS-PAGE) which uses denaturing conditions to separate proteins from nanoparticles to identify adsorbed proteins. Additionally, size and zeta potential can be assessed using dynamic light scattering^12,15,16^ to quantify changes in nanoparticle size and charge due to protein adsorption. Further, protein concentration can be quantified using a bicinchoninic acid (BCA) assay or ultraviolet-visible absorbance spectroscopy (UV-Vis)^15,16^. Other centrifugation-based methods including using a sucrose gradient^26^ or ultracentrifugation^27,28^ have also been applied to separate nanoparticle-protein complexes. A limitation is that centrifuging, in general, works well for larger, more dense nanoparticles. Smaller, less dense nanoparticles, like PAMAM dendrimers which are 4-6nm in size or roughly the size of the average protein, are not suited for traditional washing and centrifuging approaches as described above. To overcome these issues, previous work^29^ used agarose gels to separate PAMAM dendrimers from human plasma proteins which were cut and run on SDS-PAGE; individual bands were then isolated and run on mass spectrometry.

Inspired by this approach, we are using native PAGE to study dendrimer-protein interactions where proteins are separated based on their mass-to-charge-ratio under non-denaturing conditions, keeping dendrimer-protein complexes together while separating unbound proteins. The tris-glycine buffer used in native PAGE has a pH of ∼8.3 and does not heat up much during electrophoresis which helps prevent protein denaturing, preserving protein function and activity. We’re avoiding using SDS, as it has been shown that SDS and PEG complex^30^, which could interfere with discerning PEGylated dendrimer-protein interactions. Dendrimers are tagged with a fluorophore, which enables tracking of dendrimers in the gel and detecting where dendrimers migrate with protein. After electrophoresis, bands are isolated and run on mass spectrometry followed by proteome analysis (**Figure 1A**). Electrical current runs from negative to positive, top to bottom. It is expected that more positively charged dendrimers (ie: unmodified or unPEGylated dendrimers) are closer to the top (**Figure 1B**). As the dendrimer surface is conjugated with PEG (covalently bonded with shielding interactions) or coated with adsorbed proteins, the dendrimers charge is reduced or less positive and should migrate through the gel. We considered using isoelectric focusing (**Supplemental Figure 1**) which separates proteins based on isoelectric point, however, there was little to no protein shift for fetal bovine serum (FBS) that was incubated with dendrimers. Further, using this technique, proteins could be subjected to varying pH as they migrate through the gel, as the gel contains a pH gradient; this could alter the proteins interacting with the dendrimer. Our primary goal was to develop an electrophoresis-based protocol to investigate dendrimer-protein interactions. We hypothesize that dendrimers with higher PEG density would interact less with proteins compared to those with lower PEG density. Studies were optimized using FBS, and mass spectrometry results for dendrimer-FBS interactions were analyzed and compared to previously published work to verify our technique.

**Figure 1:**
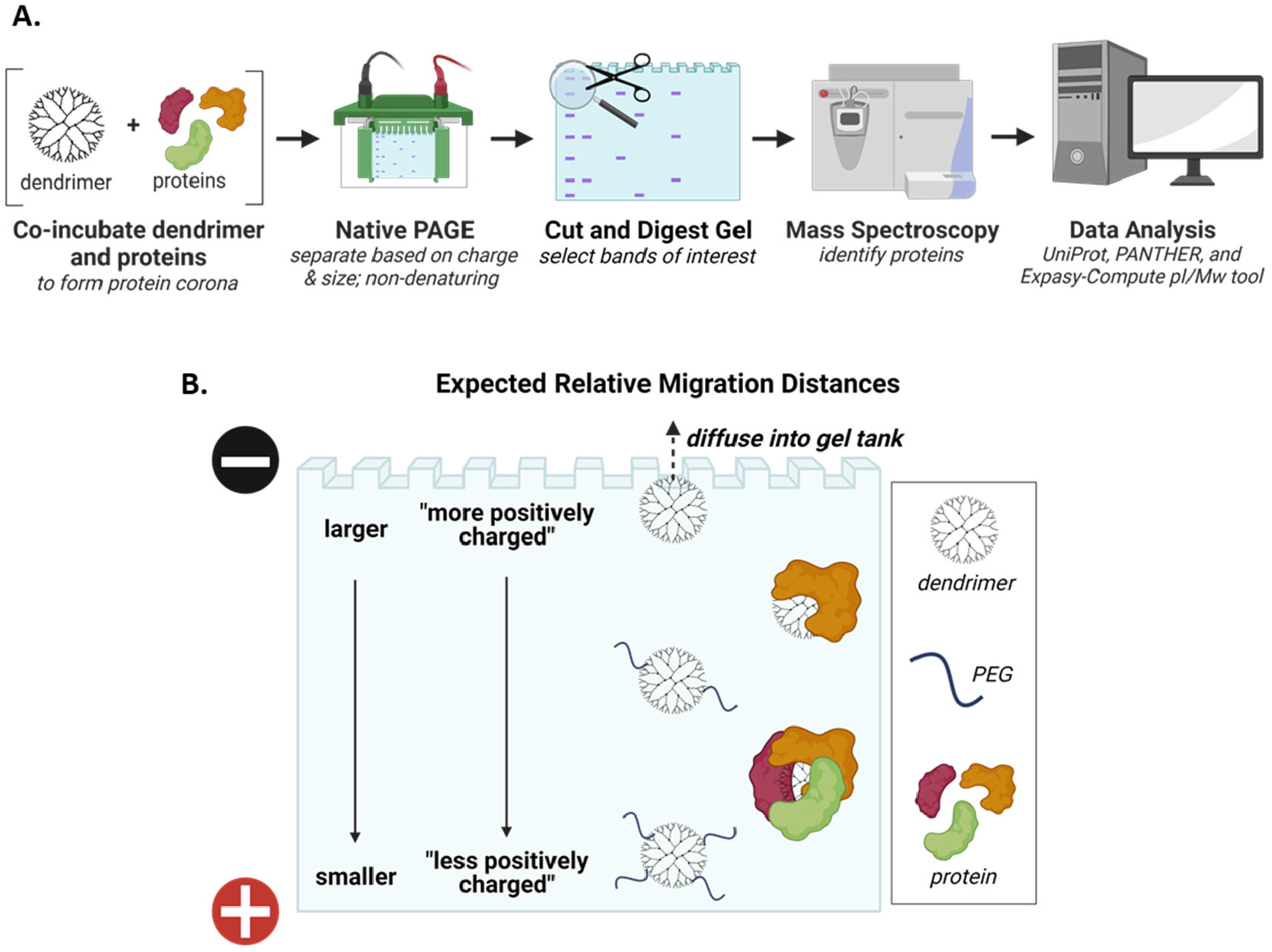
Native PAGE as a technique for separating PAMAM dendrimer-protein complexes. **(A)** Native PAGE separates proteins based on size and charge under non-denaturing conditions, which will keep dendrimer-protein interactions together. After electrophoretic separation, bands of interest are cut and digested before running mass spectrometry followed by analysis of identified proteins. **(B)** Electrical current runs top to bottom, negative to positive, which means more positively charged molecules are closer to the top of the gel while less positively charged molecules migrate towards the bottom. Pristine positively charged PAMAM dendrimers are expected to remain close to the top of the gel, and PEGylated dendrimers or protein coated dendrimers are expected to migrate further in the gel as dendrimers become less positive.

## Materials and Methods

### Materials

Generation 4 PAMAM dendrimers with ethylenediamine cores, were purchased from Dendritech through Sigma as solutions in methanol (G4 – 10 wt% in methanol). Methanol was removed via dry ice rotary evaporation from the dendrimer solution. After, the dendrimer solution was washed 4 times with DI H2O using ultracentrifugal filtration (Amicon Ultra 10k, 4 mL, Fischer Scientific). Purified dendrimer was lyophilized and reconstituted in MilliQ H2O and stored at 4°C. Methoxy PEG succinimidyl carboxymethyl ester (mPEG-SCM, MW=550 Da) was purchased from Creative PEGworks. Atto488 NHS ester was purchased from Sigma-Aldrich. Alexa Fluor™ 647 NHS ester (succinimidyl ester) was purchased from ThermoFisher Scientific. Anhydrous dimethyl sulfoxide (DMSO, HPLC grade, 99.9+%) was purchased from Alfa Aesar. Sodium bicarbonate (molecular biology grade) was purchased from Sigma. Deuterium oxide (99.9 atom % D, glass distilled) was purchased from Sigma-Aldrich. Fetal bovine serum (qualified) was purchased from Gibo (note: the same lot was used for all experiments). 1x phosphate-buffered saline (1x PBS, 150 mM salt, without calcium or magnesium) was purchased from Lonza. 4– 20% Mini-PROTEAN® TGX Stain-Free™ Gel-15 well, 4–20% Mini-PROTEAN® TGX™ Precast Protein -15 well, 2X Native Sample Buffer, 10X Tris/Glycine Buffer, and Coomassie Brilliant Blue R-250 Staining Solution were purchased from BioRad. Destain solution was prepared with 10% acetic acid (Sigma Aldrich) and 10% isopropanol (Sigma Aldrich) in MilliQ water.

### Dendrimer PEGylation

Generation 4 polyamidoamine (PAMAM) dendrimers were conjugated with PEG8 at various densities-25%, 35%, and 45% as previously described^8^. Briefly, 12.5 mM of PAMAM G4 dendrimer solution was made in a 10% v/v solution in 1M sodium bicarbonate solution and pH adjusted to 8 using hydrochloric acid. The amount of PEG needed to reach desired % PEGylation of end group was calculated and solubilized in anhydrous DMSO. Dendrimer solutions were diluted in 0.1M sodium bicarbonate ensuring DMSO was no more than 10% of the final volume. The PEG solution and dendrimer solution were combined, vortexed, and reacted covered while shaking at room temperature for 2 hours. Dendrimers were purified using ultracentrifugal flirtation (washed 4 times in DI water, 10k molecular weight cutoff filter) at 3500 x g. Following lyophilization, dendrimers were reconstituted to 1000μM in MilliQ water and stored at 4°C.

### NMR

PEGylation of PAMAM dendrimers was quantified using ^1^H NMR (Bruker AVANCE, 400 MHz or 500 MHz). Samples were lyophilized to remove water and reconstituted in D2O. Proton chemical shifts are reported in ppm (δ). ^1^H NMR (D_2_O) δ 2.39-2.44 (PAMAM, NCH_2_C**H**_**2**_CONH), δ 2.61-2.68 (PAMAM, CH_2_C**H**_**2**_N(CH_2_)_2_), δ 2.80-2.84 (PAMAM, NC**H**_**2**_CH_2_CONH), δ 3.00-3.15 (PAMAM, NH_2_C**H**_**2**_C**H**_**2**_NH), δ 3.36-3.39 (PEG, OC**H**_**3**_), δ 3.68-3.71 (PEG, OC**H**_**2**_C**H**_**2**_), δ 4.05-4.08 (PAMAM-PEG, NHCOC**H**_**2**_O). The integral ratio between PEG (OC**H**_**2**_C**H**_**2**_) and PAMAM methylene (NCH_2_C**H**_**2**_CONH) protons was used to calculate %PEG.

### Fluorescent Labeling of Dendrimers

1mM dendrimer was diluted in 0.1M solution bicarbonate at a 1:2 v/v ratio and vortexed. Atto 488 and Alexa Fluor 647 was prepared in anhydrous DMSO to a final concentration of 10mg/mL. Fluorophore was added to the dendrimer solution at a 1:1 molar ratio, vortexed, and reacted covered while shaking at room temperature for 2 hours. Fluorescently labeled dendrimers were purified using ultracentrifugal flirtation (washed 4 times in PBS, 10k molecular weight cutoff filter) at 3500 x g. Final solutions were stored at 4°C. To quantify fluorescent tags, the absorbance of fluorophores was measured using UV-Vis on a NanoDrop 1000 Spectrophotometer (ThermoFisher) with Atto 488 absorbance at 488nm (ε = 90,000 M-1cm-1) and Alexa Fluor 647 absorbance at 650nm (ε = 270,000 M-1cm-1).

### Protein Incubation & Native PAGE

Atto488 or Alexa Fluor 647 tagged PEGylated PAMAM dendrimers (G4-0%, G4P8-25, 35, and 45%) were incubated in fetal bovine serum (10% FBS or 100% FBS) for 1 hour at 37°C prior. 1X Native Sample Buffer was prepared using 2X Sample Buffer in a 1:1 v/v ratio in 1X PBS. Samples were prepped in 1X Native Buffer in a 1:1 v/v ratio. Prepped samples and run on Native PAGE on a 4–20% Mini-PROTEAN® TGX Stain-Free™ gel or 4–20% Mini-PROTEAN® TGX™ Precast Protein gel at 120V in 1X Tris-Glycine Buffer. Stain-Free gels were imaged on a ChemiDoc Imaging System (BioRad) with the blot 488 setting and protein gel Coomassie stain-free settings to image dendrimers and protein, respectively. Densitometry was done on Stain-Free gels using ImageJ to measure the fluorescent intensity of dendrimer and protein bands.”Regular” Precast Protein gels were imaged with the blot 647 setting to visualize dendrimers, and then stained with Coomassie Brilliant Blue R-250 Staining Solution for 1 hour and de-stained in de-staining solution overnight. Gels were then imaged with rapid auto exposure on the Coomassie Blue stain setting to image proteins before preparation for mass spectrometry.

### Protein Preparation for Mass Spectrometry

Protein isolation was performed by first cutting the top or shielded band (**Figure 3A**) of each gel lane, followed by reduction with 10mM dithiothreitol (Sigma) for 1 hour at 56°C and alkylation with 20mM iodoacetamide (Sigma) for 1 hour at 25°C in the dark. 12.5 ng/uL modified trypsin (Promega) was then used to digest the proteins in 100mM ammonium bicarbonate (pH 8.9) at 25°C overnight. To extract the proteins from the gel, each mixture was incubated in 50% acetonitrile/5% formic acid then 100mM ammonium bicarbonate, repeated two times, followed by incubation in 100% acetonitrile then 100mM ammonium bicarbonate, repeated two times. Finally, the proteins were dried using a vacuum centrifuge, desalted using Pierce Peptide Desalting Spin Columns (cat# 89852), and lyophilized.

Protein samples were separated by reverse phase HPLC using a PepMap RSLC C18 column (2μm tip, 75μm x 50cm, Thermo Fisher Scientific) before entering the mass spectrometer. All mass spectrometry analysis was performed using an Orbitrap Exploris 480 mass spectrometer (Thermo Fisher Scientific). Data-dependent analysis was performed on the samples with a resolution of 120,000, scan range of 375 to 1600 m/z, and maximum injection time of 25 seconds. The initial scan was followed by MS/MS with a NCE of 28, dynamic exclusion of 20 seconds, and resolution of 30,000.

The HPLC mobile phases were 0.1% formic acid in water (A) and 0.1% formic acid in acetonitrile (B). The gradient conditions were as follows: 1% B (0-10 min at 300 nL/min) 1% B (10-15 min, 300 nL/min to 200 nL/min) 1-7% B (15-20 min, 200 nL/min), 7-25% B (20-54.8 min, 200 nL/min), 25-36 B (54.8-65 min, 200 nL/min), 36-80% B (65-65.5 min, 200 nL/min), 80% B (65.5-70 min, 200 nL/min), 80-1% B (70-70.1 min, 200 nL/min), and 1% B (70.1-90 min, 200 nL/min).

### Proteomic Analysis

Protein identification was performed on the raw data files from MS using Sequest HT found in Proteome Discoverer (Thermo Fisher Scientific). The following criteria were used in the search against the Bovine protein database: 10 ppm peptide mass tolerance, 0.02 Da fragment mass tolerance, and 2 missed cleavages. Carbamidomethylation on cysteines was set to fixed modification. Methionine oxidation, methionine loss at the N-terminus, acetylation of the N-terminus, and Met-loss plus acetylation of the N-terminus were set to variable modifications. The resulting data was analyzed using Scaffold 5 software found in Proteome Discoverer (Thermo Fisher Scientific); total spectrum count was reported with accession number and molecular weight. The percent total of spectrum counts was calculated by dividing the total spectrum count of an individual protein by the sum total spectrum count for each experimental group. If the same protein was identified in the top and shielded band, the total spectrum count for both bands were summed and then divided by the total spectrum count of the individual protein. UniProt and PantherDB was used to identify the biological functions of identified proteins. ExPASy-Compute pI/Mw tool was used to sort identified proteins by isoelectric point (pI).

## Results & Discussion

### Identifying electrophoretic migration of PAMAM dendrimers in native PAGE

Migration distances of Atto 488 tagged dendrimers in the absence of proteins were assessed first. Dendrimers are tagged with dye in a 1:1 mole ratio (1 Atto 488 dye per dendrimer); surface charge properties from dendrimers come from the primary amines and dye labeling converts one charged amine to an amide bond-a 1.6% decrease in positive charge which should not have a significant effect on the surface charge properties^9^. Previous work studied Generation 4 PAMAM dendrimers conjugated with PEG8 (550 Da) at varying percentages-25, 35, and 45% (denoted as G4-0%, G4P8-25$, G4P8-35%, and G4P8-45%) which were promising nanocarriers for targeted cartilage delivery^8^. Various concentrations (0.6, 1.25, and 2.5 μM) of dendrimers were run on a stain-free gel that contains a proprietary trihalo compound that covalently binds to tryptophan residues after photoactivation (**Figure 2**). Dendrimers and proteins were visualized in the 488 channel (**Figure 2**, green box) and Coomassie-700nm (**Figure 2**, blue box) channels, respectively. UnPEGylated dendrimers stay at the top of the gel (black arrow, top band), however, PEGylated dendrimers have a second lower band (white arrow). We hypothesize the shift in migration of PEGylated dendrimers is due to charge differences between unPEGylated and PEGylated dendrimers caused by a combination of covalent amide bond formation and non-covalent PEG shielding which results in a less positive charge. Accordingly, we refer to this second band as the”shielded band”. This observation suggests that in native PAGE, the difference between the mass-to-charge ratio is negligible between these PEGylated dendrimers. As dendrimer concentration increases, the fluorescence for the top and shielded bands increases. A faint top band was also observed for PEGylated dendrimers; we hypothesize this is in part due to the conjugation efficiency of the dendrimer with PEG. Fluorescence was observed within the well (**Figure 2**, orange box) for unPEGylated dendrimers at higher concentrations. This likely occurs due to positively charged dendrimers migrating toward the top or negative end of the gel, as previously observed in agarose gels^29^. Dendrimers were not expected to be detected in the Coomassie-700nm channel, however, there is a slight spectrum overlap between 488nm and 700nm. Likely as a result, high concentrations of unPEGylated dendrimer were detected in the 700nm channel. Overall, there was minimal overlap between dendrimer and protein signals (**Figure 2**, grey box). For the FBS only control, with no dendrimer, protein was not detected in the 488nm channel, confirming there was no overlap of protein and dendrimer signal. Collectively, this enables us to distinguish between dendrimers and proteins after co-incubation and assess interactions between them.

**Figure 2:**
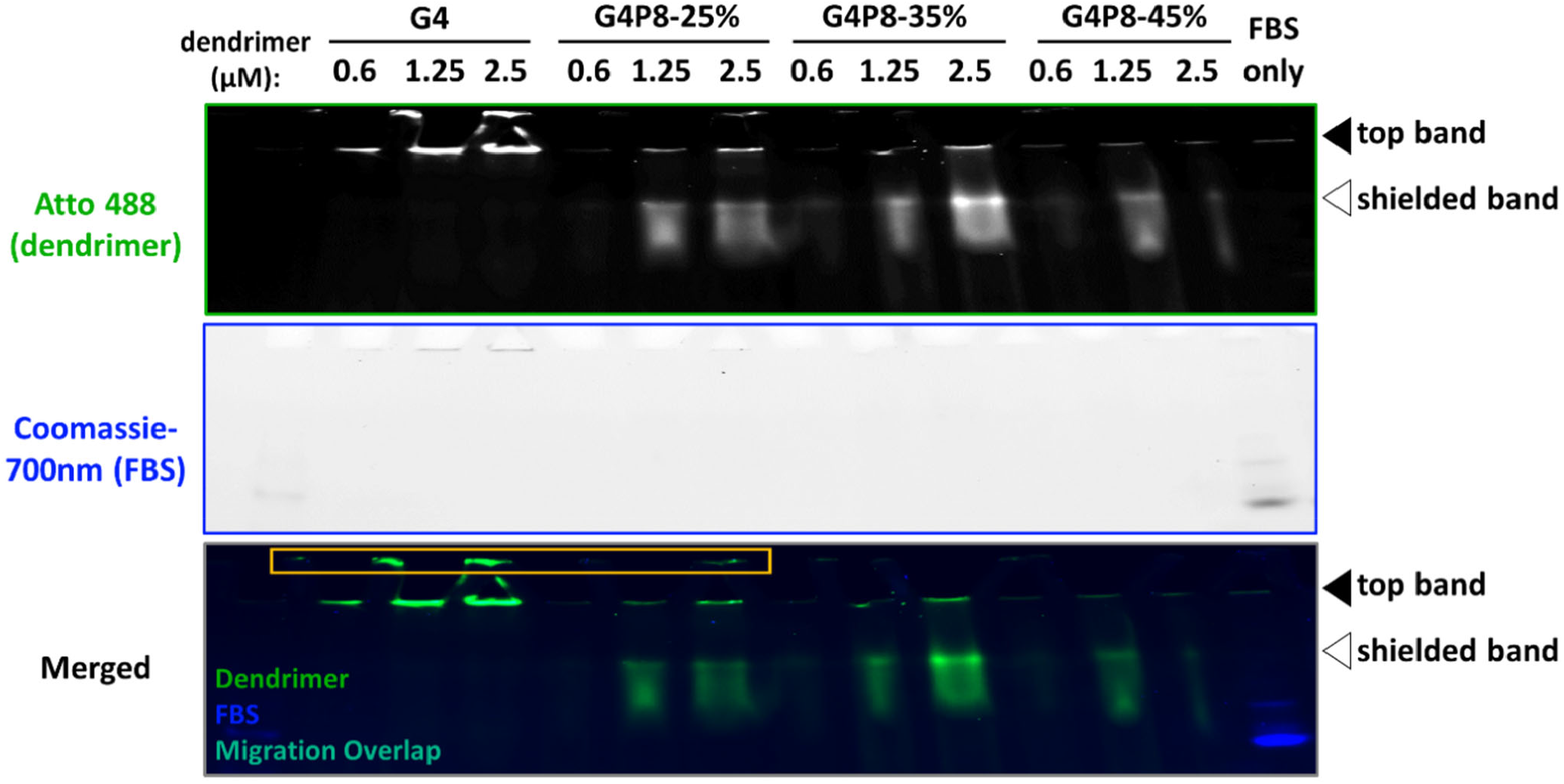
Identifying electrophoretic migration of PAMAM dendrimers in Native PAGE. Various concentrations of PEGylated PAMAM Generation 4 dendrimers tagged with Atto 488, as well as protein only control (10% FBS), were run on a”stain-free” Native PAGE gel. Dendrimers and proteins are imaged simultaneously, where dendrimers (green box) and proteins (blue box) are identified in the 488nm and Coomassie-700nm channels, respectively. As dendrimer concentration increases, there is an increase in fluorescence as observed in the Atto 488 channel. UnPEGylated and PEGylated dendrimers are at the top of the gel (black arrow, top band), however, PEGylated dendrimers have a lower second band (white arrow, shielded band). As more PEG is added, there is little to no difference in the migration distance of the shielded band. In the Coomassie-700nm channel, protein only controls were detected. Fluorescence was observed within the well (orange box) for unPEGylated dendrimers at higher concentrations. This demonstrates how a”stain-free” gel can be used to discern between fluorescently tagged dendrimers and protein.

### Determining interactions between PAMAM dendrimers and proteins

Selection of biological fluid is important for protein corona studies, as varying protein content and concentrations affect which proteins comprise the corona^24^ Although serum lacks coagulation factors found in plasma, FBS was used as a test biological fluid as it is commonly used to supplement cell media, contains similar proteins found in human plasma, and can be used to compare our results to published work^29^. **Supplemental Table 4** lists the protein content of FBS used in these studies which we compared to previously published tables^29^. Dendrimers (0.6 μM) are incubated in FBS (10% and 100%) for 1 hour at 37°C to form a protein corona. Samples were run on native PAGE (**Figure 3A**). Dendrimers (**Figure 3A**, green box) and proteins (**Figure 3A**, blue box) are identified in the 488nm and Coomassie-700nm channels, respectively. Densitometry was used to quantify fluorescence in the top band and shielded band. For the Atto 488/dendrimer channel (**Figures 3B** and **3C**) data were normalized to max fluorescence intensity and for the Coomassie-700nm/FBS channel (**Figures 3C** and **3D**) data were normalized to 100% FBS control fluorescence intensity. The ratio of shielded band to top band for the 488nm (dendrimer) and Coomassie 700nm (protein) channels were reported to quantify relative distribution of proteins and dendrimer migration between the two bands. It was determined that the relative ratio of dendrimers and proteins are consistent across unPEGyalted and PEGylated dendrimers (**Supplement Figure 3**).

**Figure 3:**
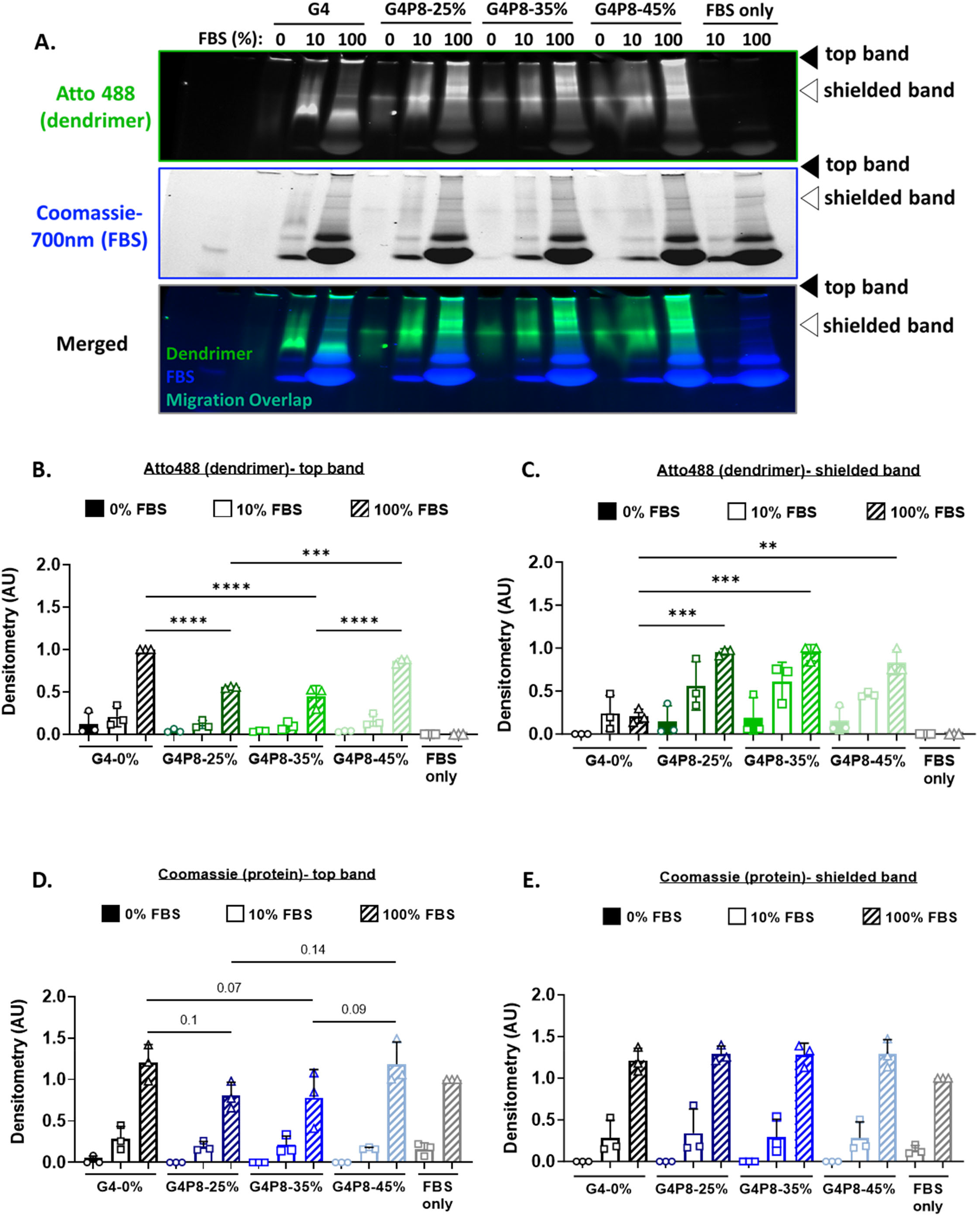
Electrophoretic migration of PAMAM dendrimers was measured to determine interactions between dendrimers and FBS proteins. 0%, 25%, 35%, and 45% PEGylated (PEG-8) PAMAM Generation 4 dendrimers tagged with Atto 488 were incubated in fetal bovine serum (FBS, 3.6 or 36 mg/mL) for 1 hour at 37°C prior to separation using Native PAGE. **(A)** Dendrimers are identified at 488nm and proteins are identified at Coomassie-700nm. Arrows indicate the previously identified migration distances of pristine dendrimers (i.e.: no protein). Across all groups, as the amount of FBS increases, there is an increase in fluorescence in top band (black arrow), which correlates to an increase in the amount of dendrimer. For unPEGylated dendrimers, when FBS is added additional bands appear lower in the gel. For PEGylated dendrimer incubated with FBS, bands appear between the top band and second band. In the 488nm channel and Coomassie-700nm channel, we observe the dendrimers and proteins (blue box) migrate similar distances suggesting there are interactions between the two. Fluorescence in the **(B**,**C)** Atto 488/dendrimer channel (data normalized to max intensity) and **(D**,**E)** Coomassie-700nm/FBS channel (data normalized to 100% FBS control) was quantified with densitometry. Densitometry with 3 technical replicates; all data reported as mean with standard deviation. **p<0.01, ***p<0.001, ****p>0.0001

When assessing the Atto 488 channel (dendrimer), in general, as more FBS is added, the amount of dendrimer fluorescence increases. Specifically, in the top band, dendrimer fluorescence significantly increases as FBS increases from 0% to 100%, as well as 10% to 100% across all groups, suggesting more dendrimers are present likely due to increased interactions with proteins. Although the same concentration of dendrimer is added to FBS for all experimental groups, we hypothesize that free dendrimer is so positively charged that it does not migrate in the gel but rather diffuses away from the gel into the gel tank when the field is applied. FBS shields charge when it binds to dendrimers, leading to a decrease in this migration. In the top band, for dendrimers incubated with 100% FBS, there was significantly more fluorescence for unPEGylated dendrimers compared to 25% and 35% PEGylated dendrimers. As such, we unexpectedly observed that fluorescence was significantly increased for 45% PEGylated dendrimers, compared to 25% and 35% PEGylated dendrimers. In the shielded band, there is a significant increase in fluorescence for PEGylated compared to unPEGylated dendrimers. This increase in fluorescence is somewhat to be expected as a shielded band was not present for unPEGylated dendrimers without protein. However, for unPEGylated dendrimers incubated in FBS, a band appears further down in the gel that is below the shielded band. Bands below the shielded band were not observed in PEGylated formulations. For PEGylated dendrimers, additional bands appear between the top and shielded charge bands. These additional bands have a higher positive charge relative to the shielded bands and may represent dendrimers with differing degrees of end-functionalization.

Taken together, we hypothesize that as proteins are interacting with dendrimers, PEG repels proteins helping preserve the positive charge of dendrimers. These observations suggest that the shielding of proteins by PEG prevent dendrimers from becoming less positively charged compared to unPEGylated dendrimers that engage more negatively charged serum proteins. This result also demonstrates proteins (which can present negative charge) can have more of an effect in altering dendrimer net charge compared to neutral PEG, which in turn has implications for how a protein corona could affect electrostatic interactions of dendrimers with a negatively charged tissue matrix, such as cartilage. Collectively, this is further evidence of dendrimer-protein interactions that lead to changes in dendrimer charge, where more proteins reduce the dendrimer’s positive charge resulting in migration towards the bottom of the gel. For PEGylated dendrimers, more smearing between the top and shielded bands was observed for dendrimers incubated with FBS. We hypothesize smearing occurs as dendrimer migration properties change due to dendrimer-protein interactions, leading to a range of differently charged nanomaterials within the sample^31^. This smearing, in combination with dendrimers and proteins migrating together, is indicative of potential interactions.

When comparing banding patterns in the Coomassie-700nm channel (protein) between FBS control and experimental groups, there are no notable differences, however, after analyzing densitometry we see differences in relative protein amounts. For all dendrimer formulations, as more FBS is added, the amount of protein in the top band significantly increases as FBS increases from 0% to 100% and 10% to 100%. The same trend was observed in the shielded band. However, in the shielded band, all dendrimer formulations had more protein compared to the FBS controls; although this was not statistically significant, it could suggest more proteins migrating to that site likely due to dendrimer-protein interactions or dendrimers enriching proteins from FBS. In the top band for dendrimers incubated with 100% FBS, for 25% and 35% PEGylated dendrimers, less protein is present compared to unPEGylated dendrimers. This was to be expected, as PEG has been used to reduce protein adsorption on biomaterial surfaces^13,32^. Compared to the FBS only controls, more protein was detected with unPEGylated dendrimers, though it was not a statistically significant increase. This suggests additional proteins, potentially ones that could not migrate through the gel, are interacting with the dendrimer. Unexpectedly, 45% PEGylated dendrimers had more protein compared to 25% and 35% PEGylated dendrimers. We hypothesize this could be due to the accumulation of a higher molecular weight proteins which is later corroborated by mass spectroscopy where we observed an increase in spectral counts of”top hits” of proteins apolipoprotein B which has a molecular weight of 516kDa (**Supplemental Figure 4A**). Notably, there is also an increase in spectral counts for other proteins (complement C3, C4a anaphylatoxin, plasminogen, and alpha 2 macroglobulin) for 45% PEGylated dendrimers relative to 25% and 35% PEGylated dendrimers.

### Mass spectroscopy analysis

For dendrimers incubated in 10% FBS, the top and shielded bands (**Figure 3**, black arrow and white arrow, respectively) were cut and digested, then run on mass spectrometry to identify proteins (note for mass spectroscopy”regular” Coomassie-stained gels are used, **Supplemental Figure 2**). These bands were selected as both appeared in unPEGylated and/or PEGylated dendrimers without protein, indicating that they represent dendrimer migration. Data was analyzed using the proteome software Scaffold 5 and total spectrum count was reported with accession number and molecular weight. The percentage of spectrum counts was quantified for each identified protein. Top hits are reported in **Table 1** with the percentage of total spectrum counts reported in a graph in **Supplemental Figure 4A**. A complete list of identified proteins is in **Supplemental Table 2**. Previous studies incubated dendrimers with human plasma, however, dendrimer-protein isolation was done using different methods^29^. Briefly, dendrimers were incubated in human plasma and then run on an agarose gel to separate dendrimer bound proteins. Samples were cut from the agarose gel and run on SDS-PAGE, then individual bands were isolated and run on mass spectrometry to identify proteins. Given that there are similar proteins in FBS and plasma, to further evaluate our Native PAGE approach, we compared our identified proteins to the”significant hits” identified in previous studies. In total, we matched 6 out of 8 proteins: Cluster of Apolipoprotein B, Complement C3, Cluster of C4b-binding protein alpha chain, vitronectin, ceruloplasmin, and prothrombin, with 2 being among our top hits. It should be noted that apolipoprotein B was only found in the top band, and ceruloplasmin and prothrombin were only identified in the shielded band-this is likely due to size differences (516kDa, 121 kDa, and 71kDa, respectively). Ceruloplasmin is identified in the stock FBS (0.045%, **Supplemental Table 4**), however, it was not identified in the FBS control after electrophoresis. Interestingly, ceruloplasmin was associated with the unPEGylated dendrimer (0.556%), 25% PEG (0.268%), and 35% PEG (0.054%). In other words, in the absence of dendrimers, ceruloplasmin in FBS does not migrate into the gel likely due to the negative charge of ceruloplasmin. Further, as PEG is added there is a decrease in the amount of ceruloplasmin associated with dendrimer demonstrating resistance to protein adsorption due to PEG shielding. There were other proteins not identified in FBS control bands, but were found associated with dendrimers; some of these include CD109 molecule, collagen-alpha chain, and tetranectin (**Supplemental Figure 5**). This demonstrates the ability of our method to capture how dendrimers enrich or deplete proteins regardless of the amount or concentration in stock FBS or proteins that migrated on electrophoresis in the FBS only control.

**Table 1:** Top hits from mass spectrometry on PAMAM dendrimer-FBS protein complexes in”top band” and”shielded band” for dendrimers incubated in 10% FBS. The top or shielded band (black arrow and white arrow, respectively, Figure 3A) from Native PAGE gels for dendrimers incubated in 10% FBS were excised, digested, and run on mass spectroscopy. Top proteins identified in all experimental conditions are reported in the table. To evaluate our approach, we compared our top hits proteins to”significant hits” that were previously published for incubated dendrimers in human plasma and matched 6 out of 8 proteins, where 2 (indicated by an asterisk) were found in our top hits.

| Identified Proteins | Accession Number | Molecular Weight | Theoretical pI | Biological Process |
| --- | --- | --- | --- | --- |
| Complement C3* | Q2UVX4 | 187 kDa | 6.37 | Complement |
| Apolipoprotein B* | E1BNR0 | 516 kDa | 6.24 | Lipoprotein |
| Cluster of Beta-1 metal-binding globulin | G3X6N3 [3] | 78 kDa | 6.63 | Transport/Carrier/Binding |
| Cluster of C4a anaphylatoxin | F1MVK1 [3] | 192 kDa | 7.26 | Complement |
| Cluster of Alpha-2-macroglobulin | Q7SIH1 [2] | 168 kDa | 5.68 | Protease Inhibitor |
| Cluster of Albumin | A0A140T897 [2] | 69 kDa | 5.77 | Transport/Carrier/Binding |
| Cluster of Plasminogen | P06868 [2] | 91 kDa | 7.39 | Coagulation |

Based on the percentage of total spectrum counts, the proteins most associated with dendrimers were Apolipoprotein B and complement C3. These two proteins have been previously identified as the most abundant proteins in the corona of silica nanoparticles with surfaces functionalized with primary amines^10^. Complement proteins C4a and C5a are enzymatically cleaved from full-length C4 and C3/C5, respectively, and are known to induce inflammatory responses^33^; these were also found to be associated with dendrimers where C4a was identified as a top hit. There is an increase in the percentage of total spectrum counts for complement C4a as the percent PEG increased on dendrimers, and a similar trend was observed for complement C3 as well. This is unexpected as protein adsorption and conformation on nanoparticles is dependent on PEG chain length and density^32,34^. However, it has been reported that PEG can generate complement activation proteins^35^; specifically, iron oxide PEG nanoparticles induced increment of C3a and C5a anaphylatoxins^36^. More recently, there is evidence that complement C3 proteins can bind to existing proteins or protein corona^37^ rather than the surface of the nanoparticle. This further suggests the dynamic nature of protein-nanoparticle or rather protein-protein interactions on nanoparticles. Although the incubation time of one hour could be considered a longer incubation time, it has been shown that protein corona forms in a relatively stable manner during a one-hour period^38^; thus, under these conditions, we are capturing the hard corona^12,17,22^ and the increase in complement C3 could be due to complement C3 binding to underlying proteins on the dendrimer surface

### PAMAM dendrimers mostly interact with complement proteins in FBS

Proteins that were identified as part of dendrimer-protein interactions were classified by biological function (**Figure 4**). Dendrimers mostly interact with complement proteins with transport proteins being the second highest, aligning with previously published work. For complement proteins, as percent PEGylation increases, there is an increase in the percentage of total spectrum counts which is to be expected based on our brief analysis of complement proteins above. In addition to biological function, proteins were sorted by molecular weight (**Supplemental Figure 4B**); most proteins (>50%) were greater than 80kDa. Given the size of the dendrimers (4.5nm) and how they tend to interact with larger molecular weight proteins, it is hypothesized that dendrimers act as cargo on proteins^39^, an interesting property to consider in future studies of small nanocarriers. Lastly, proteins were analyzed by isoelectric point (**Supplemental Figure 4C**); a majority of proteins (∼85%) are between pI 5 and 7. There is a notable shift in the distribution of pIs falling between 5-5.9 and 6-6.9, and an increase in pI > 7 for dendrimers incubated in FBS.

**Figure 4:**
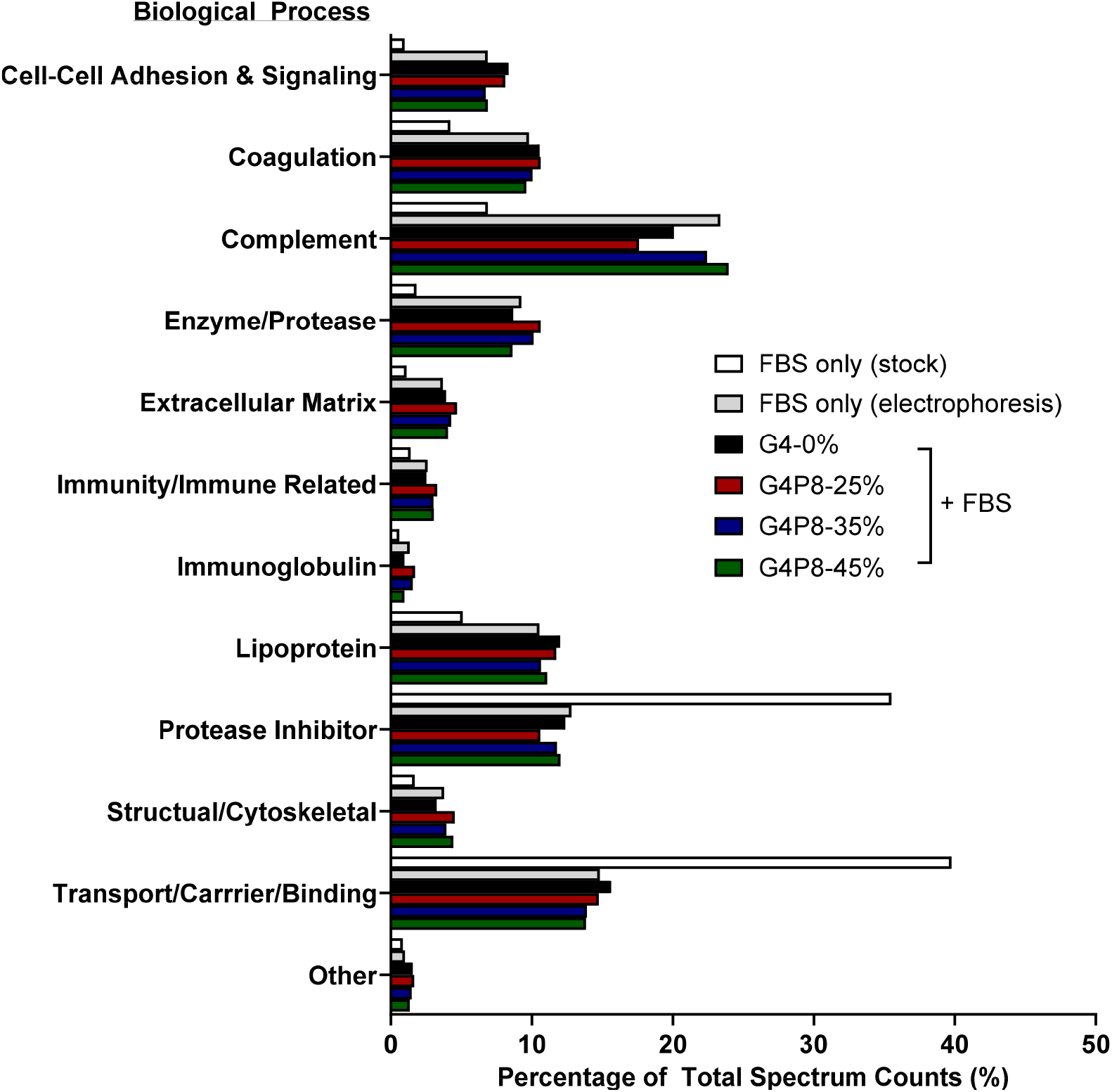
PAMAM dendrimers mostly interact with complement proteins in FBS. The top band and shielded band (black arrow and white arrow, respectively Figure 3A) from Native PAGE gels for dendrimers incubated in 10% FBS were excised, digested, and run on mass spectroscopy. All identified proteins were sorted by biological process and reported as the percentage of total spectrum counts.

Something to take under consideration for this technique is that it requires selecting which bands to run on mass spectrometry, and other methods using SDS-PAGE to study the protein corona used this approach as well^14,28,29^. Here we decided to run the top and shielded bands for 10% FBS because both bands were identified in dendrimers not incubated with proteins. We also analyzed the top band for 100% FBS on mass spectroscopy. To do this, regular gels stained with Coomassie were used for mass spectroscopy; the shielded band was not run for these conditions as it is difficult to clearly identify and isolate bands due to the high protein concentrations (**Supplemental Figure 2**). For this band, similar trends were observed when compared to 10% FBS (**Supplemental Figure 6, Supplemental Tables 1** and **3**)-dendrimers mostly associated with complement proteins. However, there was a greater number of lipoproteins associated, likely due to the amount of apolipoprotein B in the top band of 100% FBS compared to 10% FBS with percentage of total spectrum counts of ∼15% and ∼8%, respectively.

Polyacrylamide electrophoresis has been used to confirm successful protein-polymer conjugation where an increase in molecular size is expected to be observed^40^. However, challenges with the technique can arise as it has been reported that interactions between PEG and sodium dodecyl sulfate (SDS) occur^30^. Native PAGE has been used previously to study protein-protein interactions^41,42^. PEG is a neutral polymer; thus it is expected to migrate on a Native PAGE gel; however, it will migrate on SDS-PAGE and complex with SDS^30^ This phenomenon justifies avoiding running PEGylated nanocarriers on SDS-PAGE as a method to study nanocarrier-protein interactions. When designing nanocarriers, attaching PEG to reduce cytotoxicity is a commonly used approach^43^. Further, PEG has shielding properties that help reduce protein adsorption^13,32^. As such, it would be anticipated that PEGylated dendrimers would have a lower quantity of bound proteins, but as our results show, there are still interactions with proteins, although the amount of protein is reduced relative to unPEGylated dendrimers. As there is more interest in studying the protein corona or protein adsorption to nanoparticles, tools must be compatible with nanoparticle surface chemistry to ensure accurate characterization. Our approach offers a way for investigators to study PEGylated nanocarriers without interference from SDS.

## Conclusions

The phenomenon of proteins adsorbing to nanoparticles is not new; some of the first studies describing protein adsorption to nanoparticles date back to the 1960s, with the term protein corona being coined in 2007^13^. Yet, the field still needs methods to study protein-nanoparticle interactions, and to improve our ability to leverage protein coronas for diagnostic and therapeutic applications. New techniques are needed, especially to overcome the challenge of studying these complexes on smaller nanocarriers (<10nm), like PAMAM dendrimers. Here, we developed an approach using Native PAGE that capitalizes on principles of nondenaturing separation to study dendrimer-protein interactions. Our protocol offers several advantages including being more streamlined, requiring one round of electrophoresis followed by mass spectrometry. Dendrimers and proteins can be detected and imaged simultaneously using stain-free gel technology, and bands in stain-free gels have a good resolution which can be quantified with densitometry. Using mass spectrometry, we identified similar proteins compared to previously published data, which verifies our approach can be used to isolate and identify proteins bound to dendrimer. This technique expands our range and ability for studying the protein corona on dendrimers; we think it has the potential to be applied to other small nanoparticles such as quantum dots or aptamers, so the field can gain more understanding about the protein corona of these types of nanoparticles.

## Supporting information

Supplemental Information_Electrophoresis-Based Approach for Characterizing Dendrimer-Protein Interactions

## Acknowledgments

The authors would like to thank and acknowledge Rick Schiavoni and the Biopolymers & Proteomics in the Swanson Biotechnology Center at the Koch Institute for Integrative Cancer Research at MIT for preparing and running samples on mass spectroscopy, as well as providing the protocol for the Materials and Methods section. We also thank and acknowledge Justin Kaskow for assisting with setting up IEF and the Love Lab at the Koch Institute for Integrative Cancer Research at MIT for loaning us their gel electrophoresis power supply to run IEF. Also, special thanks to Brandon Johnston, Joon Ho Park, and Justin Kaskow for critical review of this manuscript.

## Author Information

Corresponding Author

Paula T. Hammond

Department of Chemical Engineering, Massachusetts Institute of Technology, 77

Massachusetts Ave, Cambridge MA, 02139

Koch Institute for Integrative Cancer Research, 500 Main St, Cambridge MA, 02142

Authors

Simone A. Douglas-Green

Department of Chemical Engineering, Massachusetts Institute of Technology, 77

Massachusetts Ave, Cambridge MA, 02139

Koch Institute for Integrative Cancer Research, 500 Main St, Cambridge MA, 02142

Juan A. Aleman

Department of Chemical Engineering, Massachusetts Institute of Technology, 77

Massachusetts Ave, Cambridge MA, 02139

## Author Contributions

Conceptualization and Methodology: SDG. Investigation and Validation: SDG and JAA. Formal Analysis: SDG. Supervision: PTH. Writing-original draft: SDG and JAA. Writing-review and editing: SDG, JAA, and PTH. Project funding acquisition: SDG and PTH.

