## Supplemental Information_Electrophoresis-Based Approach for Characterizing Dendrimer-Protein Interactions for "Electrophoresis-Based Approach for Characterizing Dendrimer-Protein Interactions: A Proof-of-Concept Study"

### Supporting Information

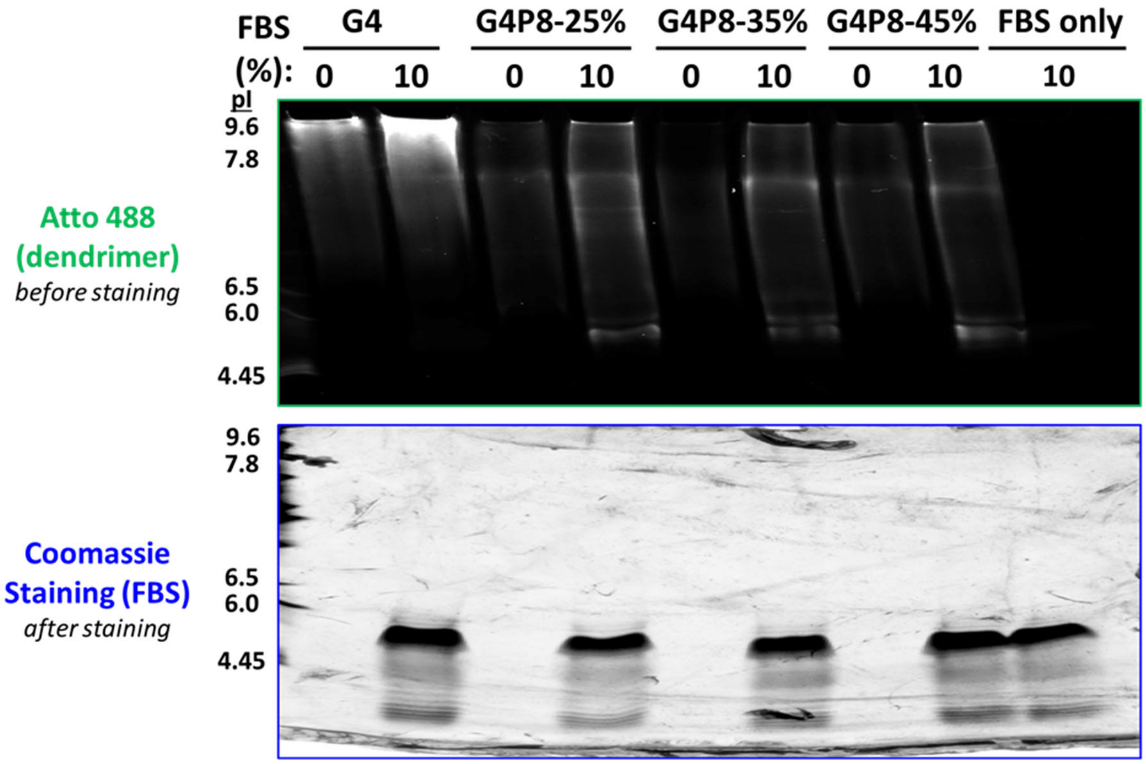

**Supplemental Figure 1: Isoelectric focusing gel on dendrimers co-incubated with FBS.** Isoelectric focusing (IEF) gels separate proteins by isoelectric point (pI) under Native or non-denaturing conditions to determine whether a protein is positive or negative at physiological conditions. IEF was run to assess if there was a shift in migration of FBS after incubation which would indicate dendrimer-FBS interactions. Dendrimers (G4-0%, G4P8-25, 35, and 45% PEG) were incubated in FBS (10%, 3.6mg/mL) for 1 hour at 37°C, and samples were prepped and run on an IEF gel. (Top) Dendrimer fluorescence increased after incubation with FBS. Without FBS, a top band was observed for unPEGylated and PEGylated dendrimers, and 2nd band was observed in the PEGylated formulations. This migration pattern is similar to what was observed in Native PAGE (Figures 2 and 3A). However, for PEGylated dendrimers incubated with FBS, additional lower bands appeared which align with where FBS was detected after staining. (Bottom) There was little to no shift for FBS proteins incubated with PEGylated dendrimers, however, there was a slight upward shift for unPEGylated dendrimers.

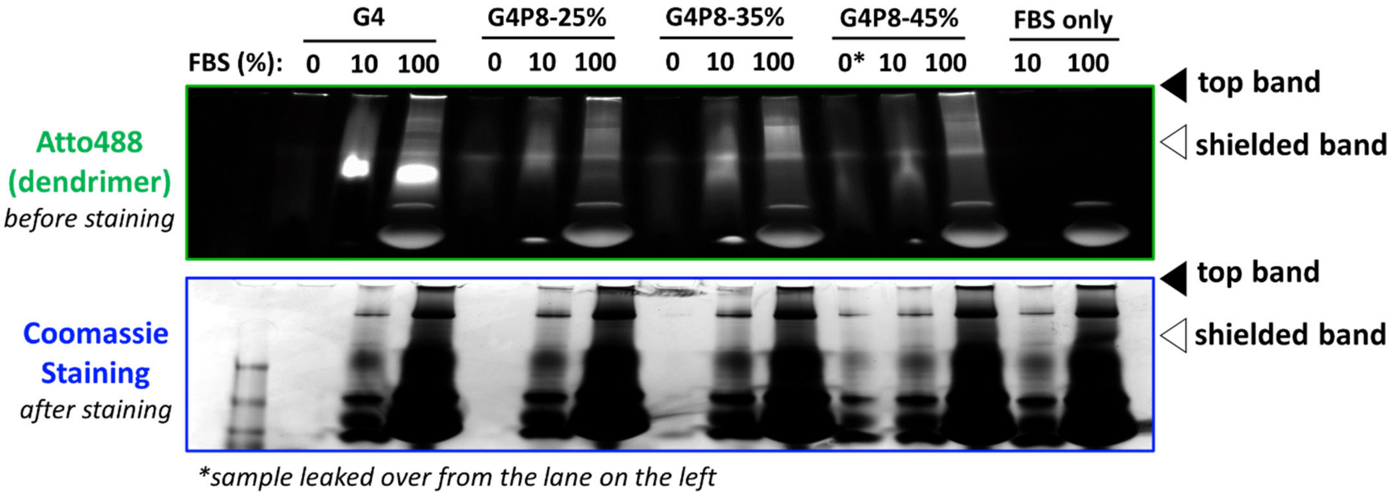

**Supplemental Figure 2: PAMAM dendrimers co-incubated with FBS are run on a regular Native PAGE gel for mass spectroscopy.** Gels are imaged at 647 to detect dendrimers before staining, as Coomassie dye will interfere with the fluorescence imaging. After gels are stained and destained, bands of interest are isolated, digested, and run on mass spectroscopy.

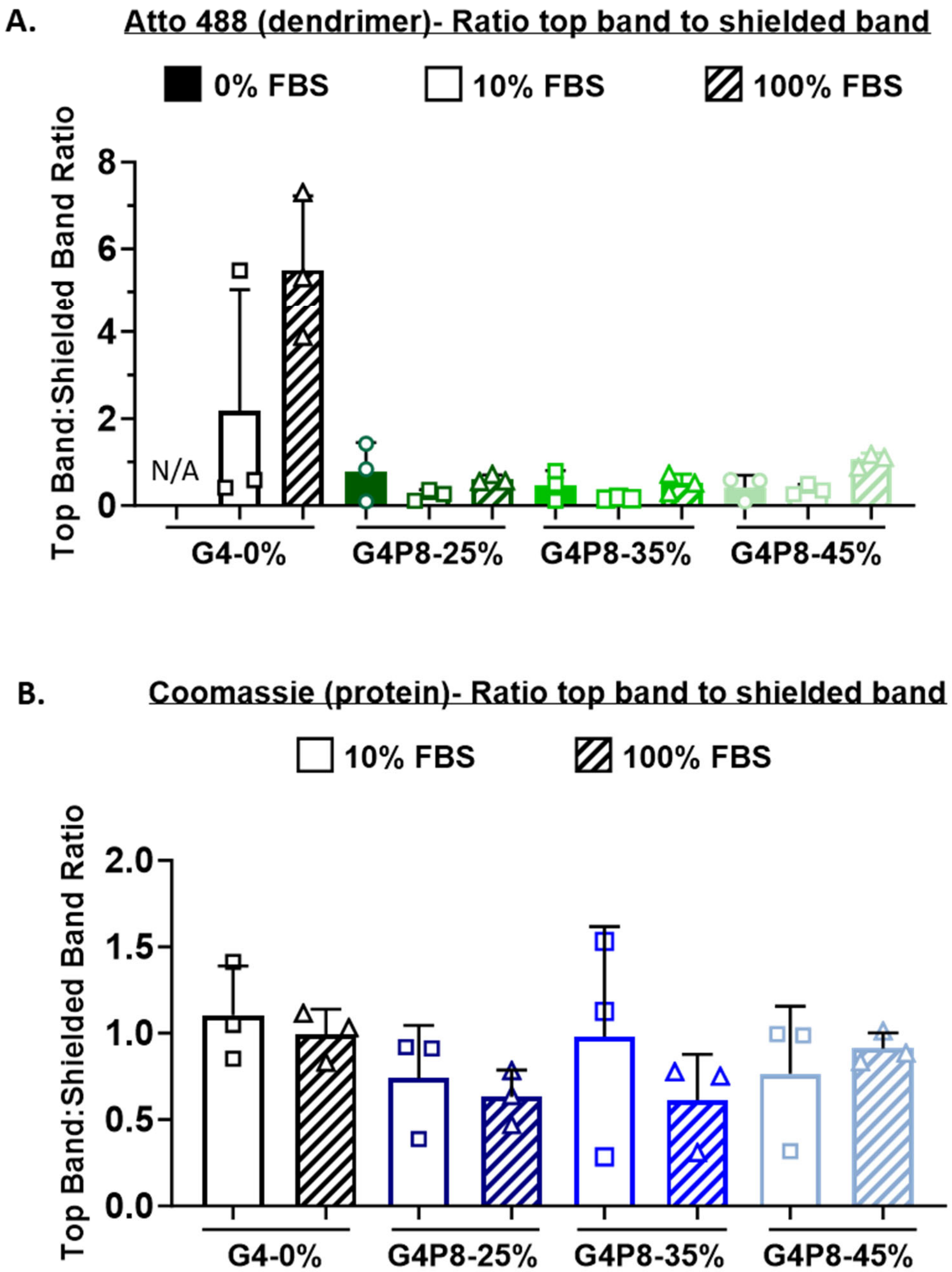

**Supplemental Figure 3: Ratio of shielded band to top band.** The ratio of top band to shielded band for the **(A)** 488nm (dendrimer) and **(B**) Coomassie 700nm (protein) channels were reported to determine relative distribution of proteins and dendrimer migration between the two bands. It was determined that the relative ratio of dendrimers and proteins are consistent across unPEGyalted and PEGylated dendrimers. Note: No shielded band was detected for G4-0% not incubated in FBS.

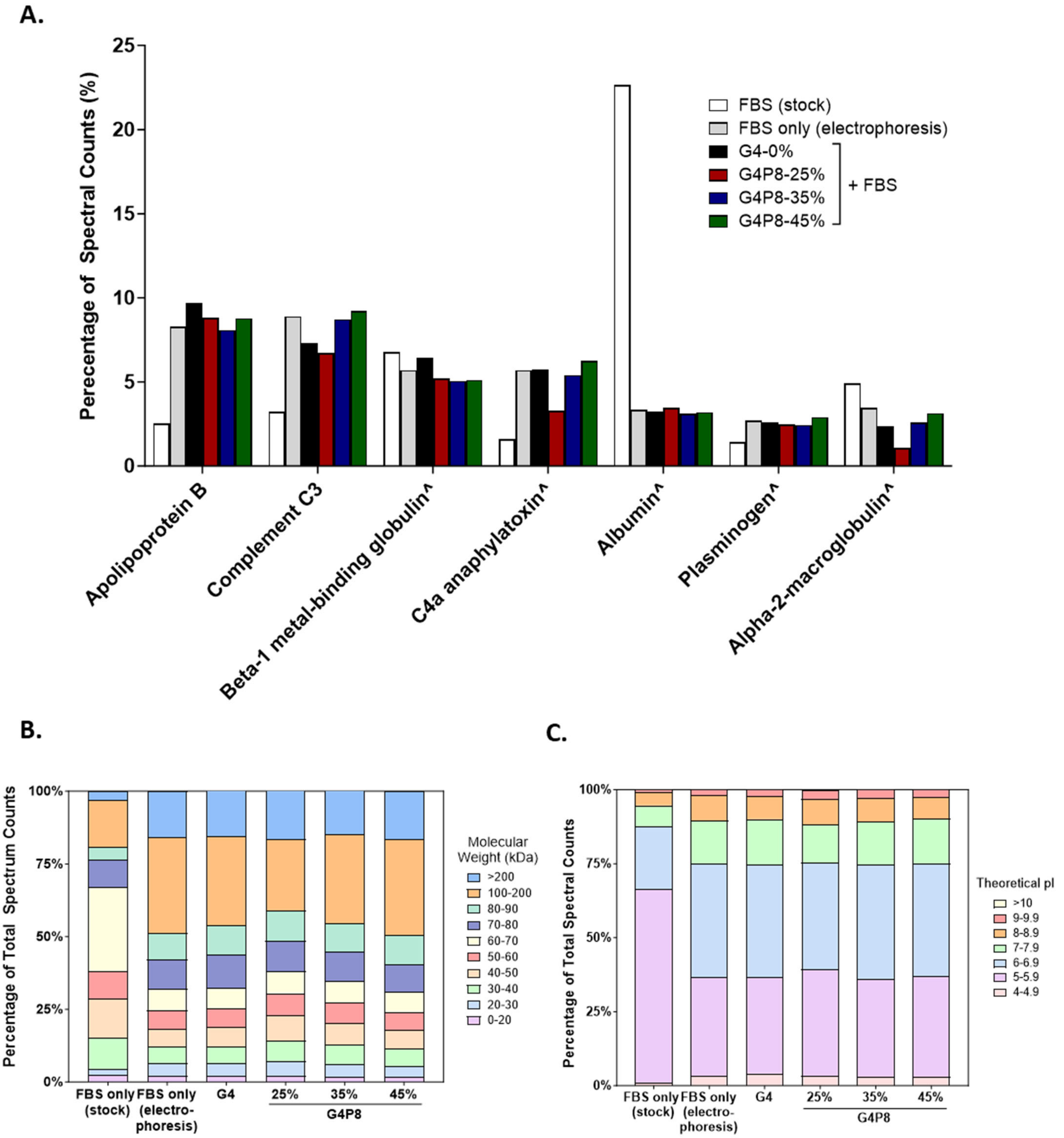

**Supplemental Figure 4: Analysis of identified proteins for PAMAM dendrimers incubated in 100% FBS. (A)** Quantification of the percentage of total spectrum counts for top hit proteins from all experimental conditions. **(B)** Molecular weight and **(C)** isoelectric point (pI) was analyzed based on mass spectroscopy data from Scaffold 5 and Expasy pI/MW tool, respectively.

^denotes proteins found as “cluster of” in the top or shielded band

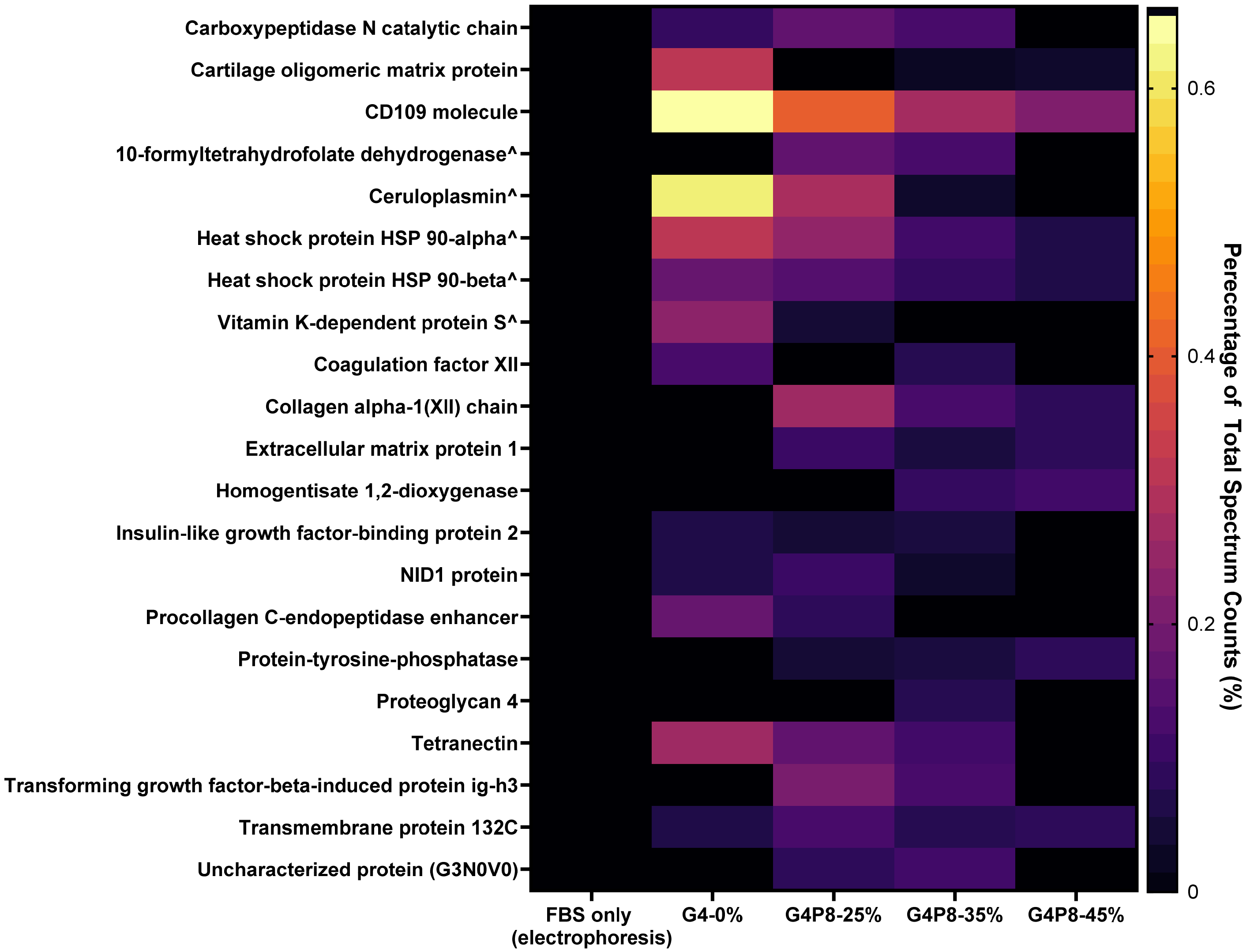

**Supplemental Figure 5: Proteins found associated with dendrimers but not in "electrophoresis" FBS control.**

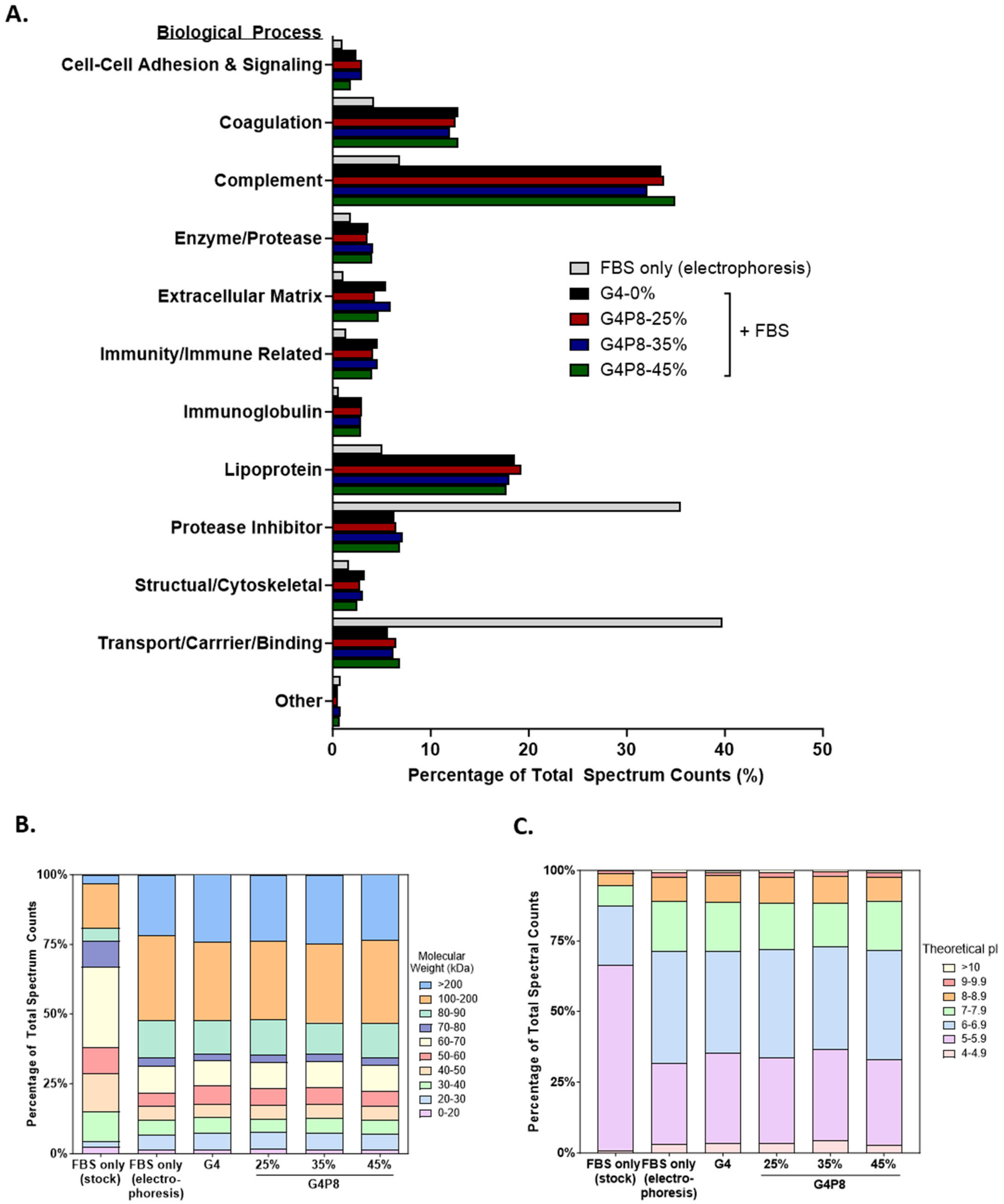

**Supplemental Figure 6: Analysis of identified proteins for dendrimers incubated in 100% FBS. (A)** Identified proteins were sorted by biological process. **(B)** Molecular weight and **(C)** isoelectric point (pI) was analyzed based on mass spectroscopy data from Scaffold 5 and Expasy pI/MW tool, respectively.

| Identified Protein | Accession Number | Molecular Weight | Theoretical pI | Biological Process |
|---|---|---|---|---|
| Cluster of Apolipoprotein B* | E1BNR0 [2] | 516 kDa | 6.24 | Apolipoprotein |
| Cluster of C4a anaphylatoxin | F1MVK1 [4] | 192 kDa | 7.26 | Complement |
| Cluster of Complement factor H | Q28085 [4] | 140 kDa | 6.33 | Complement |
| Complement C3* | Q2UVX4 | 187 kDa | 6.37 | Complement |
| Cluster of Plasminogen | P06868 [2] | 91 kDa | 7.39 | Coagulation |
| Alpha-2-macroglobulin | Q7SIH1 | 168 kDa | 5.68 | Protease Inhibitor |
| Cluster of Albumin | A0A140T897 [2] | 69 kDa | 5.77 | Transport |
| Cluster of C4b-binding protein alpha chain* | Q28065 [3] | 69 kDa | 5.66 | Complement |

**Supplemental Table 1: Top hits from mass spectrometry on PAMAM dendrimer-FBS protein complexes in the "top band" for dendrimers incubated in 100% FBS.** The top band (black arrow, Figure 3A) from native PAGE gels for dendrimers incubated in 100% FBS were excised, digested, and run on mass spectroscopy. The top proteins identified in all experimental conditions are reported in the table. Compared to previously published results, we matched 4 out of 8 proteins, where 3 (indicated by an asterisk) were found in our top hits.

` **Supplemental Table 2: Mass spectroscopy data for dendrimers incubated in 10% FBS**

| Identified Proteins | Accession Number | Biological Process | Molecular Weight | Theoretical pI | G4-0% | | G4P8-25% | | G4P8-35% | | G4P8-45% | | 10% FBS (electrophores | |
|---|---|---|---|---|---|---|---|---|---|---|---|---|---|---|
| | | | | | Total Spectrum Counts | Percentage of Total Spectral Counts | Total Spectrum Counts | Percentage of Total Spectral Counts | Total Spectrum Counts | Percentage of Total Spectral Counts | Total Spectrum Counts | Percentage of Total Spectral Counts | Total Spectrum Counts | Percentage Total Spectr Counts |
| Apolipoprotein B OS=Bos taurus OX=9913 GN=APOB PE=1 SV=3 | E1BNR0 | Apolipoprotein | 516 kDa | 6.24 | 353 | 9.71% | 442 | 8.82% | 446 | 8.07% | 339 | 8.79% | 310 | 8.30% |
| Cluster of C4a anaphylatoxin OS=Bos taurus OX=9913 GN=LOC107131209 PE=4 SV=3 (F1MVK1) | F1MVK1 [3] | Complement | 192 kDa | 7.26 | 208 | 5.72% | 166 | 3.31% | 298 | 5.39% | 241 | 6.25% | 212 | 5.68% |
| Cluster of Alpha-2-macroglobulin OS=Bos taurus OX=9913 GN=A2M PE=1 SV=2 (Q7SIH1) | Q7SIH1 [2] | Protease Inhibitor | 168 kDa | 5.68 | 86 | 2.37% | 53 | 1.06% | 142 | 2.57% | 121 | 3.14% | 129 | 3.45% |
| Complement C3 OS=Bos taurus OX=9913 GN=C3 PE=1 SV=2 | Q2UVX4 | Complement | 187 kDa | 6.37 | 266 | 7.32% | 336 | 6.71% | 481 | 8.71% | 355 | 9.21% | 332 | 8.89% |
| Cluster of Albumin OS=Bos taurus OX=9913 GN=ALB PE=4 SV=1 (A0A140T897) | A0A140T897 [2] | Transport/Carrier/Binding | 69 kDa | 5.77 | 117 | 3.22% | 173 | 3.45% | 171 | 3.10% | 123 | 3.19% | 125 | 3.35% |
| Cluster of Plasminogen OS=Bos taurus OX=9913 GN=PLG PE=1 SV=2 (P06868) | P06868 [2] | Coagulation | 91 kDa | 7.39 | 94 | 2.59% | 123 | 2.46% | 134 | 2.43% | 112 | 2.91% | 101 | 2.70% |
| Cluster of Complement factor H OS=Bos taurus OX=9913 GN=CFH PE=1 SV=3 (Q28085) | Q28085 [6] | Complement | 140 kDa | 6.33 | 36 | 0.99% | 0 | 0.00% | 50 | 0.90% | 48 | 1.25% | 64 | 1.71% |
| Cluster of Thrombospondin-1 OS=Bos taurus OX=9913 GN=THBS1 PE=3 SV=1 (F1N3A1) | F1N3A1 [2] | Cell-Cell Adhesion | 129 kDa | 4.7 | 43 | 1.18% | 50 | 1.00% | 51 | 0.92% | 42 | 1.09% | 59 | 1.58% |
| Fibronectin OS=Bos taurus OX=9913 GN=FN1 PE=4 SV=1 | G5E5A8 | Structual/Cytoskeleton | 276 kDa | 5.79 | 55 | 1.51% | 90 | 1.80% | 100 | 1.81% | 91 | 2.36% | 79 | 2.12% |
| Cluster of Beta-1 metal-binding globulin OS=Bos taurus OX=9913 GN=TF PE=1 SV=2 (G3X6N3) | G3X6N3 [3] | Transport/Carrier/Binding | 78 kDa | 6.63 | 233 | 6.41% | 261 | 5.21% | 279 | 5.05% | 197 | 5.11% | 212 | 5.68% |
| Cluster of Uncharacterized protein OS=Bos taurus OX=9913 PE=1 SV=2 (G5E5T5) | G5E5T5 [2] | Immunity/Immunity Related | 56 kDa | 5.87 | 25 | 0.69% | 34 | 0.68% | 29 | 0.52% | 27 | 0.70% | 26 | 0.70% |
| Apolipoprotein E OS=Bos taurus OX=9913 GN=APOE PE=3 SV=2 | A0A140T881 | Apolipoprotein | 37 kDa | 5.55 | 24 | 0.66% | 38 | 0.76% | 33 | 0.60% | 26 | 0.67% | 22 | 0.59% |
| Cluster of Trypsin OS=Sus scrofa OX=9823 (P00761) | P00761 [2] | Enzyme/Protease | 24 kDa | 8.26 | 58 | 1.60% | 63 | 1.26% | 67 | 1.21% | 46 | 1.19% | 58 | 1.55% |
| Coagulation factor V OS=Bos taurus OX=9913 GN=F5 PE=1 SV=1 | Q28107 | Coagulation | 249 kDa | 5.55 | 28 | 0.77% | 52 | 1.04% | 44 | 0.80% | 28 | 0.73% | 27 | 0.72% |
| Cluster of C4b-binding protein alpha chain OS=Bos taurus OX=9913 GN=C4BPA PE=2 SV=1 (Q28065) | Q28065 [2] | Complement | 69 kDa | 5.66 | 20 | 0.55% | 24 | 0.48% | 29 | 0.52% | 22 | 0.57% | 18 | 0.48% |
| Alpha-1-antiproteinase OS=Bos taurus OX=9913 GN=SERPINA1 PE=1 SV=1 | P34955 | Protease Inhibitor | 46 kDa | 5.98 | 47 | 1.29% | 78 | 1.56% | 72 | 1.30% | 50 | 1.30% | 55 | 1.47% |
| Cluster of von Willebrand factor OS=Bos taurus OX=9913 GN=VWF PE=4 SV=1 (A0A3Q1LLU1) | A0A3Q1LLU1 [2] | Coagulation | 308 kDa | 5.35 | 16 | 0.44% | 21 | 0.42% | 21 | 0.38% | 16 | 0.42% | 20 | 0.54% |
| Complement factor B OS=Bos taurus OX=9913 GN=CFB PE=1 SV=2 | P81187 | Complement | 85 kDa | 7.68 | 38 | 1.05% | 61 | 1.22% | 54 | 0.98% | 43 | 1.12% | 36 | 0.96% |
| Fibrinogen alpha chain OS=Bos taurus OX=9913 GN=FGA PE=4 SV=1 | F6QND5 | Coagulation | 95 kDa | 5.69 | 24 | 0.66% | 35 | 0.70% | 42 | 0.76% | 27 | 0.70% | 27 | 0.72% |
| Alpha-2-HS-glycoprotein OS=Bos taurus OX=9913 GN=AHSG PE=1 SV=2 | P12763 | Protease Inhibitor | 38 kDa | 5.1 | 34 | 0.94% | 52 | 1.04% | 47 | 0.85% | 39 | 1.01% | 42 | 1.12% |
| Cluster of Uncharacterized protein OS=Bos taurus OX=9913 PE=4 SV=3 (F1N160) | F1N160 [3] | Immunity/Immunity Related | 34 kDa | 8.86 | 31 | 0.85% | 50 | 1.00% | 41 | 0.74% | 38 | 0.99% | 29 | 0.78% |
| Cluster of Complement component C7 OS=Bos taurus OX=9913 GN=C7 PE=3 SV=1 (A0A3Q1LI40) | A0A3Q1LI40 [3] | Complement | 93 kDa | 7.08 | 19 | 0.52% | 35 | 0.70% | 27 | 0.49% | 24 | 0.62% | 29 | 0.78% |
| Cluster of Hemoglobin fetal subunit beta OS=Bos taurus OX=9913 PE=1 SV=1 (P02081) | P02081 [4] | Transport/Carrier/Binding | 16 kDa | 6.51 | 30 | 0.83% | 36 | 0.72% | 34 | 0.62% | 29 | 0.75% | 26 | 0.70% |
| ITIH2 protein OS=Bos taurus OX=9913 GN=ITIH2 PE=1 SV=1 | A5D7R6 | Protease Inhibitor | 106 kDa | 7.75 | 62 | 1.71% | 67 | 1.34% | 70 | 1.27% | 46 | 1.19% | 49 | 1.31% |
| Cluster of Uncharacterized protein OS=Bos taurus OX=9913 GN=LOC506828 PE=3 SV=3 (F1MJK3) | F1MJK3 [4] | Protease Inhibitor | 163 kDa | 6.91 | 57 | 1.57% | 72 | 1.44% | 106 | 1.92% | 86 | 2.23% | 76 | 2.04% |
| Uncharacterized protein OS=Bos taurus OX=9913 PE=4 SV=2 | G3N0S9 | Complement | 21 kDa | 5.37 | 17 | 0.47% | 30 | 0.60% | 29 | 0.52% | 22 | 0.57% | 21 | 0.56% |
| Collagen type VI alpha 3 chain OS=Bos taurus OX=9913 GN=COL6A3 PE=1 SV=2 | E1BB91 | Extracellular Matrix | 340 kDa | 5.87 | 47 | 1.29% | 71 | 1.42% | 68 | 1.23% | 42 | 1.09% | 39 | 1.04% |
| Fibrinogen gamma-B chain OS=Bos taurus OX=9913 GN=FGG PE=4 SV=1 | F1MGU7 (+1) | Coagulation | 50 kDa | 5.38 | 13 | 0.36% | 26 | 0.52% | 21 | 0.38% | 19 | 0.49% | 18 | 0.48% |
| Cluster of Fibrinogen beta chain OS=Bos taurus OX=9913 GN=FGB PE=4 SV=1 (A0A3Q1MG04) | A0A3Q1MG04 [4] | Coagulation | 57 kDa | 8.33 | 14 | 0.39% | 25 | 0.50% | 27 | 0.49% | 20 | 0.52% | 22 | 0.59% |
| Complement component C9 OS=Bos taurus OX=9913 GN=C9 PE=2 SV=1 | Q3MHN2 | Complement | 62 kDa | 5.57 | 8 | 0.22% | 10 | 0.20% | 7 | 0.13% | 8 | 0.21% | 8 | 0.21% |
| Uncharacterized protein OS=Bos taurus OX=9913 PE=1 SV=1 | A0A3Q1M3L6 | Immunoglobulin | 40 kDa | 5.16 | 28 | 0.77% | 40 | 0.80% | 47 | 0.85% | 30 | 0.78% | 27 | 0.72% |
| Cluster of Clusterin OS=Bos taurus OX=9913 GN=CLU PE=1 SV=1 (P17697) | P17697 [2] | Apolipoprotein | 51 kDa | 5.72 | 19 | 0.52% | 34 | 0.68% | 36 | 0.65% | 19 | 0.49% | 20 | 0.54% |
| Cluster of Fructose-bisphosphate aldolase B OS=Bos taurus OX=9913 GN=ALDOB PE=2 SV=1 (Q3T0S5) | Q3T0S5 [2] | Enzyme/Protease | 40 kDa | 8.45 | 7 | 0.19% | 14 | 0.28% | 14 | 0.25% | 13 | 0.34% | 16 | 0.43% |
| Collagen type VI alpha 1 chain OS=Bos taurus OX=9913 GN=COL6A1 PE=1 SV=1 | E1BI98 | Extracellular Matrix | 109 kDa | 5.18 | 18 | 0.50% | 24 | 0.48% | 23 | 0.42% | 18 | 0.47% | 14 | 0.37% |
| Hemoglobin subunit alpha OS=Bos taurus OX=9913 GN=HBA PE=1 SV=2 | P01966 | Transport/Carrier/Binding | 15 kDa | 8.19 | 15 | 0.41% | 18 | 0.36% | 18 | 0.33% | 12 | 0.31% | 15 | 0.40% |
| Peptidoglycan recognition protein 1 OS=Bos taurus OX=9913 GN=PGLYRP1 PE=1 SV=1 | Q8SPP7 | Immunity/Immunity Related | 21 kDa | 9.38 | 7 | 0.19% | 7 | 0.14% | 6 | 0.11% | 6 | 0.16% | 3 | 0.08% |
| Tubulin beta chain OS=Bos taurus OX=9913 GN=TUBB1 PE=3 SV=1 | A0A3Q1M442 (+1) | Structual/Cytoskeleton | 55 kDa | 5.41 | 7 | 0.19% | 16 | 0.32% | 16 | 0.29% | 12 | 0.31% | 11 | 0.29% |
| Adiponectin B OS=Bos taurus OX=9913 GN=C1QC PE=4 SV=1 | A0A3B0IZF8 | Complement | 26 kDa | 8.62 | 6 | 0.17% | 7 | 0.14% | 6 | 0.11% | 7 | 0.18% | 5 | 0.13% |
| Apolipoprotein H OS=Bos taurus OX=9913 GN=APOH PE=4 SV=1 | A0A140T843 (+1) | Apolipoprotein | 38 kDa | 8.55 | 16 | 0.44% | 29 | 0.58% | 26 | 0.47% | 23 | 0.60% | 26 | 0.70% |
| Complement C5a anaphylatoxin OS=Bos taurus OX=9913 GN=C5 PE=1 SV=3 | F1MY85 | Complement | 189 kDa | 6.16 | 13 | 0.36% | 34 | 0.68% | 63 | 1.14% | 48 | 1.25% | 43 | 1.15% |
| Platelet-activating factor acetylhydrolase OS=Bos taurus OX=9913 GN=PLA2G7 PE=2 SV=1 | Q1RML9 (+1) | Coagulation | 50 kDa | 5.96 | 6 | 0.17% | 9 | 0.18% | 5 | 0.09% | 4 | 0.10% | 3 | 0.08% |
| Apolipoprotein A-I OS=Bos taurus OX=9913 GN=APOA1 PE=1 SV=3 | P15497 | Apolipoprotein | 30 kDa | 5.36 | 21 | 0.58% | 39 | 0.78% | 44 | 0.80% | 17 | 0.44% | 12 | 0.32% |
| Cluster of Antithrombin-III OS=Bos taurus OX=9913 GN=SERPINC1 PE=3 SV=1 (A0A3Q1NJR8) | A0A3Q1NJR8 [2] | Coagulation | 60 kDa | 8.85 | 17 | 0.47% | 19 | 0.38% | 24 | 0.43% | 9 | 0.23% | 13 | 0.35% |
| Cluster of Bradykinin OS=Bos taurus OX=9913 GN=KNG1 PE=4 SV=1 (A0A140T8C8) | A0A140T8C8 [2] | Protease Inhibitor | 69 kDa | 6.09 | 18 | 0.50% | 23 | 0.46% | 20 | 0.36% | 16 | 0.42% | 13 | 0.35% |
| Cluster of Tubulin beta-5 chain OS=Bos taurus OX=9913 GN=TUBB5 PE=2 SV=1 (Q2KJD0) | Q2KJD0 [8] | Structual/Cytoskeleton | 50 kDa | 4.78 | 5 | 0.14% | 10 | 0.20% | 6 | 0.11% | 3 | 0.08% | 3 | 0.08% |
| Fetuin-B OS=Bos taurus OX=9913 GN=FETUB PE=1 SV=1 | Q58D62 | Protease Inhibitor | 43 kDa | 5.59 | 21 | 0.58% | 26 | 0.52% | 26 | 0.47% | 20 | 0.52% | 19 | 0.51% |
| Serpin family D member 1 OS=Bos taurus OX=9913 GN=SERPIND1 PE=3 SV=1 | F6R4P6 | Coagulation | 63 kDa | 6.23 | 11 | 0.30% | 8 | 0.16% | 10 | 0.18% | 7 | 0.18% | 6 | 0.16% |
| Apolipoprotein D OS=Bos taurus OX=9913 GN=APOD PE=3 SV=3 | F1MS32 | Apolipoprotein | 24 kDa | 5.07 | 4 | 0.11% | 6 | 0.12% | 4 | 0.07% | 4 | 0.10% | 4 | 0.11% |
| Ig-like domain-containing protein OS=Bos taurus OX=9913 PE=1 SV=1 | A0A3Q1LUE9 (+1) | Immunoglobulin | 17 kDa | 8.76 | 4 | 0.11% | 4 | 0.08% | 3 | 0.05% | 3 | 0.08% | 4 | 0.11% |
| Laminin subunit alpha 2 OS=Bos taurus OX=9913 GN=LAMA2 PE=4 SV=1 | A0A3Q1MFB8 (+1) | Extracellular Matrix | 224 kDa | 5.6 | 4 | 0.11% | 3 | 0.06% | 0 | 0.00% | 0 | 0.00% | 3 | 0.08% |
| Vitronectin OS=Bos taurus OX=9913 GN=VTN PE=1 SV=1 | Q3ZBS7 | Cell-Cell Adhesion | 54 kDa | 5.79 | 10 | 0.28% | 15 | 0.30% | 15 | 0.27% | 11 | 0.29% | 11 | 0.29% |
| Adenosylhomocysteinase OS=Bos taurus OX=9913 GN=AHCY PE=2 SV=3 | Q3MHL4 | Structual/Cytoskeleton | 48 kDa | 5.88 | 13 | 0.36% | 22 | 0.44% | 19 | 0.34% | 9 | 0.23% | 7 | 0.19% |
| Cluster of Inter-alpha-trypsin inhibitor heavy chain H4 OS=Bos taurus OX=9913 GN=ITIH4 PE=3 SV=3 (F1MM | F1MMD7 [3] | Protease Inhibitor | 102 kDa | 5.92 | 24 | 0.66% | 29 | 0.58% | 32 | 0.58% | 24 | 0.62% | 27 | 0.72% |
| Cluster of Mannan binding lectin serine peptidase 1 OS=Bos taurus OX=9913 GN=MASP1 PE=4 SV=1 (F1MV | F1MVS9 [2] | Complement | 81 kDa | 4.93 | 8 | 0.22% | 9 | 0.18% | 5 | 0.09% | 6 | 0.16% | 7 | 0.19% |
| Complement component 8 subunit beta OS=Bos taurus OX=9913 GN=C8B PE=3 SV=3 | F1N102 | Complement | 67 kDa | 8.15 | 3 | 0.08% | 0 | 0.00% | 2 | 0.04% | 0 | 0.00% | 5 | 0.13% |
| Complement component C6 OS=Bos taurus OX=9913 GN=C6 PE=3 SV=2 | F1MM86 (+1) | Complement | 105 kDa | 6.74 | 3 | 0.08% | 4 | 0.08% | 0 | 0.00% | 0 | 0.00% | 0 | 0.00% |
| Immunoglobulin J chain OS=Bos taurus OX=9913 GN=JCHAIN PE=1 SV=1 | Q3SYR8 | Immunoglobulin | 18 kDa | 4.92 | 3 | 0.08% | 6 | 0.12% | 3 | 0.05% | 4 | 0.10% | 5 | 0.13% |
| Inter-alpha-trypsin inhibitor heavy chain H3 OS=Bos taurus OX=9913 GN=ITIH3 PE=3 SV=1 | A0A3Q1LQ21 (+1) | Transport/Carrier/Binding | 99 kDa | 5.91 | 24 | 0.66% | 28 | 0.56% | 30 | 0.54% | 21 | 0.54% | 15 | 0.40% |
| Laminin subunit gamma 1 OS=Bos taurus OX=9913 GN=LAMC1 PE=1 SV=2 | F1MD77 | Extracellular Matrix | 178 kDa | 4.97 | 3 | 0.08% | 0 | 0.00% | 3 | 0.05% | 0 | 0.00% | 0 | 0.00% |
| Angiotensinogen OS=Bos taurus OX=9913 GN=AGT PE=1 SV=2 | P01017 | Protease Inhibitor | 51 kDa | 6.49 | 37 | 1.02% | 51 | 1.02% | 52 | 0.94% | 29 | 0.75% | 31 | 0.83% |
| Complement factor properdin OS=Bos taurus OX=9913 GN=CFP PE=4 SV=1 | A0A3Q1MHU8 (+1) | Complement | 50 kDa | 8.51 | 2 | 0.06% | 0 | 0.00% | 3 | 0.05% | 0 | 0.00% | 4 | 0.11% |
| Laminin subunit beta 1 OS=Bos taurus OX=9913 GN=LAMB1 PE=4 SV=1 | A0A3Q1MY30 (+1) | Extracellular Matrix | 219 kDa | 5.04 | 2 | 0.06% | 2 | 0.04% | 0 | 0.00% | 2 | 0.05% | 0 | 0.00% |
| 2-iminobutanoate/2-iminopropanoate deaminase OS=Bos taurus OX=9913 GN=RIDA PE=2 SV=3 | Q3T114 | Enzyme/Protease | 14 kDa | 6.15 | 4 | 0.11% | 6 | 0.12% | 6 | 0.11% | 3 | 0.08% | 4 | 0.11% |
| 6-phosphogluconate dehydrogenase, decarboxylating OS=Bos taurus OX=9913 GN=PGD PE=1 SV=1 | A0A3S5ZPM3 | Enzyme/Protease | 61 kDa | 8.46 | 2 | 0.06% | 3 | 0.06% | 4 | 0.07% | 0 | 0.00% | 0 | 0.00% |

| | | | | | | | | | | | | | | |
|---|---|---|---|---|---|---|---|---|---|---|---|---|---|---|
| 72 kDa type IV collagenase OS=Bos taurus OX=9913 GN=MMP2 PE=2 SV=1 | Q9GLE5 | Enzyme/Protease | 74 kDa | 5.09 | 0 | 0.00% | 3 | 0.06% | 0 | 0.00% | 0 | 0.00% | 0 | 0.00% |
| Actin-depolymerizing factor OS=Bos taurus OX=9913 GN=GSN PE=1 SV=2 | F1N1I6 | Structual/Cytoskeleton | 92 kDa | 5.98 | 17 | 0.47% | 30 | 0.60% | 26 | 0.47% | 19 | 0.49% | 16 | 0.43% |
| ADAM metallopeptidase with thrombospondin type 1 motif 13 OS=Bos taurus OX=9913 GN=ADAMTS13 PE=4 | F1MVP0 | Enzyme/Protease | 103 kDa | 6.42 | 12 | 0.33% | 20 | 0.40% | 21 | 0.38% | 14 | 0.36% | 11 | 0.29% |
| Adiponectin M OS=Bos taurus OX=9913 GN=C1QTNF3 PE=2 SV=1 | A7MB82 | Cell-Cell Adhesion | 27 kDa | 5.91 | 0 | 0.00% | 0 | 0.00% | 3 | 0.05% | 0 | 0.00% | 0 | 0.00% |
| Adiponectin OS=Bos taurus OX=9913 GN=ADIPOQ PE=4 SV=1 | A0A3Q1M564 (+1) | Cell-Cell Adhesion | 35 kDa | 5.33 | 11 | 0.30% | 23 | 0.46% | 18 | 0.33% | 12 | 0.31% | 11 | 0.29% |
| Alpha-1-acid glycoprotein OS=Bos taurus OX=9913 GN=ORM1 PE=2 SV=1 | Q3SZR3 (+1) | Transport/Carrier/Binding | 23 kDa | 5.67 | 0 | 0.00% | 5 | 0.10% | 4 | 0.07% | 0 | 0.00% | 0 | 0.00% |
| Alpha-1B-glycoprotein OS=Bos taurus OX=9913 GN=A1BG PE=1 SV=1 | A0A3Q1MJT2 | Immunity/Immunity Related | 62 kDa | 5.76 | 4 | 0.11% | 7 | 0.14% | 6 | 0.11% | 5 | 0.13% | 6 | 0.16% |
| Alpha-1-microglobulin OS=Bos taurus OX=9913 GN=KIF12 PE=3 SV=3 | F1MMK9 | N/A | 53 kDa | 8.87 | 10 | 0.28% | 10 | 0.20% | 12 | 0.22% | 10 | 0.26% | 11 | 0.29% |
| Alpha-2-antiplasmin OS=Bos taurus OX=9913 GN=SERPINF2 PE=1 SV=2 | P28800 | Protease Inhibitor | 55 kDa | 5.71 | 11 | 0.30% | 26 | 0.52% | 25 | 0.45% | 13 | 0.34% | 14 | 0.37% |
| Alpha-fetoprotein OS=Bos taurus OX=9913 GN=AFP PE=1 SV=1 | A0A3Q1MIW0 (+1) | Transport/Carrier/Binding | 68 kDa | 6.17 | 3 | 0.08% | 13 | 0.26% | 14 | 0.25% | 6 | 0.16% | 11 | 0.29% |
| Amine oxidase OS=Bos taurus OX=9913 GN=AOC3 PE=1 SV=2 | E1BD43 | Enzyme/Protease | 86 kDa | 6.86 | 3 | 0.08% | 4 | 0.08% | 3 | 0.05% | 0 | 0.00% | 0 | 0.00% |
| Annexin A2 OS=Bos taurus OX=9913 GN=ANXA2 PE=1 SV=2 | P04272 | Cell-Cell Adhesion | 39 kDa | 6.9 | 5 | 0.14% | 0 | 0.00% | 0 | 0.00% | 0 | 0.00% | 0 | 0.00% |
| Arginase-1 OS=Bos taurus OX=9913 GN=ARG1 PE=2 SV=1 | Q2KJ64 | Enzyme/Protease | 35 kDa | 6.09 | 3 | 0.08% | 0 | 0.00% | 2 | 0.04% | 0 | 0.00% | 0 | 0.00% |
| Argininosuccinate synthase OS=Bos taurus OX=9913 GN=ASS1 PE=2 SV=1 | P14568 | Enzyme/Protease | 46 kDa | 7.15 | 0 | 0.00% | 4 | 0.08% | 5 | 0.09% | 2 | 0.05% | 0 | 0.00% |
| Aspartate aminotransferase, cytoplasmic OS=Bos taurus OX=9913 GN=GOT1 PE=1 SV=3 | P33097 | Enzyme/Protease | 46 kDa | 7.09 | 3 | 0.08% | 8 | 0.16% | 10 | 0.18% | 7 | 0.18% | 2 | 0.05% |
| Beta-1,4-galactosyltransferase 1 OS=Bos taurus OX=9913 GN=B4GALT1 PE=1 SV=3 | P08037 | Enzyme/Protease | 45 kDa | 9.38 | 0 | 0.00% | 2 | 0.04% | 0 | 0.00% | 0 | 0.00% | 0 | 0.00% |
| Beta-hexosaminidase OS=Bos taurus OX=9913 GN=HEXB PE=3 SV=1 | H7BWW2 | Enzyme/Protease | 61 kDa | 6.54 | 0 | 0.00% | 3 | 0.06% | 0 | 0.00% | 0 | 0.00% | 0 | 0.00% |
| C3/C5 convertase OS=Bos taurus OX=9913 GN=C2 PE=4 SV=1 | A0A3S5ZPC6 (+1) | Enzyme/Protease | 87 kDa | 8.86 | 0 | 0.00% | 2 | 0.04% | 0 | 0.00% | 3 | 0.08% | 2 | 0.05% |
| Carbonic anhydrase 2 OS=Bos taurus OX=9913 GN=CA2 PE=1 SV=3 | P00921 | Enzyme/Protease | 29 kDa | 6.4 | 6 | 0.17% | 8 | 0.16% | 8 | 0.14% | 6 | 0.16% | 4 | 0.11% |
| Carboxylic ester hydrolase OS=Bos taurus OX=9913 GN=ACHE PE=3 SV=1 | A0A140T835 (+1) | Enzyme/Protease | 68 kDa | 5.72 | 2 | 0.06% | 5 | 0.10% | 5 | 0.09% | 5 | 0.13% | 4 | 0.11% |
| Carboxypeptidase B2 OS=Bos taurus OX=9913 GN=CPB2 PE=1 SV=1 | Q2KIG3 | Enzyme/Protease | 49 kDa | 8.49 | 12 | 0.33% | 18 | 0.36% | 23 | 0.42% | 12 | 0.31% | 13 | 0.35% |
| Carboxypeptidase N catalytic chain OS=Bos taurus OX=9913 GN=CPN1 PE=3 SV=1 | G5E5V0 (+1) | Enzyme/Protease | 53 kDa | 8.76 | 4 | 0.11% | 9 | 0.18% | 8 | 0.14% | 0 | 0.00% | 0 | 0.00% |
| Cartilage oligomeric matrix protein OS=Bos taurus OX=9913 GN=COMP PE=1 SV=2 | P35445 | Cell-Cell Adhesion | 82 kDa | 4.36 | 12 | 0.33% | 0 | 0.00% | 2 | 0.04% | 2 | 0.05% | 0 | 0.00% |
| Catalase OS=Bos taurus OX=9913 GN=CAT PE=1 SV=3 | P00432 | Enzyme/Protease | 60 kDa | 6.82 | 2 | 0.06% | 4 | 0.08% | 4 | 0.07% | 4 | 0.10% | 8 | 0.21% |
| Cathepsin C OS=Bos taurus OX=9913 GN=CTSC PE=3 SV=1 | F1N455 | Enzyme/Protease | 52 kDa | 6.74 | 5 | 0.14% | 8 | 0.16% | 7 | 0.13% | 4 | 0.10% | 4 | 0.11% |
| Cation-independent mannose-6-phosphate receptor OS=Bos taurus OX=9913 GN=IGF2R PE=1 SV=2 | P08169 | Transport/Carrier/Binding | 275 kDa | 5.57 | 56 | 1.54% | 90 | 1.80% | 92 | 1.67% | 73 | 1.89% | 68 | 1.82% |
| CD109 molecule OS=Bos taurus OX=9913 GN=CD109 PE=3 SV=1 | A0A3Q1LTK9 | Cell-Cell Adhesion | 162 kDa | 5.38 | 24 | 0.66% | 21 | 0.42% | 16 | 0.29% | 9 | 0.23% | 0 | 0.00% |
| Cell adhesion molecule L1 like OS=Bos taurus OX=9913 GN=CHL1 PE=3 SV=1 | A0A3Q1M2M5 | Cell-Cell Adhesion | 130 kDa | 5.19 | 3 | 0.08% | 0 | 0.00% | 0 | 0.00% | 0 | 0.00% | 0 | 0.00% |
| Chitinase OS=Bos taurus OX=9913 GN=CHIA PE=3 SV=2 | F1MH27 (+1) | Enzyme/Protease | 52 kDa | 5.36 | 5 | 0.14% | 3 | 0.06% | 0 | 0.00% | 3 | 0.08% | 0 | 0.00% |
| Chondroitin sulfate proteoglycan 4 OS=Bos taurus OX=9913 GN=CSPG4 PE=4 SV=3 | F1MY84 | Cell-Cell Adhesion | 251 kDa | 5.24 | 0 | 0.00% | 4 | 0.08% | 0 | 0.00% | 0 | 0.00% | 0 | 0.00% |
| Cluster of 10-formyltetrahydrofolate dehydrogenase OS=Bos taurus OX=9913 GN=ALDH1L1 PE=1 SV=3 (E1E | E1BMG9 [3] | Enzyme/Protease | 99 kDa | 5.53 | 0 | 0.00% | 9 | 0.18% | 8 | 0.14% | 0 | 0.00% | 0 | 0.00% |
| Cluster of 15-oxoprostaglandin 13-reductase OS=Bos taurus OX=9913 GN=PTGR1 PE=3 SV=1 (F1N2W0) | F1N2W0 [2] | Enzyme/Protease | 36 kDa | 7.65 | 0 | 0.00% | 3 | 0.06% | 5 | 0.09% | 0 | 0.00% | 0 | 0.00% |
| Cluster of 2-phospho-D-glycerate hydro-lyase OS=Bos taurus OX=9913 GN=ENO1 PE=3 SV=2 (F1MB08) | F1MB08 [3] | Enzyme/Protease | 54 kDa | 9.2 | 8 | 0.22% | 28 | 0.56% | 26 | 0.47% | 19 | 0.49% | 12 | 0.32% |
| Cluster of 4-hydroxyphenylpyruvate dioxygenase OS=Bos taurus OX=9913 GN=HPD PE=1 SV=1 (A0A3Q1N2 | A0A3Q1N2P4 [2] | Enzyme/Protease | 45 kDa | 6.29 | 9 | 0.25% | 12 | 0.24% | 12 | 0.22% | 10 | 0.26% | 6 | 0.16% |
| Cluster of 4-trimethylaminobutyraldehyde dehydrogenase OS=Bos taurus OX=9913 GN=ALDH9A1 PE=2 SV= | Q2KJH9 [2] | Enzyme/Protease | 54 kDa | 5.84 | 0 | 0.00% | 0 | 0.00% | 3 | 0.05% | 0 | 0.00% | 0 | 0.00% |
| Cluster of Actin, cytoplasmic 1 OS=Bos taurus OX=9913 GN=ACTB PE=1 SV=1 (P60712) | P60712 [4] | Structual/Cytoskeleton | 42 kDa | 5.31 | 19 | 0.52% | 28 | 0.56% | 27 | 0.49% | 21 | 0.54% | 15 | 0.40% |
| Cluster of Alpha-actinin-4 OS=Bos taurus OX=9913 GN=ACTN4 PE=1 SV=1 (A0A3Q1MFS2) | A0A3Q1MFS2 [5] | Cell-Cell Adhesion | 115 kDa | 5.64 | 3 | 0.08% | 0 | 0.00% | 0 | 0.00% | 0 | 0.00% | 0 | 0.00% |
| Cluster of Betaine--homocysteine S-methyltransferase 1 OS=Bos taurus OX=9913 GN=BHMT PE=2 SV=1 (Q5 | Q5I597 [2] | Enzyme/Protease | 45 kDa | 6.4 | 14 | 0.39% | 17 | 0.34% | 19 | 0.34% | 16 | 0.42% | 15 | 0.40% |
| Cluster of CD5 molecule like OS=Bos taurus OX=9913 GN=CD5L PE=2 SV=1 (A6QNW7) | A6QNW7 [3] | Immunity/Immunity Related | 50 kDa | 5.24 | 0 | 0.00% | 3 | 0.06% | 3 | 0.05% | 0 | 0.00% | 0 | 0.00% |
| Cluster of Cerebellin 4 precursor OS=Bos taurus OX=9913 GN=CBLN4 PE=4 SV=1 (A0A3Q1M0U1) | A0A3Q1M0U1 [2] | Cell-Cell Adhesion | 22 kDa | 6.16 | 0 | 0.00% | 0 | 0.00% | 3 | 0.05% | 4 | 0.10% | 4 | 0.11% |
| Cluster of Ceruloplasmin OS=Bos taurus OX=9913 GN=CP PE=1 SV=1 (A0A3Q1NJB1) | A0A3Q1NJB1 [3] | Transport/Carrier/Binding | 121 kDa | 5.67 | 23 | 0.63% | 15 | 0.30% | 3 | 0.05% | 0 | 0.00% | 0 | 0.00% |
| Cluster of Coagulation factor XIII A chain OS=Bos taurus OX=9913 GN=F13A1 PE=3 SV=1 (A0A3Q1LTB9) | A0A3Q1LTB9 [3] | Coagulation | 76 kDa | 6 | 14 | 0.39% | 21 | 0.42% | 23 | 0.42% | 9 | 0.23% | 13 | 0.35% |
| Cluster of Collagen type VI alpha 2 chain OS=Bos taurus OX=9913 GN=COL6A2 PE=1 SV=3 (F1MKG2) | F1MKG2 [2] | Extracellular Matrix | 105 kDa | 6.2 | 0 | 0.00% | 3 | 0.06% | 0 | 0.00% | 0 | 0.00% | 0 | 0.00% |
| Cluster of Collectin-43 OS=Bos taurus OX=9913 GN=CL43 PE=1 SV=2 (P42916) | P42916 [4] | Immunity/Immunity Related | 34 kDa | 5.04 | 0 | 0.00% | 9 | 0.18% | 28 | 0.51% | 14 | 0.36% | 10 | 0.27% |
| Cluster of Complement C1s subcomponent OS=Bos taurus OX=9913 GN=C1S PE=2 SV=2 (Q0VCX1) | Q0VCX1 [2] | Complement | 77 kDa | 5.06 | 6 | 0.17% | 10 | 0.20% | 12 | 0.22% | 9 | 0.23% | 6 | 0.16% |
| Cluster of Complement factor I OS=Bos taurus OX=9913 GN=CFI PE=1 SV=1 (A0A3Q1MF14) | A0A3Q1MF14 [2] | Complement | 69 kDa | 8.02 | 14 | 0.39% | 26 | 0.52% | 27 | 0.49% | 14 | 0.36% | 11 | 0.29% |
| Cluster of Dihydrodiol dehydrogenase 3 OS=Bos taurus OX=9913 PE=2 SV=1 (P52898) | P52898 [7] | Enzyme/Protease | 37 kDa | 7.65 | 0 | 0.00% | 0 | 0.00% | 3 | 0.05% | 0 | 0.00% | 0 | 0.00% |
| Cluster of Dihydropyrimidinase OS=Bos taurus OX=9913 GN=DPYS PE=1 SV=3 (E1BFN6) | E1BFN6 [2] | Enzyme/Protease | 56 kDa | 6.29 | 13 | 0.36% | 20 | 0.40% | 17 | 0.31% | 8 | 0.21% | 10 | 0.27% |
| Cluster of Elongation factor 1-alpha 1 OS=Bos taurus OX=9913 GN=EEF1A1 PE=1 SV=1 (P68103) | P68103 [3] | Other | 50 kDa | 9.1 | 4 | 0.11% | 8 | 0.16% | 11 | 0.20% | 6 | 0.16% | 4 | 0.11% |
| Cluster of Elongation factor 2 OS=Bos taurus OX=9913 GN=EEF2 PE=2 SV=3 (Q3SYU2) | Q3SYU2 [3] | Other | 95 kDa | 6.41 | 5 | 0.14% | 4 | 0.08% | 6 | 0.11% | 4 | 0.10% | 4 | 0.11% |
| Cluster of Ferritin light chain OS=Bos taurus OX=9913 GN=FTL PE=2 SV=3 (O46415) | O46415 [2] | Transport/Carrier/Binding | 20 kDa | 5.88 | 0 | 0.00% | 0 | 0.00% | 4 | 0.07% | 2 | 0.05% | 3 | 0.08% |
| Cluster of Fructose-1,6-bisphosphatase 1 OS=Bos taurus OX=9913 GN=FBP1 PE=2 SV=3 (Q3SZB7) | Q3SZB7 [3] | Enzyme/Protease | 37 kDa | 6.51 | 0 | 0.00% | 2 | 0.04% | 0 | 0.00% | 0 | 0.00% | 0 | 0.00% |
| Cluster of Glyceraldehyde-3-phosphate dehydrogenase OS=Bos taurus OX=9913 GN=GAPDH PE=1 SV=4 (P | P10096 [2] | Enzyme/Protease | 36 kDa | 8.52 | 11 | 0.30% | 20 | 0.40% | 20 | 0.36% | 11 | 0.29% | 13 | 0.35% |
| Cluster of Glycogen phosphorylase, liver form OS=Bos taurus OX=9913 GN=PYGL PE=2 SV=1 (Q0VCM4) | Q0VCM4 [6] | Enzyme/Protease | 97 kDa | 6.7 | 7 | 0.19% | 25 | 0.50% | 33 | 0.60% | 10 | 0.26% | 12 | 0.32% |
| Cluster of Heat shock 70 kDa protein 1B OS=Bos taurus OX=9913 GN=HSPA1B PE=2 SV=1 (Q27965) | Q27965 [8] | Other | 70 kDa | 5.68 | 5 | 0.14% | 17 | 0.34% | 17 | 0.31% | 15 | 0.39% | 8 | 0.21% |
| Cluster of Heat shock protein HSP 90-alpha OS=Bos taurus OX=9913 GN=HSP90AA1 PE=1 SV=3 (Q76LV2) | Q76LV2 [2] | Other | 85 kDa | 4.92 | 12 | 0.33% | 13 | 0.26% | 7 | 0.13% | 3 | 0.08% | 0 | 0.00% |
| Cluster of Heat shock protein HSP 90-beta OS=Bos taurus OX=9913 GN=HSP90AB1 PE=1 SV=2 (G5E507) | G5E507 [2] | Other | 82 kDa | 4.92 | 7 | 0.19% | 8 | 0.16% | 6 | 0.11% | 3 | 0.08% | 0 | 0.00% |
| Cluster of Ig-like domain-containing protein OS=Bos taurus OX=9913 PE=4 SV=1 (A0A3Q1LSF0) | A0A3Q1LSF0 [3] | Immunoglobulin | 15 kDa | 9.82 | 0 | 0.00% | 5 | 0.10% | 8 | 0.14% | 0 | 0.00% | 6 | 0.16% |
| Cluster of Ig-like domain-containing protein OS=Bos taurus OX=9913 PE=4 SV=1 (A0A3Q1M0K3) | A0A3Q1M0K3 [8] | Immunoglobulin | 23 kDa | 6.97 | 0 | 0.00% | 13 | 0.26% | 13 | 0.24% | 0 | 0.00% | 6 | 0.16% |
| Cluster of Ig-like domain-containing protein OS=Bos taurus OX=9913 PE=4 SV=2 (G5E5V1) | G5E5V1 [2] | Immunoglobulin | 14 kDa | 8.02 | 0 | 0.00% | 3 | 0.06% | 2 | 0.04% | 0 | 0.00% | 2 | 0.05% |
| Cluster of Inter-alpha-trypsin inhibitor heavy chain H1 OS=Bos taurus OX=9913 GN=ITIH1 PE=1 SV=1 (A0A3C | A0A3Q1LW96 [3] | Protease Inhibitor | 101 kDa | 7.03 | 44 | 1.21% | 40 | 0.80% | 40 | 0.72% | 20 | 0.52% | 21 | 0.56% |
| Cluster of Latent transforming growth factor beta binding protein 4 OS=Bos taurus OX=9913 GN=LTBP4 PE=4 | G3MZT8 [2] | Extracellular Matrix | 173 kDa | 5.11 | 0 | 0.00% | 0 | 0.00% | 6 | 0.11% | 0 | 0.00% | 0 | 0.00% |
| Cluster of Lipocln_cytosolic_FA-bd_dom domain-containing protein OS=Bos taurus OX=9913 GN=C8G PE=3 S | A0A3Q1MRQ2 [2] | Complement | 41 kDa | 10.83 | 0 | 0.00% | 7 | 0.14% | 0 | 0.00% | 0 | 0.00% | 0 | 0.00% |
| Cluster of L-lactate dehydrogenase OS=Bos taurus OX=9913 GN=LDHB PE=1 SV=1 (A0A3Q1M5R4) | A0A3Q1M5R4 [3] | Enzyme/Protease | 37 kDa | 5.86 | 9 | 0.25% | 12 | 0.24% | 11 | 0.20% | 8 | 0.21% | 10 | 0.27% |
| Cluster of L-lactate dehydrogenase OS=Bos taurus OX=9913 PE=3 SV=1 (A0A3Q1LY19) | A0A3Q1LY19 [4] | Enzyme/Protease | 37 kDa | 7.64 | 6 | 0.17% | 9 | 0.18% | 10 | 0.18% | 7 | 0.18% | 9 | 0.24% |
| Cluster of Moesin OS=Bos taurus OX=9913 GN=MSN PE=2 SV=3 (Q2HJ49) | Q2HJ49 [3] | Structual/Cytoskeleton | 68 kDa | 5.9 | 0 | 0.00% | 7 | 0.14% | 8 | 0.14% | 3 | 0.08% | 3 | 0.08% |
| Cluster of Myosin heavy chain 9 OS=Bos taurus OX=9913 GN=MYH9 PE=1 SV=3 (F1MQ37) | F1MQ37 [3] | Structual/Cytoskeleton | 227 kDa | 5.48 | 0 | 0.00% | 3 | 0.06% | 0 | 0.00% | 3 | 0.08% | 2 | 0.05% |
| Cluster of Periostin OS=Bos taurus OX=9913 GN=POSTN PE=1 SV=1 (A0A3Q1MF21) | A0A3Q1MF21 [4] | Cell-Cell Adhesion | 90 kDa | 7.33 | 40 | 1.10% | 66 | 1.32% | 69 | 1.25% | 53 | 1.37% | 38 | 1.02% |
| Cluster of Pyruvate kinase OS=Bos taurus OX=9913 GN=PKM PE=1 SV=1 (A5D984) | A5D984 [2] | Enzyme/Protease | 58 kDa | 7.96 | 2 | 0.06% | 11 | 0.22% | 10 | 0.18% | 2 | 0.05% | 3 | 0.08% |
| Cluster of Rab GDP dissociation inhibitor beta OS=Bos taurus OX=9913 GN=GDI2 PE=2 SV=3 (P50397) | P50397 [2] | Cell-Cell Adhesion | 50 kDa | 5.93 | 0 | 0.00% | 0 | 0.00% | 3 | 0.05% | 0 | 0.00% | 0 | 0.00% |
| Cluster of Regucalcin OS=Bos taurus OX=9913 GN=RGN PE=2 SV=1 (Q9TTJ5) | Q9TTJ5 [2] | Enzyme/Protease | 33 kDa | 5.54 | 18 | 0.50% | 20 | 0.40% | 19 | 0.34% | 16 | 0.42% | 17 | 0.46% |
| Cluster of Serpin A3-4 OS=Bos taurus OX=9913 GN=SERPINA3-4 PE=3 SV=1 (A2I7N0) | A2I7N0 [3] | Protease Inhibitor | 46 kDa | 5.62 | 0 | 0.00% | 3 | 0.06% | 4 | 0.07% | 0 | 0.00% | 0 | 0.00% |
| Cluster of SERPIN domain-containing protein OS=Bos taurus OX=9913 GN=LOC784932 PE=1 SV=1 (A0A0A( | A0A0A0MPA0 [3] | Protease Inhibitor | 47 kDa | 5.44 | 3 | 0.08% | 6 | 0.12% | 6 | 0.11% | 0 | 0.00% | 3 | 0.08% |

| | | | | | | | | | | | | | | |
|---|---|---|---|---|---|---|---|---|---|---|---|---|---|---|
| Cluster of SERPINA10 protein OS=Bos taurus OX=9913 GN=SERPINA10 PE=2 SV=1 (A5PJ69) | A5PJ69 [2] | Protease Inhibitor | 52 kDa | 6.05 | 0 | 0.00% | 0 | 0.00% | 2 | 0.04% | 0 | 0.00% | 0 | 0.00% |
| Cluster of Talin 1 OS=Bos taurus OX=9913 GN=TLN1 PE=1 SV=1 (A0A3Q1MLQ7) | A0A3Q1MLQ7 [6] | Cell-Cell Adhesion | 271 kDa | 5.79 | 4 | 0.11% | 28 | 0.56% | 33 | 0.60% | 20 | 0.52% | 24 | 0.64% |
| Cluster of Thyroglobulin OS=Bos taurus OX=9913 GN=TG PE=3 SV=1 (A0A3Q1M6P2) | A0A3Q1M6P2 [3] | Cell-Cell Adhesion | 303 kDa | 5.48 | 0 | 0.00% | 0 | 0.00% | 0 | 0.00% | 5 | 0.13% | 11 | 0.29% |
| Cluster of Transferrin receptor protein 1 OS=Bos taurus OX=9913 GN=TFRC PE=3 SV=3 (E1BIG6) | E1BIG6 [2] | Transport/Carrier/Binding | 85 kDa | 5.95 | 0 | 0.00% | 11 | 0.22% | 11 | 0.20% | 8 | 0.21% | 10 | 0.27% |
| Cluster of Tubulin alpha chain OS=Bos taurus OX=9913 GN=TUBA4A PE=3 SV=1 (A0A452DIH7) | A0A452DIH7 [6] | Structual/Cytoskeleton | 49 kDa | 5.91 | 0 | 0.00% | 17 | 0.34% | 16 | 0.29% | 10 | 0.26% | 5 | 0.13% |
| Cluster of Uncharacterized protein OS=Bos taurus OX=9913 PE=1 SV=3 (F1MZ96) | F1MZ96 [3] | Immunity/Immunity Related | 27 kDa | 5.94 | 10 | 0.28% | 23 | 0.46% | 21 | 0.38% | 10 | 0.26% | 9 | 0.24% |
| Cluster of Vascular cell adhesion molecule-1 7D variant OS=Bos taurus OX=9913 GN=VCAM1 PE=2 SV=1 (Q | Q9GKR3 [2] | Cell-Cell Adhesion | 81 kDa | 5.24 | 6 | 0.17% | 16 | 0.32% | 11 | 0.20% | 10 | 0.26% | 8 | 0.21% |
| Cluster of Vitamin K-dependent protein S OS=Bos taurus OX=9913 GN=PROS1 PE=1 SV=1 (P07224) | P07224 [2] | Coagulation | 75 kDa | 5.01 | 9 | 0.25% | 3 | 0.06% | 0 | 0.00% | 0 | 0.00% | 0 | 0.00% |
| Coagulation factor XI OS=Bos taurus OX=9913 GN=F11 PE=4 SV=1 | F1MUT4 (+1) | Coagulation | 70 kDa | 7.85 | 16 | 0.44% | 23 | 0.46% | 23 | 0.42% | 18 | 0.47% | 20 | 0.54% |
| Coagulation factor XII OS=Bos taurus OX=9913 GN=F12 PE=4 SV=2 | F1MTT3 (+1) | Coagulation | 66 kDa | 7.61 | 5 | 0.14% | 0 | 0.00% | 5 | 0.09% | 0 | 0.00% | 0 | 0.00% |
| Coagulation factor XIII B chain OS=Bos taurus OX=9913 GN=F13B PE=4 SV=1 | A0A3Q1MWQ1 | Coagulation | 68 kDa | 6.24 | 0 | 0.00% | 3 | 0.06% | 6 | 0.11% | 6 | 0.16% | 8 | 0.21% |
| Collagen alpha-1(I) chain OS=Bos taurus OX=9913 GN=COL1A1 PE=1 SV=3 | P02453 | Extracellular Matrix | 139 kDa | 9.28 | 13 | 0.36% | 16 | 0.32% | 21 | 0.38% | 18 | 0.47% | 17 | 0.46% |
| Collagen alpha-1(II) chain OS=Bos taurus OX=9913 GN=COL2A1 PE=1 SV=4 | P02459 | Extracellular Matrix | 142 kDa | 8.38 | 0 | 0.00% | 0 | 0.00% | 5 | 0.09% | 0 | 0.00% | 0 | 0.00% |
| Collagen alpha-1(XII) chain OS=Bos taurus OX=9913 GN=COL12A1 PE=4 SV=3 | F1N401 | Extracellular Matrix | 333 kDa | 5.35 | 0 | 0.00% | 14 | 0.28% | 8 | 0.14% | 4 | 0.10% | 0 | 0.00% |
| Collagen alpha-2(I) chain OS=Bos taurus OX=9913 GN=COL1A2 PE=1 SV=2 | P02465 | Extracellular Matrix | 129 kDa | 9.93 | 16 | 0.44% | 25 | 0.50% | 21 | 0.38% | 20 | 0.52% | 15 | 0.40% |
| Collagen type V alpha 1 chain OS=Bos taurus OX=9913 GN=COL5A1 PE=4 SV=2 | G3MZI7 | Extracellular Matrix | 185 kDa | 4.84 | 4 | 0.11% | 3 | 0.06% | 3 | 0.05% | 0 | 0.00% | 3 | 0.08% |
| Collectin-11 OS=Bos taurus OX=9913 GN=COLEC11 PE=4 SV=1 | A0A3Q1M944 (+2) | Immunity/Immunity Related | 29 kDa | 5.62 | 0 | 0.00% | 3 | 0.06% | 2 | 0.04% | 0 | 0.00% | 0 | 0.00% |
| Complement C1q subcomponent subunit A OS=Bos taurus OX=9913 GN=C1QA PE=2 SV=1 | Q5E9E3 | Complement | 26 kDa | 9.12 | 0 | 0.00% | 6 | 0.12% | 6 | 0.11% | 5 | 0.13% | 4 | 0.11% |
| Complement C1q subcomponent subunit B OS=Bos taurus OX=9913 GN=C1QB PE=1 SV=1 | Q2KIV9 | Complement | 26 kDa | 9.26 | 0 | 0.00% | 4 | 0.08% | 4 | 0.07% | 0 | 0.00% | 0 | 0.00% |
| Complement C8 alpha chain OS=Bos taurus OX=9913 GN=C8A PE=3 SV=1 | A0A3Q1LSP4 (+1) | Complement | 65 kDa | 6.51 | 0 | 0.00% | 0 | 0.00% | 2 | 0.04% | 2 | 0.05% | 2 | 0.05% |
| Complement subcomponent C1r OS=Bos taurus OX=9913 GN=C1R PE=4 SV=1 | A0A3Q1MGP1 (+1) | Complement | 81 kDa | 5.8 | 3 | 0.08% | 7 | 0.14% | 7 | 0.13% | 6 | 0.16% | 8 | 0.21% |
| Contactin-1 OS=Bos taurus OX=9913 GN=CNTN1 PE=3 SV=1 | A0A3Q1M7H6 (+1) | Extracellular Matrix | 113 kDa | 5.75 | 24 | 0.66% | 42 | 0.84% | 41 | 0.74% | 32 | 0.83% | 31 | 0.83% |
| CPN2 protein OS=Bos taurus OX=9913 GN=CPN2 PE=2 SV=1 | A6QP30 | Cell-Cell Adhesion | 60 kDa | 5.66 | 4 | 0.11% | 5 | 0.10% | 6 | 0.11% | 3 | 0.08% | 4 | 0.11% |
| C-type lectin domain containing 11A OS=Bos taurus OX=9913 GN=CLEC11A PE=2 SV=1 | A5D7L1 | Cell-Cell Adhesion | 36 kDa | 5.38 | 2 | 0.06% | 2 | 0.04% | 0 | 0.00% | 0 | 0.00% | 0 | 0.00% |
| Cystatin E/M OS=Bos taurus OX=9913 GN=CST6 PE=4 SV=1 | Q5DPW9 | Protease Inhibitor | 16 kDa | 6.31 | 3 | 0.08% | 0 | 0.00% | 0 | 0.00% | 0 | 0.00% | 0 | 0.00% |
| Desmoglein-1 OS=Bos taurus OX=9913 GN=DSG1 PE=4 SV=2 | F1MIW8 | Cell-Cell Adhesion | 112 kDa | 4.88 | 3 | 0.08% | 0 | 0.00% | 0 | 0.00% | 0 | 0.00% | 0 | 0.00% |
| Dihydropteridine reductase OS=Bos taurus OX=9913 GN=QDPR PE=2 SV=1 | Q3T0Z7 | Enzyme/Protease | 26 kDa | 6.9 | 2 | 0.06% | 0 | 0.00% | 0 | 0.00% | 0 | 0.00% | 0 | 0.00% |
| EGF containing fibulin extracellular matrix protein 1 OS=Bos taurus OX=9913 GN=EFEMP1 PE=1 SV=1 | A2VE41 | Extracellular Matrix | 55 kDa | 4.81 | 3 | 0.08% | 5 | 0.10% | 7 | 0.13% | 5 | 0.13% | 6 | 0.16% |
| Extracellular matrix protein 1 OS=Bos taurus OX=9913 GN=ECM1 PE=4 SV=1 | A0A3Q1M5Q6 (+1) | Extracellular Matrix | 63 kDa | 6.6 | 0 | 0.00% | 6 | 0.12% | 4 | 0.07% | 4 | 0.10% | 0 | 0.00% |
| Fatty acid-binding protein, liver OS=Bos taurus OX=9913 GN=FABP1 PE=1 SV=1 | P80425 | Transport/Carrier/Binding | 14 kDa | 7.78 | 7 | 0.19% | 9 | 0.18% | 7 | 0.13% | 3 | 0.08% | 5 | 0.13% |
| Fermitin family homolog 3 OS=Bos taurus OX=9913 GN=FERMT3 PE=3 SV=3 | F1MMJ5 | Cell-Cell Adhesion | 75 kDa | 6.26 | 0 | 0.00% | 3 | 0.06% | 2 | 0.04% | 0 | 0.00% | 0 | 0.00% |
| Fibulin-1 OS=Bos taurus OX=9913 GN=FBLN1 PE=1 SV=1 | A0A3Q1MJB2 (+1) | Coagulation | 77 kDa | 4.9 | 39 | 1.07% | 51 | 1.02% | 63 | 1.14% | 41 | 1.06% | 37 | 0.99% |
| Fructose-bisphosphate aldolase OS=Bos taurus OX=9913 GN=ALDOA PE=1 SV=1 | A6QLL8 | Enzyme/Protease | 39 kDa | 8.75 | 3 | 0.08% | 14 | 0.28% | 17 | 0.31% | 10 | 0.26% | 14 | 0.37% |
| Fumarylacetoacetase OS=Bos taurus OX=9913 GN=FAH PE=2 SV=1 | A5PKH3 (+1) | Enzyme/Protease | 46 kDa | 6.49 | 0 | 0.00% | 5 | 0.10% | 7 | 0.13% | 3 | 0.08% | 4 | 0.11% |
| Galactokinase OS=Bos taurus OX=9913 GN=GALK1 PE=2 SV=2 | A6H768 | Enzyme/Protease | 42 kDa | 5.77 | 0 | 0.00% | 3 | 0.06% | 0 | 0.00% | 0 | 0.00% | 0 | 0.00% |
| Galectin-3-binding protein OS=Bos taurus OX=9913 GN=LGALS3BP PE=1 SV=1 | A7E3W2 | Immunity/Immunity Related | 62 kDa | 5.25 | 5 | 0.14% | 14 | 0.28% | 16 | 0.29% | 7 | 0.18% | 4 | 0.11% |
| Gamma-glutamyl hydrolase OS=Bos taurus OX=9913 GN=GGH PE=2 SV=1 | A7YWG4 | Enzyme/Protease | 36 kDa | 8.96 | 0 | 0.00% | 0 | 0.00% | 2 | 0.04% | 0 | 0.00% | 0 | 0.00% |
| Gc-globulin OS=Bos taurus OX=9913 GN=GC PE=4 SV=2 | F1N5M2 | Transport/Carrier/Binding | 53 kDa | 5.24 | 6 | 0.17% | 10 | 0.20% | 10 | 0.18% | 8 | 0.21% | 8 | 0.21% |
| Glucose-6-phosphate isomerase OS=Bos taurus OX=9913 GN=GPI PE=2 SV=4 | Q3ZBD7 | Enzyme/Protease | 63 kDa | 7.45 | 0 | 0.00% | 0 | 0.00% | 3 | 0.05% | 0 | 0.00% | 0 | 0.00% |
| GTP-binding nuclear protein Ran OS=Bos taurus OX=9913 PE=3 SV=2 | G3MZP2 (+1) | Transport/Carrier/Binding | 24 kDa | 7.74 | 2 | 0.06% | 3 | 0.06% | 0 | 0.00% | 0 | 0.00% | 3 | 0.08% |
| Haptoglobin OS=Bos taurus OX=9913 GN=HP PE=3 SV=1 | A0A3Q1MB98 (+1) | Complement | 44 kDa | 8.09 | 0 | 0.00% | 4 | 0.08% | 4 | 0.07% | 0 | 0.00% | 2 | 0.05% |
| Hemopexin OS=Bos taurus OX=9913 GN=HPX PE=4 SV=1 | A0A452DI25 | Transport/Carrier/Binding | 52 kDa | 6.89 | 7 | 0.19% | 7 | 0.14% | 10 | 0.18% | 7 | 0.18% | 8 | 0.21% |
| Heparan sulfate proteoglycan 2 OS=Bos taurus OX=9913 GN=HSPG2 PE=1 SV=2 | F1MER7 | Coagulation | 464 kDa | 6.01 | 0 | 0.00% | 0 | 0.00% | 2 | 0.04% | 0 | 0.00% | 7 | 0.19% |
| Hepatocyte growth factor-like protein OS=Bos taurus OX=9913 GN=MST1 PE=2 SV=1 | Q24K22 | Enzyme/Protease | 80 kDa | 8.42 | 0 | 0.00% | 6 | 0.12% | 11 | 0.20% | 8 | 0.21% | 9 | 0.24% |
| HGF activator OS=Bos taurus OX=9913 GN=HGFAC PE=4 SV=3 | E1BCW0 | Enzyme/Protease | 70 kDa | 7.66 | 13 | 0.36% | 13 | 0.26% | 14 | 0.25% | 8 | 0.21% | 9 | 0.24% |
| Homogentisate 1,2-dioxygenase OS=Bos taurus OX=9913 GN=HGD PE=1 SV=1 | A0A3Q1N7G4 | Enzyme/Protease | 50 kDa | 6.86 | 0 | 0.00% | 0 | 0.00% | 6 | 0.11% | 5 | 0.13% | 0 | 0.00% |
| Ig-like domain-containing protein OS=Bos taurus OX=9913 PE=4 SV=1 | A0A3Q1MN72 (+2) | Immunoglobulin | 14 kDa | 7.71 | 0 | 0.00% | 3 | 0.06% | 0 | 0.00% | 0 | 0.00% | 0 | 0.00% |
| Ig-like domain-containing protein OS=Bos taurus OX=9913 PE=4 SV=1 | A0A3Q1LLT0 (+1) | Immunoglobulin | 24 kDa | 9.39 | 0 | 0.00% | 6 | 0.12% | 6 | 0.11% | 0 | 0.00% | 0 | 0.00% |
| Ig-like domain-containing protein OS=Bos taurus OX=9913 PE=4 SV=2 | G3MXG6 | Immunoglobulin | 15 kDa | 6.53 | 0 | 0.00% | 6 | 0.12% | 4 | 0.07% | 0 | 0.00% | 0 | 0.00% |
| Insulin-like growth factor binding protein acid labile subunit OS=Bos taurus OX=9913 GN=IGFALS PE=2 SV=1 | Q09TE3 | Cell-Cell Adhesion | 66 kDa | 6.73 | 9 | 0.25% | 9 | 0.18% | 12 | 0.22% | 8 | 0.21% | 6 | 0.16% |
| Insulin-like growth factor-binding protein 2 OS=Bos taurus OX=9913 GN=IGFBP2 PE=1 SV=2 | P13384 | Protease Inhibitor | 34 kDa | 6.93 | 3 | 0.08% | 3 | 0.06% | 4 | 0.07% | 0 | 0.00% | 0 | 0.00% |
| Junction plakoglobin OS=Bos taurus OX=9913 GN=JUP PE=1 SV=1 | A0A3Q1M1M7 (+1) | Cell-Cell Adhesion | 81 kDa | 5.95 | 19 | 0.52% | 0 | 0.00% | 0 | 0.00% | 0 | 0.00% | 0 | 0.00% |
| LDL receptor related protein 1 OS=Bos taurus OX=9913 GN=LRP1 PE=1 SV=3 | E1BGJ0 | Extracellular Matrix | 505 kDa | 5.12 | 0 | 0.00% | 0 | 0.00% | 0 | 0.00% | 3 | 0.08% | 3 | 0.08% |
| Lumican OS=Bos taurus OX=9913 GN=LUM PE=1 SV=1 | Q05443 | Extracellular Matrix | 39 kDa | 5.93 | 7 | 0.19% | 14 | 0.28% | 12 | 0.22% | 6 | 0.16% | 5 | 0.13% |
| Malic enzyme OS=Bos taurus OX=9913 GN=ME1 PE=3 SV=3 | F1N3V0 | Enzyme/Protease | 64 kDa | 6.14 | 0 | 0.00% | 0 | 0.00% | 5 | 0.09% | 0 | 0.00% | 3 | 0.08% |
| Mannan binding lectin serine peptidase 2 OS=Bos taurus OX=9913 GN=MASP2 PE=4 SV=3 | E1BJ49 | Complement | 76 kDa | 5.75 | 3 | 0.08% | 0 | 0.00% | 0 | 0.00% | 0 | 0.00% | 0 | 0.00% |
| Mannose receptor C type 2 OS=Bos taurus OX=9913 GN=MRC2 PE=4 SV=1 | A0A3Q1M5H7 | Cell-Cell Adhesion | 167 kDa | 5.62 | 5 | 0.14% | 7 | 0.14% | 9 | 0.16% | 5 | 0.13% | 4 | 0.11% |
| Mannose-binding protein C OS=Bos taurus OX=9913 GN=MBL PE=2 SV=1 | O02659 | Complement | 26 kDa | 5.11 | 0 | 0.00% | 7 | 0.14% | 5 | 0.09% | 3 | 0.08% | 2 | 0.05% |
| Metavinculin OS=Bos taurus OX=9913 GN=VCL PE=1 SV=1 | A0A3Q1MN97 (+1) | Cell-Cell Adhesion | 117 kDa | 5.83 | 56 | 1.54% | 79 | 1.58% | 62 | 1.12% | 60 | 1.56% | 55 | 1.47% |
| Myocilin OS=Bos taurus OX=9913 GN=MYOC PE=2 SV=1 | Q9XTA3 | Structual/Cytoskeleton | 55 kDa | 5.47 | 3 | 0.08% | 0 | 0.00% | 0 | 0.00% | 0 | 0.00% | 0 | 0.00% |
| Neogenin 1 OS=Bos taurus OX=9913 GN=NEO1 PE=3 SV=1 | A0A3Q1MFP4 | Cell-Cell Adhesion | 154 kDa | 6.03 | 6 | 0.17% | 13 | 0.26% | 15 | 0.27% | 7 | 0.18% | 8 | 0.21% |
| NID1 protein OS=Bos taurus OX=9913 GN=NID1 PE=2 SV=1 | A6QNS6 | Cell-Cell Adhesion | 136 kDa | 5.24 | 3 | 0.08% | 6 | 0.12% | 3 | 0.05% | 0 | 0.00% | 0 | 0.00% |
| Osteomodulin OS=Bos taurus OX=9913 GN=OMD PE=4 SV=1 | G3X6Y4 | Structual/Cytoskeleton | 49 kDa | 5.21 | 0 | 0.00% | 4 | 0.08% | 0 | 0.00% | 0 | 0.00% | 0 | 0.00% |
| Parkinson disease protein 7 homolog OS=Bos taurus OX=9913 GN=PARK7 PE=2 SV=1 | Q5E946 | N/A | 20 kDa | 7.05 | 2 | 0.06% | 3 | 0.06% | 4 | 0.07% | 0 | 0.00% | 3 | 0.08% |
| Pentraxin-related protein PTX3 OS=Bos taurus OX=9913 GN=PTX3 PE=2 SV=1 | Q0VCG9 | Extracellular Matrix | 42 kDa | 5.02 | 0 | 0.00% | 4 | 0.08% | 3 | 0.05% | 0 | 0.00% | 0 | 0.00% |
| Phosphatidylinositol-glycan-specific phospholipase D OS=Bos taurus OX=9913 GN=GPLD1 PE=1 SV=1 | P80109 | Enzyme/Protease | 93 kDa | 6.06 | 7 | 0.19% | 10 | 0.20% | 8 | 0.14% | 5 | 0.13% | 4 | 0.11% |
| Phosphoglucomutase-1 OS=Bos taurus OX=9913 GN=PGM1 PE=3 SV=1 | A0A3Q1LRD1 (+1) | Enzyme/Protease | 62 kDa | 6.2 | 8 | 0.22% | 15 | 0.30% | 20 | 0.36% | 11 | 0.29% | 16 | 0.43% |
| Phospholipid transfer protein OS=Bos taurus OX=9913 GN=PLTP PE=2 SV=1 | Q58DL9 | Immunity/Immune Related | 56 kDa | 6.32 | 0 | 0.00% | 3 | 0.06% | 3 | 0.05% | 0 | 0.00% | 0 | 0.00% |
| Pigment epithelium-derived factor OS=Bos taurus OX=9913 GN=SERPINF1 PE=1 SV=1 | Q95121 | Coagulation | 46 kDa | 6.31 | 39 | 1.07% | 60 | 1.20% | 61 | 1.10% | 36 | 0.93% | 32 | 0.86% |
| Plasma kallikrein OS=Bos taurus OX=9913 GN=KLKB1 PE=2 SV=1 | Q2KJ63 | Enzyme/Protease | 71 kDa | 8.64 | 24 | 0.66% | 33 | 0.66% | 34 | 0.62% | 22 | 0.57% | 22 | 0.59% |
| Plasma serine protease inhibitor OS=Bos taurus OX=9913 GN=SERPINA5 PE=3 SV=1 | A0A452DHZ3 | Coagulation | 44 kDa | 9.36 | 24 | 0.66% | 39 | 0.78% | 31 | 0.56% | 12 | 0.31% | 5 | 0.13% |
| Procollagen C-endopeptidase enhancer OS=Bos taurus OX=9913 GN=PCOLCE PE=2 SV=1 | Q2HJB6 | Enzyme/Protease | 48 kDa | 8.15 | 7 | 0.19% | 5 | 0.10% | 0 | 0.00% | 0 | 0.00% | 0 | 0.00% |
| Prolyl endopeptidase FAP OS=Bos taurus OX=9913 GN=FAP PE=1 SV=1 | A5D7B7 | Enzyme/Protease | 88 kDa | 7.54 | 0 | 0.00% | 4 | 0.08% | 5 | 0.09% | 0 | 0.00% | 2 | 0.05% |

| | | | | | | | | | | | | | | |
|---|---|---|---|---|---|---|---|---|---|---|---|---|---|---|
| Protein HP-20 homolog OS=Bos taurus OX=9913 PE=2 SV=1 | Q2KIT0 | Cell-Cell Adhesion | 21 kDa | 8.86 | 8 | 0.22% | 13 | 0.26% | 10 | 0.18% | 7 | 0.18% | 4 | 0.11% |
| Protein HP-25 homolog 1 OS=Bos taurus OX=9913 PE=1 SV=1 | Q2KIX7 | Cell-Cell Adhesion | 23 kDa | 5.71 | 0 | 0.00% | 0 | 0.00% | 3 | 0.05% | 0 | 0.00% | 0 | 0.00% |
| Protein HP-25 homolog 2 OS=Bos taurus OX=9913 PE=2 SV=1 | Q2KIU3 | Cell-Cell Adhesion | 23 kDa | 5.29 | 5 | 0.14% | 6 | 0.12% | 4 | 0.07% | 0 | 0.00% | 0 | 0.00% |
| Protein kinase C-binding protein NELL2 OS=Bos taurus OX=9913 GN=NELL2 PE=2 SV=1 | A6QR11 | Transport/Carrier/Binding | 91 kDa | 5.42 | 6 | 0.17% | 8 | 0.16% | 9 | 0.16% | 7 | 0.18% | 6 | 0.16% |
| Protein tyrosine kinase 7 (inactive) OS=Bos taurus OX=9913 GN=PTK7 PE=3 SV=2 | F1MWK8 | Enzyme/Protease | 118 kDa | 6.41 | 0 | 0.00% | 3 | 0.06% | 0 | 0.00% | 0 | 0.00% | 0 | 0.00% |
| Protein-tyrosine-phosphatase OS=Bos taurus OX=9913 GN=PTPRF PE=3 SV=1 | A0A3Q1M5Q9 (+2) | Enzyme/Protease | 213 kDa | 5.95 | 0 | 0.00% | 3 | 0.06% | 4 | 0.07% | 4 | 0.10% | 0 | 0.00% |
| Protein-tyrosine-phosphatase OS=Bos taurus OX=9913 GN=PTPRM PE=3 SV=1 | A0A3Q1MQL1 (+2) | Enzyme/Protease | 167 kDa | 6.13 | 0 | 0.00% | 7 | 0.14% | 5 | 0.09% | 3 | 0.08% | 5 | 0.13% |
| Proteoglycan 4 OS=Bos taurus OX=9913 GN=PRG4 PE=4 SV=1 | A0A3Q1LZ67 | Extracellular Matrix | 158 kDa | 8.91 | 0 | 0.00% | 0 | 0.00% | 5 | 0.09% | 0 | 0.00% | 0 | 0.00% |
| Prothrombin OS=Bos taurus OX=9913 GN=F2 PE=1 SV=2 | P00735 | Coagulation | 71 kDa | 5.4 | 15 | 0.41% | 15 | 0.30% | 14 | 0.25% | 7 | 0.18% | 7 | 0.19% |
| Receptor protein-tyrosine kinase OS=Bos taurus OX=9913 GN=TIE1 PE=4 SV=1 | A0A3Q1M0U5 | Cell-Cell Adhesion | 125 kDa | 6.33 | 0 | 0.00% | 2 | 0.04% | 0 | 0.00% | 0 | 0.00% | 0 | 0.00% |
| Serpin family G member 1 OS=Bos taurus OX=9913 GN=SERPING1 PE=3 SV=3 | E1BMJ0 | Complement | 52 kDa | 6.28 | 8 | 0.22% | 12 | 0.24% | 16 | 0.29% | 4 | 0.10% | 6 | 0.16% |
| Serpin H1 OS=Bos taurus OX=9913 GN=SERPINH1 PE=2 SV=1 | Q2KJH6 | Protease Inhibitor | 47 kDa | 8.82 | 0 | 0.00% | 2 | 0.04% | 3 | 0.05% | 0 | 0.00% | 0 | 0.00% |
| Serum amyloid P-component OS=Bos taurus OX=9913 GN=APCS PE=2 SV=1 | Q3T004 | Immunity/Immune Related | 25 kDa | 8.42 | 10 | 0.28% | 12 | 0.24% | 12 | 0.22% | 11 | 0.29% | 11 | 0.29% |
| Sulfhydryl oxidase OS=Bos taurus OX=9913 GN=QSOX1 PE=3 SV=1 | A0A3Q1N9Y5 (+1) | Enzyme/Protease | 86 kDa | 9.14 | 6 | 0.17% | 8 | 0.16% | 14 | 0.25% | 8 | 0.21% | 7 | 0.19% |
| Tetranectin OS=Bos taurus OX=9913 GN=CLEC3B PE=4 SV=1 | A0A452DIU4 | Cell-Cell Adhesion | 31 kDa | 6.52 | 10 | 0.28% | 9 | 0.18% | 7 | 0.13% | 0 | 0.00% | 0 | 0.00% |
| Threonine--tRNA ligase 1, cytoplasmic OS=Bos taurus OX=9913 GN=TARS1 PE=2 SV=1 | Q3ZBV8 | Enzyme/Protease | 83 kDa | 6.34 | 0 | 0.00% | 2 | 0.04% | 0 | 0.00% | 0 | 0.00% | 0 | 0.00% |
| Thrombospondin-4 OS=Bos taurus OX=9913 GN=THBS4 PE=3 SV=1 | A0A452DJ62 | Cell-Cell Adhesion | 106 kDa | 4.41 | 3 | 0.08% | 6 | 0.12% | 0 | 0.00% | 2 | 0.05% | 3 | 0.08% |
| Transforming growth factor beta receptor 3 OS=Bos taurus OX=9913 GN=TGFBR3 PE=4 SV=3 | E1B9H5 | Cell-Cell Adhesion | 95 kDa | 5.73 | 3 | 0.08% | 0 | 0.00% | 0 | 0.00% | 0 | 0.00% | 0 | 0.00% |
| Transforming growth factor-beta-induced protein ig-h3 OS=Bos taurus OX=9913 GN=TGFBI PE=1 SV=2 | P55906 | Cell-Cell Adhesion | 74 kDa | 6.69 | 0 | 0.00% | 11 | 0.22% | 8 | 0.14% | 0 | 0.00% | 0 | 0.00% |
| Transmembrane protein 132C OS=Bos taurus OX=9913 GN=TMEM132C PE=3 SV=2 | G3N019 | N/A | 120 kDa | 5.99 | 3 | 0.08% | 7 | 0.14% | 5 | 0.09% | 4 | 0.10% | 0 | 0.00% |
| Transthyretin OS=Bos taurus OX=9913 GN=TTR PE=1 SV=1 | O46375 | Enzyme/Protease | 16 kDa | 5.91 | 4 | 0.11% | 4 | 0.08% | 4 | 0.07% | 4 | 0.10% | 6 | 0.16% |
| Triosephosphate isomerase OS=Bos taurus OX=9913 GN=TPI1 PE=2 SV=3 | Q5E956 | Enzyme/Protease | 27 kDa | 6.51 | 11 | 0.30% | 20 | 0.40% | 18 | 0.33% | 13 | 0.34% | 12 | 0.32% |
| Uncharacterized protein OS=Bos taurus OX=9913 GN=HMCN2 PE=4 SV=3 | E1B9K4 | Cell-Cell Adhesion | 547 kDa | 5.45 | 0 | 0.00% | 4 | 0.08% | 0 | 0.00% | 0 | 0.00% | 2 | 0.05% |
| Uncharacterized protein OS=Bos taurus OX=9913 GN=LOC525947 PE=1 SV=1 | A0A3Q1NEQ0 | Transport/Carrier/Binding | 78 kDa | 6.8 | 39 | 1.07% | 52 | 1.04% | 72 | 1.30% | 38 | 0.99% | 39 | 1.04% |
| Uncharacterized protein OS=Bos taurus OX=9913 GN=LOC528040 PE=4 SV=3 | E1B805 | Complement | 186 kDa | 6.54 | 49 | 1.35% | 78 | 1.56% | 95 | 1.72% | 57 | 1.48% | 48 | 1.29% |
| Uncharacterized protein OS=Bos taurus OX=9913 GN=TNC PE=3 SV=1 | A0A3Q1LIW2 (+3) | Extracellular Matrix | 221 kDa | 4.94 | 2 | 0.06% | 4 | 0.08% | 7 | 0.13% | 3 | 0.08% | 2 | 0.05% |
| Uncharacterized protein OS=Bos taurus OX=9913 PE=1 SV=1 | G3N0V0 | Complement | 36 kDa | 8.05 | 0 | 0.00% | 5 | 0.10% | 7 | 0.13% | 0 | 0.00% | 0 | 0.00% |
| Uncharacterized protein OS=Bos taurus OX=9913 PE=4 SV=1 | A0A3Q1M0L5 | N/A | 23 kDa | 6.55 | 9 | 0.25% | 13 | 0.26% | 14 | 0.25% | 7 | 0.18% | 8 | 0.21% |
| VASN protein OS=Bos taurus OX=9913 GN=VASN PE=2 SV=1 | A4IFA5 | Cell-Cell Adhesion | 72 kDa | 8.63 | 7 | 0.19% | 9 | 0.18% | 7 | 0.13% | 6 | 0.16% | 5 | 0.13% |
| Vitamin K-dependent protein C OS=Bos taurus OX=9913 GN=PROC PE=4 SV=2 | A0A140T851 | Enzyme/Protease | 51 kDa | 6.06 | 3 | 0.08% | 0 | 0.00% | 0 | 0.00% | 0 | 0.00% | 0 | 0.00% |
| Vitamin K-dependent protein Z OS=Bos taurus OX=9913 GN=PROZ PE=4 SV=1 | A0A3Q1M8B7 (+2) | Enzyme/Protease | 53 kDa | 6.95 | 3 | 0.08% | 3 | 0.06% | 0 | 0.00% | 0 | 0.00% | 0 | 0.00% |

**Supplemental Table 3: Mass spectroscopy data for dendrimers incubated in 100% FBS**

| | | | | | G4-0% | | G4P8-25% | | G4P8-35% | | G4P8-45% | | 10% FBS (el |
|---|---|---|---|---|---|---|---|---|---|---|---|---|---|
| Identified Proteins | Accession Number | Biological Process | Molecular Weight | Theoretical pI | Total Spectrum Counts | Percentage of Total Spectral Counts | Total Spectrum Counts | Percentage of Total Spectral Counts | Total Spectrum Counts | Percentage of Total Spectral Counts | Total Spectrum Counts | Percentage of Total Spectral Counts | Total Spectrum Counts |
| Cluster of Apolipoprotein B OS=Bos taurus OX=9913 GN=APOB PE=1 SV=3 (E1BNR0) | E1BNR0 [2] | Apolipoprotein | 516 kDa | 6.24 | 718 | 15.30% | 932 | 16.25% | 678 | 14.92% | 847 | 15.15% | 906 |
| Cluster of C4a anaphylatoxin OS=Bos taurus OX=9913 GN=LOC107131209 PE=4 SV=3 (F1MVK1) | F1MVK1 [4] | Complement | 192 kDa | 7.26 | 350 | 7.46% | 370 | 6.45% | 315 | 5.49% | 428 | 7.65% | 449 |
| Cluster of Complement factor H OS=Bos taurus OX=9913 GN=CFH PE=1 SV=3 (Q28085) | Q28085 [4] | Complement | 140 kDa | 6.33 | 274 | 5.84% | 306 | 5.34% | 224 | 3.91% | 318 | 5.69% | 351 |
| Complement C3 OS=Bos taurus OX=9913 GN=C3 PE=1 SV=2 | Q2UVX4 | Complement | 187 kDa | 6.37 | 247 | 5.26% | 342 | 5.96% | 223 | 3.89% | 327 | 5.85% | 389 |
| Cluster of Plasminogen OS=Bos taurus OX=9913 GN=PLG PE=1 SV=2 (P06868) | P06868 [2] | Coagulation | 91 kDa | 7.39 | 245 | 5.22% | 310 | 5.41% | 196 | 3.42% | 287 | 5.13% | 335 |
| Cluster of Albumin OS=Bos taurus OX=9913 GN=ALB PE=4 SV=1 (A0A140T897) | A0A140T897 [2] | Transport/Carrier/Binding | 69 kDa | 5.77 | 130 | 2.77% | 190 | 3.31% | 122 | 2.13% | 202 | 3.61% | 211 |
| Alpha-2-macroglobulin OS=Bos taurus OX=9913 GN=A2M PE=1 SV=2 | Q7SIH1 | Protease Inhibitor | 168 kDa | 5.68 | 132 | 2.81% | 174 | 3.03% | 140 | 2.44% | 181 | 3.24% | 236 |
| Complement factor B OS=Bos taurus OX=9913 GN=CFB PE=1 SV=2 | P81187 | Complement | 85 kDa | 7.68 | 90 | 1.92% | 129 | 2.25% | 74 | 1.29% | 128 | 2.29% | 155 |
| Cluster of Serotransferrin OS=Bos taurus OX=9913 GN=TF PE=2 SV=1 (Q29443) | Q29443 [3] | Transport/Carrier/Binding | 78 kDa | 6.5 | 54 | 1.15% | 90 | 1.57% | 71 | 1.24% | 84 | 1.50% | 93 |
| Cluster of C4b-binding protein alpha chain OS=Bos taurus OX=9913 GN=C4BPA PE=2 SV=1 (Q28065) | Q28065 [3] | Complement | 69 kDa | 5.66 | 114 | 2.43% | 121 | 2.11% | 99 | 1.73% | 126 | 2.25% | 133 |
| Cluster of von Willebrand factor OS=Bos taurus OX=9913 GN=VWF PE=4 SV=1 (A0A3Q1LLU1) | A0A3Q1LLU1 [2] | Coagulation | 308 kDa | 5.35 | 91 | 1.94% | 105 | 1.83% | 102 | 1.78% | 121 | 2.16% | 115 |
| Cluster of Complement component C7 OS=Bos taurus OX=9913 GN=C7 PE=2 SV=1 (Q29RQ1) | Q29RQ1 [3] | Complement | 93 kDa | 6.9 | 70 | 1.49% | 93 | 1.62% | 70 | 1.22% | 102 | 1.82% | 129 |
| Cluster of Thrombospondin-1 OS=Bos taurus OX=9913 GN=THBS1 PE=3 SV=1 (F1N3A1) | F1N3A1 [2] | Cell-Cell Adhesion | 129 kDa | 4.7 | 74 | 1.58% | 107 | 1.87% | 92 | 1.60% | 80 | 1.43% | 107 |
| Cluster of Uncharacterized protein OS=Bos taurus OX=9913 GN=LOC506828 PE=3 SV=3 (F1MJK3) | F1MJK3 [4] | Protease Inhibitor | 163 kDa | 6.91 | 62 | 1.32% | 85 | 1.48% | 66 | 1.15% | 89 | 1.59% | 111 |
| Cluster of Uncharacterized protein OS=Bos taurus OX=9913 PE=1 SV=2 (G5E5T5) | G5E5T5 [3] | Immunity/Immunity Related | 56 kDa | 5.87 | 65 | 1.39% | 96 | 1.67% | 68 | 1.19% | 87 | 1.56% | 96 |
| Fibronectin OS=Bos taurus OX=9913 GN=FN1 PE=4 SV=1 | G5E5A8 | Structural/Cytoskeleton | 276 kDa | 5.79 | 67 | 1.43% | 87 | 1.52% | 66 | 1.15% | 91 | 1.63% | 95 |
| Cluster of Complement C5a anaphylatoxin OS=Bos taurus OX=9913 GN=C5 PE=1 SV=3 (F1MY85) | F1MY85 [2] | Complement | 189 kDa | 6.16 | 56 | 1.19% | 86 | 1.50% | 73 | 1.27% | 102 | 1.82% | 103 |
| Cluster of Trypsin OS=Sus scrofa OX=9823 (P00761) | P00761 [3] | Enzyme/Protease | 24 kDa | 8.26 | 39 | 0.83% | 64 | 1.12% | 37 | 0.65% | 45 | 0.80% | 42 |
| Collagen type VI alpha 3 chain OS=Bos taurus OX=9913 GN=COL6A3 PE=1 SV=2 | E1BB91 | Extracellular Matrix | 340 kDa | 5.87 | 57 | 1.21% | 60 | 1.05% | 67 | 1.17% | 75 | 1.34% | 57 |
| Coagulation factor V OS=Bos taurus OX=9913 GN=F5 PE=1 SV=1 | Q28107 | Coagulation | 249 kDa | 5.55 | 66 | 1.41% | 66 | 1.15% | 59 | 1.03% | 57 | 1.02% | 63 |
| Cluster of Fibrinogen beta chain OS=Bos taurus OX=9913 GN=FGB PE=4 SV=1 (A0A3Q1MG04) | A0A3Q1MG04 [3] | Coagulation | 57 kDa | 8.33 | 45 | 0.96% | 59 | 1.03% | 50 | 0.87% | 48 | 0.86% | 53 |
| Apolipoprotein E OS=Bos taurus OX=9913 GN=APOE PE=3 SV=1 | A0A3Q1LPF0 (+1) | Apolipoprotein | 36 kDa | 5.55 | 55 | 1.17% | 50 | 0.87% | 48 | 0.84% | 47 | 0.84% | 53 |
| Alpha-1-antiproteinase OS=Bos taurus OX=9913 GN=SERPINA1 PE=1 SV=1 | P34955 | Protease Inhibitor | 46 kDa | 5.98 | 23 | 0.49% | 36 | 0.63% | 29 | 0.51% | 39 | 0.70% | 45 |
| Uncharacterized protein OS=Bos taurus OX=9913 PE=4 SV=2 | G3N0S9 | Complement | 21 kDa | 5.37 | 41 | 0.87% | 61 | 1.06% | 38 | 0.66% | 52 | 0.93% | 59 |
| Cluster of Uncharacterized protein OS=Bos taurus OX=9913 PE=4 SV=3 (F1N160) | F1N160 [3] | Immunity/Immunity Related | 34 kDa | 8.86 | 45 | 0.96% | 39 | 0.68% | 40 | 0.70% | 40 | 0.72% | 38 |
| Complement C8 alpha chain OS=Bos taurus OX=9913 GN=C8A PE=3 SV=1 | A0A3Q1LSP4 (+1) | Complement | 65 kDa | 6.51 | 29 | 0.62% | 55 | 0.96% | 41 | 0.72% | 44 | 0.79% | 64 |
| Apolipoprotein H OS=Bos taurus OX=9913 GN=APOH PE=4 SV=1 | A0A140T843 (+1) | Apolipoprotein | 38 kDa | 8.55 | 36 | 0.77% | 45 | 0.78% | 25 | 0.44% | 42 | 0.75% | 57 |
| Inter-alpha-trypsin inhibitor heavy chain H2 OS=Bos taurus OX=9913 GN=ITIH2 PE=1 SV=1 | A0A3Q1LK49 (+1) | Protease Inhibitor | 97 kDa | 8.72 | 36 | 0.77% | 40 | 0.70% | 40 | 0.70% | 38 | 0.68% | 44 |
| Cluster of Mannan binding lectin serine peptidase 1 OS=Bos taurus OX=9913 GN=MASP1 PE=4 SV=1 (F1MVS9 | F1MVS9 [2] | Complement | 81 kDa | 4.93 | 42 | 0.89% | 45 | 0.78% | 45 | 0.78% | 45 | 0.80% | 55 |
| Fibrinogen gamma-B chain OS=Bos taurus OX=9913 GN=FGG PE=4 SV=1 | F1MGU7 (+1) | Coagulation | 50 kDa | 5.38 | 38 | 0.81% | 44 | 0.77% | 28 | 0.49% | 44 | 0.79% | 52 |
| Uncharacterized protein OS=Bos taurus OX=9913 PE=1 SV=1 | A0A3Q1M3L6 | Immunoglobulin | 40 kDa | 5.16 | 38 | 0.81% | 45 | 0.78% | 33 | 0.58% | 46 | 0.82% | 64 |
| Alpha-2-HS-glycoprotein OS=Bos taurus OX=9913 GN=AHSG PE=1 SV=2 | P12763 | Protease Inhibitor | 38 kDa | 5.1 | 25 | 0.53% | 21 | 0.37% | 22 | 0.38% | 22 | 0.39% | 30 |
| Fibrinogen alpha chain OS=Bos taurus OX=9913 GN=FGA PE=4 SV=1 | F6QND5 | Coagulation | 95 kDa | 5.69 | 38 | 0.81% | 41 | 0.72% | 32 | 0.56% | 40 | 0.72% | 42 |
| Cluster of Ig-like domain-containing protein OS=Bos taurus OX=9913 PE=4 SV=1 (A0A3Q1MIL5) | A0A3Q1MIL5 [12] | Immunoglobulin | 23 kDa | 6.97 | 34 | 0.72% | 47 | 0.82% | 35 | 0.61% | 49 | 0.88% | 51 |
| Cluster of Uncharacterized protein OS=Bos taurus OX=9913 GN=LOC790886 PE=4 SV=1 (A0A3Q1M315) | A0A3Q1M315 [2] | Complement | 45 kDa | 7.5 | 25 | 0.53% | 28 | 0.49% | 19 | 0.33% | 27 | 0.48% | 28 |
| Complement component C6 OS=Bos taurus OX=9913 GN=C6 PE=3 SV=2 | F1MM86 | Complement | 105 kDa | 6.74 | 23 | 0.49% | 34 | 0.59% | 30 | 0.52% | 40 | 0.72% | 41 |
| Cluster of Uncharacterized protein OS=Bos taurus OX=9913 PE=1 SV=1 (G3N0V0) | G3N0V0 [2] | Complement | 36 kDa | 8.05 | 23 | 0.49% | 39 | 0.68% | 25 | 0.44% | 38 | 0.68% | 44 |
| Pigment epithelium-derived factor OS=Bos taurus OX=9913 GN=SERPINF1 PE=1 SV=1 | Q95121 | Coagulation | 46 kDa | 6.31 | 23 | 0.49% | 35 | 0.61% | 24 | 0.42% | 36 | 0.64% | 38 |
| Fructose-bisphosphate aldolase OS=Bos taurus OX=9913 GN=ALDOA PE=1 SV=1 | A6QLL8 | Enzyme/Protease | 39 kDa | 8.45 | 28 | 0.60% | 27 | 0.47% | 28 | 0.49% | 41 | 0.73% | 45 |
| Complement component 8 subunit beta OS=Bos taurus OX=9913 GN=C8B PE=3 SV=3 | F1N102 | Complement | 67 kDa | 8.15 | 18 | 0.38% | 42 | 0.73% | 29 | 0.51% | 28 | 0.50% | 38 |
| Laminin subunit alpha 2 OS=Bos taurus OX=9913 GN=LAMA2 PE=4 SV=1 | A0A3Q1MFB8 (+2) | Extracellular Matrix | 224 kDa | 5.6 | 28 | 0.60% | 20 | 0.35% | 30 | 0.52% | 35 | 0.63% | 18 |
| Haptoglobin OS=Bos taurus OX=9913 GN=HP PE=3 SV=1 | A0A3Q1MB98 (+1) | Complement | 44 kDa | 8.09 | 18 | 0.38% | 32 | 0.56% | 28 | 0.49% | 27 | 0.48% | 21 |
| Uncharacterized protein OS=Bos taurus OX=9913 PE=1 SV=3 | F1MZ96 | Immunity/Immunity Related | 27 kDa | 5.94 | 30 | 0.64% | 28 | 0.49% | 26 | 0.45% | 23 | 0.41% | 21 |
| Glucose-6-phosphate isomerase OS=Bos taurus OX=9913 GN=GPI PE=2 SV=4 | Q3ZBD7 | Enzyme/Protease | 63 kDa | 7.45 | 23 | 0.49% | 27 | 0.47% | 23 | 0.40% | 29 | 0.52% | 29 |
| Galectin-3-binding protein OS=Bos taurus OX=9913 GN=LGALS3BP PE=1 SV=1 | A7E3W2 | Immunity/Immunity Related | 62 kDa | 5.25 | 26 | 0.55% | 30 | 0.52% | 34 | 0.59% | 23 | 0.41% | 26 |
| Cluster of Pyruvate kinase OS=Bos taurus OX=9913 GN=PKM PE=1 SV=1 (A5D984) | A5D984 [2] | Enzyme/Protease | 58 kDa | 7.96 | 19 | 0.40% | 26 | 0.45% | 15 | 0.26% | 26 | 0.46% | 27 |
| Apolipoprotein A-I OS=Bos taurus OX=9913 GN=APOA1 PE=1 SV=3 | P15497 | Apolipoprotein | 30 kDa | 5.36 | 20 | 0.43% | 24 | 0.42% | 23 | 0.40% | 22 | 0.39% | 27 |
| Cluster of Transferrin receptor protein 1 OS=Bos taurus OX=9913 GN=TFRC PE=3 SV=3 (E1BIG6) | E1BIG6 [3] | Transport/Carrier/Binding | 85 kDa | 5.95 | 16 | 0.34% | 23 | 0.40% | 23 | 0.40% | 23 | 0.41% | 13 |
| Coagulation factor XIII A chain OS=Bos taurus OX=9913 GN=F13A1 PE=3 SV=1 | A0A3Q1LTB9 (+1) | Coagulation | 76 kDa | 6 | 14 | 0.30% | 20 | 0.35% | 18 | 0.31% | 27 | 0.48% | 32 |
| Laminin subunit gamma 1 OS=Bos taurus OX=9913 GN=LAMC1 PE=1 SV=2 | F1MD77 | Extracellular Matrix | 178 kDa | 4.97 | 27 | 0.58% | 16 | 0.28% | 28 | 0.49% | 18 | 0.32% | 12 |
| Proteoglycan 4 OS=Bos taurus OX=9913 GN=PRG4 PE=4 SV=1 | A0A3Q1LZ67 | Extracellular Matrix | 158 kDa | 8.91 | 17 | 0.36% | 23 | 0.40% | 16 | 0.28% | 14 | 0.25% | 17 |
| Apolipoprotein D OS=Bos taurus OX=9913 GN=APOD PE=3 SV=3 | F1MS32 | Apolipoprotein | 24 kDa | 5.07 | 22 | 0.47% | 23 | 0.40% | 16 | 0.28% | 18 | 0.32% | 24 |
| Complement component C9 OS=Bos taurus OX=9913 GN=C9 PE=2 SV=1 | Q3MHN2 | Complement | 62 kDa | 5.57 | 19 | 0.40% | 22 | 0.38% | 17 | 0.30% | 19 | 0.34% | 17 |
| Cluster of Myosin heavy chain 9 OS=Bos taurus OX=9913 GN=MYH9 PE=1 SV=3 (F1MQ37) | F1MQ37 [5] | Structural/Cytoskeleton | 227 kDa | 5.48 | 24 | 0.51% | 7 | 0.12% | 24 | 0.42% | 16 | 0.29% | 7 |
| Cluster of Tubulin alpha chain OS=Bos taurus OX=9913 GN=TUBA4A PE=3 SV=1 (A0A3Q1M1Z2) | A0A3Q1M1Z2 [7] | Structural/Cytoskeleton | 46 kDa | 5.91 | 20 | 0.43% | 22 | 0.38% | 20 | 0.35% | 18 | 0.32% | 7 |
| Laminin subunit beta 1 OS=Bos taurus OX=9913 GN=LAMB1 PE=4 SV=1 | A0A3Q1MY30 | Extracellular Matrix | 219 kDa | 5.04 | 16 | 0.34% | 15 | 0.26% | 16 | 0.28% | 23 | 0.41% | 16 |
| Alpha-mannosidase OS=Bos taurus OX=9913 GN=MAN2B1 PE=3 SV=1 | A0A452DI46 (+1) | Enzyme/Protease | 111 kDa | 9.27 | 5 | 0.11% | 20 | 0.35% | 11 | 0.19% | 25 | 0.45% | 28 |
| Vitronectin OS=Bos taurus OX=9913 GN=VTN PE=1 SV=1 | Q3ZBS7 | Cell-Cell Adhesion | 54 kDa | 5.79 | 18 | 0.38% | 19 | 0.33% | 13 | 0.23% | 10 | 0.18% | 15 |
| Sushi, von Willebrand factor type A, EGF and pentraxin domain containing 1 OS=Bos taurus OX=9913 GN=SVE | F1MNH3 | Complement | 350 kDa | 5.25 | 22 | 0.47% | 22 | 0.38% | 17 | 0.30% | 13 | 0.23% | 10 |
| Inter-alpha-trypsin inhibitor heavy chain H3 OS=Bos taurus OX=9913 GN=ITIH3 PE=3 SV=1 | A0A3Q1LQ21 (+1) | Transport/Carrier/Binding | 99 kDa | 5.91 | 16 | 0.34% | 13 | 0.23% | 16 | 0.28% | 20 | 0.36% | 17 |
| Adiponectin M OS=Bos taurus OX=9913 GN=C1QTNF3 PE=2 SV=1 | A7MB82 | Other | 27 kDa | 5.91 | 4 | 0.09% | 10 | 0.17% | 10 | 0.17% | 30 | 0.54% | 23 |
| Reelin OS=Bos taurus OX=9913 GN=RELN PE=3 SV=3 | F1MUS6 | Extracellular Matrix | 388 kDa | 5.48 | 18 | 0.38% | 16 | 0.28% | 21 | 0.37% | 12 | 0.21% | 3 |
| Coagulation factor XIII B chain OS=Bos taurus OX=9913 GN=F13B PE=2 SV=1 | Q2TBQ1 | Coagulation | 75 kDa | 6.24 | 0 | 0.00% | 3 | 0.05% | 3 | 0.05% | 12 | 0.21% | 50 |
| Cluster of CD5 molecule like OS=Bos taurus OX=9913 GN=CD5L PE=2 SV=1 (A6QNW7) | A6QNW7 [2] | Immunity/Immunity Related | 50 kDa | 5.24 | 20 | 0.43% | 14 | 0.24% | 15 | 0.26% | 13 | 0.23% | 13 |
| Cluster of Tubulin beta chain OS=Bos taurus OX=9913 GN=TUBB1 PE=3 SV=1 (A0A3Q1M442) | A0A3Q1M442 [4] | Structural/Cytoskeleton | 55 kDa | 5.41 | 26 | 0.55% | 20 | 0.35% | 15 | 0.26% | 11 | 0.20% | 6 |
| Cluster of Hemoglobin fetal subunit beta OS=Bos taurus OX=9913 PE=1 SV=1 (P02081) | P02081 [2] | Transport/Carrier/Binding | 16 kDa | 6.51 | 13 | 0.28% | 18 | 0.31% | 13 | 0.23% | 15 | 0.27% | 19 |
| Uncharacterized protein OS=Bos taurus OX=9913 PE=1 SV=1 | A0A3Q1LPG0 | Immunity/Immunity Related | 36 kDa | 7.59 | 13 | 0.28% | 15 | 0.26% | 8 | 0.14% | 18 | 0.32% | 18 |
| Collagen alpha-1(II) chain OS=Bos taurus OX=9913 GN=COL2A1 PE=1 SV=4 | P02459 | Extracellular Matrix | 142 kDa | 8.38 | 13 | 0.28% | 17 | 0.30% | 11 | 0.19% | 11 | 0.20% | 19 |
| Apolipoprotein M OS=Bos taurus OX=9913 GN=APOM PE=3 SV=1 | A0A3Q1MB70 | Apolipoprotein | 25 kDa | 6.89 | 13 | 0.28% | 11 | 0.19% | 10 | 0.17% | 10 | 0.18% | 10 |
| Collagen type VI alpha 1 chain OS=Bos taurus OX=9913 GN=COL6A1 PE=1 SV=1 | E1BI98 | Extracellular Matrix | 109 kDa | 5.18 | 14 | 0.30% | 15 | 0.26% | 13 | 0.23% | 14 | 0.25% | 12 |
| Pentraxin-related protein PTX3 OS=Bos taurus OX=9913 GN=PTX3 PE=4 SV=1 | G3X6Q8 (+1) | Extracellular Matrix | 42 kDa | 5.02 | 11 | 0.23% | 19 | 0.33% | 13 | 0.23% | 12 | 0.21% | 20 |
| Complement factor properdin OS=Bos taurus OX=9913 GN=CFP PE=4 SV=1 | A0A3Q1M2R9 (+2) | Complement | 63 kDa | 8.51 | 17 | 0.36% | 9 | 0.16% | 7 | 0.12% | 10 | 0.18% | 13 |
| LDL receptor related protein 1 OS=Bos taurus OX=9913 GN=LRP1 PE=1 SV=3 | E1BGJ0 | Apolipoprotein | 505 kDa | 5.12 | 3 | 0.06% | 11 | 0.19% | 12 | 0.21% | 6 | 0.11% | 20 |
| Antithrombin-III OS=Bos taurus OX=9913 GN=SERPINC1 PE=3 SV=1 | A0A3Q1NJR8 | Coagulation | 60 kDa | 8.85 | 9 | 0.19% | 8 | 0.14% | 8 | 0.14% | 12 | 0.21% | 10 |
| Alpha-fetoprotein OS=Bos taurus OX=9913 GN=AFP PE=1 SV=1 | A0A3Q1MIW0 (+1) | Transport/Carrier/Binding | 68 kDa | 6.17 | 9 | 0.19% | 6 | 0.10% | 9 | 0.16% | 11 | 0.20% | 7 |

| | | | | | | | | | | | | | |
|---|---|---|---|---|---|---|---|---|---|---|---|---|---|
| Cluster of Ig-like domain-containing protein OS=Bos taurus OX=9913 PE=4 SV=1 (A0A3Q1LSF0) | A0A3Q1LSF0 [4] | Immunoglobulin | 15 kDa | 9.82 | 15 | 0.32% | 18 | 0.31% | 10 | 0.17% | 14 | 0.25% | 15 |
| Cluster of Gc-globulin OS=Bos taurus OX=9913 GN=GC PE=4 SV=2 (F1N5M2) | F1N5M2 [2] | Transport/Carrier/Binding | 53 kDa | 5.24 | 12 | 0.26% | 12 | 0.21% | 8 | 0.14% | 12 | 0.21% | 0 |
| Lipocln_cytosolic_FA-bd_dom domain-containing protein OS=Bos taurus OX=9913 GN=C8G PE=3 SV=1 | A0A3Q1MRQ2 | Complement | 41 kDa | 10.83 | 16 | 0.34% | 12 | 0.21% | 11 | 0.19% | 10 | 0.18% | 15 |
| Adiponectin B OS=Bos taurus OX=9913 GN=C1QC PE=4 SV=1 | A0A3B0IZF8 | Complement | 26 kDa | 8.62 | 18 | 0.38% | 15 | 0.26% | 12 | 0.21% | 12 | 0.21% | 10 |
| Mannan binding lectin serine peptidase 2 OS=Bos taurus OX=9913 GN=MASP2 PE=4 SV=3 | E1BJ49 | Complement | 76 kDa | 5.75 | 13 | 0.28% | 11 | 0.19% | 14 | 0.24% | 11 | 0.20% | 10 |
| Cluster of Collagen type VI alpha 2 chain OS=Bos taurus OX=9913 GN=COL6A2 PE=1 SV=3 (F1MKG2) | F1MKG2 [2] | Extracellular Matrix | 105 kDa | 6.2 | 13 | 0.28% | 10 | 0.17% | 17 | 0.30% | 10 | 0.18% | 7 |
| Fetuin-B OS=Bos taurus OX=9913 GN=FETUB PE=1 SV=1 | Q58D62 | Protease Inhibitor | 43 kDa | 5.59 | 5 | 0.11% | 4 | 0.07% | 8 | 0.14% | 5 | 0.09% | 3 |
| Cluster of Ig-like domain-containing protein OS=Bos taurus OX=9913 PE=1 SV=1 (A0A3Q1LUE9) | A0A3Q1LUE9 [4] | Immunoglobulin | 17 kDa | 8.76 | 10 | 0.21% | 12 | 0.21% | 7 | 0.12% | 12 | 0.21% | 8 |
| Ig-like domain-containing protein OS=Bos taurus OX=9913 PE=4 SV=1 | A0A3Q1MR79 | Immunoglobulin | 21 kDa | 10.28 | 11 | 0.23% | 11 | 0.19% | 7 | 0.12% | 11 | 0.20% | 14 |
| GLOBIN domain-containing protein OS=Bos taurus OX=9913 GN=HBA1 PE=3 SV=1 | A0A452DIQ5 (+1) | Transport/Carrier/Binding | 14 kDa | 7.07 | 5 | 0.11% | 13 | 0.23% | 5 | 0.09% | 10 | 0.18% | 8 |
| Complement C1q subcomponent subunit B OS=Bos taurus OX=9913 GN=C1QB PE=1 SV=1 | Q2KIV9 | Complement | 26 kDa | 9.26 | 4 | 0.09% | 14 | 0.24% | 8 | 0.14% | 11 | 0.20% | 12 |
| Cluster of Glutathione S-transferase A1 OS=Bos taurus OX=9913 GN=GSTA1 PE=2 SV=3 (Q28035) | Q28035 [10] | Enzyme/Protease | 25 kDa | 8.66 | 10 | 0.21% | 7 | 0.12% | 8 | 0.14% | 11 | 0.20% | 9 |
| Peptidoglycan recognition protein 1 OS=Bos taurus OX=9913 GN=PGLYRP1 PE=1 SV=1 | Q8SPP7 | Immunity/Immunity Related | 21 kDa | 9.38 | 10 | 0.21% | 11 | 0.19% | 10 | 0.17% | 13 | 0.23% | 12 |
| Platelet-activating factor acetylhydrolase OS=Bos taurus OX=9913 GN=PLA2G7 PE=2 SV=1 | Q1RML9 (+1) | Coagulation | 50 kDa | 5.96 | 8 | 0.17% | 10 | 0.17% | 7 | 0.12% | 14 | 0.25% | 9 |
| Cluster of Plasma serine protease inhibitor OS=Bos taurus OX=9913 GN=SERPINA5 PE=3 SV=1 (A0A452DHZ3 | A0A452DHZ3 [3] | Coagulation | 44 kDa | 9.36 | 6 | 0.13% | 11 | 0.19% | 8 | 0.14% | 11 | 0.20% | 12 |
| Extracellular matrix protein 1 OS=Bos taurus OX=9913 GN=ECM1 PE=4 SV=1 | A0A3Q1M5Q6 (+1) | Extracellular Matrix | 63 kDa | 6.6 | 9 | 0.19% | 10 | 0.17% | 9 | 0.16% | 8 | 0.14% | 12 |
| Cluster of Fructose-bisphosphate aldolase OS=Bos taurus OX=9913 GN=ALDOB PE=2 SV=1 (A5PK73) | A5PK73 [2] | Enzyme/Protease | 40 kDa | 8.69 | 0 | 0.00% | 0 | 0.00% | 11 | 0.19% | 10 | 0.18% | 10 |
| Adiponectin OS=Bos taurus OX=9913 GN=ADIPOQ PE=1 SV=1 | Q3Y5Z3 | Cell-Cell Adhesion | 26 kDa | 5.33 | 8 | 0.17% | 10 | 0.17% | 8 | 0.14% | 10 | 0.18% | 10 |
| Uncharacterized protein OS=Bos taurus OX=9913 GN=C8G PE=4 SV=1 | F6R7W8 | Complement | 47 kDa | 10.95 | 4 | 0.09% | 13 | 0.23% | 5 | 0.09% | 11 | 0.20% | 14 |
| Hemicentin 1 OS=Bos taurus OX=9913 GN=HMCN1 PE=4 SV=3 | E1BE11 | Extracellular Matrix | 613 kDa | 6.02 | 8 | 0.17% | 4 | 0.07% | 11 | 0.19% | 6 | 0.11% | 5 |
| Heparan sulfate proteoglycan 2 OS=Bos taurus OX=9913 GN=HSPG2 PE=1 SV=2 | F1MER7 | Coagulation | 464 kDa | 6.01 | 4 | 0.09% | 0 | 0.00% | 3 | 0.05% | 5 | 0.09% | 6 |
| Uncharacterized protein OS=Bos taurus OX=9913 PE=4 SV=1 | A0A3Q1MHP4 (+1) | Immunoglobulin | 101 kDa | 6.55 | 7 | 0.15% | 6 | 0.10% | 6 | 0.10% | 10 | 0.18% | 5 |
| Mannose-binding protein C OS=Bos taurus OX=9913 GN=MBL PE=2 SV=1 | O02659 | Complement | 26 kDa | 5.11 | 9 | 0.19% | 8 | 0.14% | 7 | 0.12% | 9 | 0.16% | 9 |
| Cluster of Clusterin OS=Bos taurus OX=9913 GN=CLU PE=1 SV=1 (P17697) | P17697 [2] | Apolipoprotein | 51 kDa | 5.72 | 8 | 0.17% | 7 | 0.12% | 8 | 0.14% | 0 | 0.00% | 0 |
| Adenylyl cyclase-associated protein OS=Bos taurus OX=9913 GN=CAP1 PE=2 SV=1 | A6QLB7 (+1) | Structural/Cytoskeleton | 51 kDa | 7.16 | 9 | 0.19% | 10 | 0.17% | 9 | 0.16% | 4 | 0.07% | 6 |
| Prenylcysteine oxidase 1 OS=Bos taurus OX=9913 GN=PCYOX1 PE=1 SV=2 | F1N2K1 | Enzyme/Protease | 57 kDa | 5.95 | 5 | 0.11% | 7 | 0.12% | 8 | 0.14% | 9 | 0.16% | 7 |
| Cluster of Thrombospondin-4 OS=Bos taurus OX=9913 GN=THBS4 PE=3 SV=1 (A0A452DJ62) | A0A452DJ62 [3] | Cell-Cell Adhesion | 106 kDa | 4.41 | 7 | 0.15% | 11 | 0.19% | 12 | 0.21% | 0 | 0.00% | 0 |
| Ig-like domain-containing protein OS=Bos taurus OX=9913 PE=4 SV=1 | A0A3Q1LI44 | Immunoglobulin | 15 kDa | 8.94 | 8 | 0.17% | 8 | 0.14% | 7 | 0.12% | 0 | 0.00% | 8 |
| Conglutinin OS=Bos taurus OX=9913 GN=CGN1 PE=1 SV=2 | P23805 | Extracellular Matrix | 38 kDa | 5.63 | 8 | 0.17% | 7 | 0.12% | 9 | 0.16% | 9 | 0.16% | 8 |
| Complement C1q subcomponent subunit A OS=Bos taurus OX=9913 GN=C1QA PE=2 SV=1 | Q5E9E3 | Complement | 26 kDa | 9.12 | 9 | 0.19% | 9 | 0.16% | 7 | 0.12% | 6 | 0.11% | 8 |
| Fructose-bisphosphate aldolase OS=Bos taurus OX=9913 GN=ALDOC PE=1 SV=1 | A0A3S5ZPB0 | Enzyme/Protease | 56 kDa | 8.75 | 5 | 0.11% | 2 | 0.03% | 8 | 0.14% | 11 | 0.20% | 7 |
| Alpha-1-acid glycoprotein OS=Bos taurus OX=9913 GN=ORM1 PE=2 SV=1 | Q3SZR3 (+1) | Transport/Carrier/Binding | 23 kDa | 5.67 | 3 | 0.06% | 0 | 0.00% | 3 | 0.05% | 0 | 0.00% | 0 |
| Cluster of L-lactate dehydrogenase OS=Bos taurus OX=9913 PE=3 SV=1 (A0A3Q1LY19) | A0A3Q1LY19 [4] | Enzyme/Protease | 37 kDa | 7.64 | 7 | 0.15% | 3 | 0.05% | 8 | 0.14% | 0 | 0.00% | 0 |
| Plasmalemma vesicle associated protein OS=Bos taurus OX=9913 GN=PLVAP PE=2 SV=1 | Q3ZC85 | Other | 50 kDa | 8.87 | 5 | 0.11% | 4 | 0.07% | 5 | 0.09% | 5 | 0.09% | 3 |
| Sorbitol dehydrogenase OS=Bos taurus OX=9913 GN=SORD PE=1 SV=3 | Q58D31 | Enzyme/Protease | 38 kDa | 7.14 | 4 | 0.09% | 0 | 0.00% | 4 | 0.07% | 4 | 0.07% | 9 |
| Immunoglobulin J chain OS=Bos taurus OX=9913 GN=JCHAIN PE=1 SV=1 | Q3SYR8 | Immunoglobulin | 18 kDa | 4.92 | 5 | 0.11% | 7 | 0.12% | 6 | 0.10% | 7 | 0.13% | 6 |
| TKT protein OS=Bos taurus OX=9913 GN=TKT PE=1 SV=1 | A7Z014 | Enzyme/Protease | 68 kDa | 7.2 | 7 | 0.15% | 0 | 0.00% | 9 | 0.16% | 8 | 0.14% | 5 |
| Kininogen-1 OS=Bos taurus OX=9913 GN=KNG1 PE=1 SV=1 | P01044 (+1) | Coagulation | 69 kDa | 6.05 | 7 | 0.15% | 0 | 0.00% | 6 | 0.10% | 0 | 0.00% | 2 |
| Aggrecan core protein OS=Bos taurus OX=9913 GN=ACAN PE=3 SV=2 | F1N367 (+2) | Cartilage Related | 243 kDa | 4.17 | 8 | 0.17% | 7 | 0.12% | 6 | 0.10% | 6 | 0.11% | 6 |
| Ig-like domain-containing protein OS=Bos taurus OX=9913 PE=4 SV=1 | A0A3Q1M3A8 | Immunoglobulin | 19 kDa | 5.89 | 4 | 0.09% | 7 | 0.12% | 6 | 0.10% | 7 | 0.13% | 7 |
| Alpha-1-microglobulin OS=Bos taurus OX=9913 GN=KIF12 PE=3 SV=3 | F1MMK9 | Other | 53 kDa | 8.87 | 10 | 0.21% | 4 | 0.07% | 9 | 0.16% | 3 | 0.05% | 3 |
| Cluster of Complement subcomponent C1r OS=Bos taurus OX=9913 GN=C1R PE=4 SV=1 (A0A3Q1MGP1) | A0A3Q1MGP1 [3] | Complement | 81 kDa | 5.8 | 5 | 0.11% | 7 | 0.12% | 5 | 0.09% | 0 | 0.00% | 6 |
| Cluster of Actin, cytoplasmic 2 OS=Bos taurus OX=9913 GN=ACTG1 PE=1 SV=1 (P63258) | P63258 [6] | Structural/Cytoskeleton | 42 kDa | 5.31 | 8 | 0.17% | 9 | 0.16% | 0 | 0.00% | 0 | 0.00% | 0 |
| Clathrin heavy chain 1 OS=Bos taurus OX=9913 GN=CLTC PE=1 SV=1 | P49951 | Cell-Cell Adhesion | 192 kDa | 5.48 | 6 | 0.13% | 3 | 0.05% | 6 | 0.10% | 2 | 0.04% | 2 |
| Cluster of Carboxypeptidase B2 OS=Bos taurus OX=9913 GN=CPB2 PE=1 SV=1 (Q2KIG3) | Q2KIG3 [2] | Enzyme/Protease | 49 kDa | 8.49 | 5 | 0.11% | 7 | 0.12% | 5 | 0.09% | 4 | 0.07% | 5 |
| Cluster of Phosphoglycerate kinase OS=Bos taurus OX=9913 GN=PGK1 PE=1 SV=2 (G3X7N4) | G3X7N4 [3] | Enzyme/Protease | 43 kDa | 8.3 | 4 | 0.09% | 5 | 0.09% | 2 | 0.03% | 0 | 0.00% | 4 |
| Ig-like domain-containing protein OS=Bos taurus OX=9913 PE=4 SV=2 | G3MXG6 | Immunoglobulin | 15 kDa | 8.56 | 6 | 0.13% | 6 | 0.10% | 6 | 0.10% | 6 | 0.11% | 6 |
| Cluster of Phosphatidylethanolamine-binding protein 1 OS=Bos taurus OX=9913 GN=PEBP1 PE=1 SV=2 (P136 | P13696 [2] | Protease Inhibitor | 21 kDa | 7.39 | 6 | 0.13% | 6 | 0.10% | 6 | 0.10% | 7 | 0.13% | 4 |
| CD9 antigen OS=Bos taurus OX=9913 GN=CD9 PE=2 SV=2 | P30932 | Other | 25 kDa | 6.31 | 0 | 0.00% | 3 | 0.05% | 4 | 0.07% | 0 | 0.00% | 3 |
| Transthyretin OS=Bos taurus OX=9913 GN=TTR PE=3 SV=1 | A0A3Q1MRM2 (+1) | Transport/Carrier/Binding | 20 kDa | 7.89 | 3 | 0.06% | 4 | 0.07% | 4 | 0.07% | 3 | 0.05% | 3 |
| Complement C1s subcomponent OS=Bos taurus OX=9913 GN=C1S PE=4 SV=1 | A0A3Q1M7Q5 (+2) | Complement | 78 kDa | 5.06 | 7 | 0.15% | 4 | 0.07% | 7 | 0.12% | 3 | 0.05% | 0 |
| Hemopexin OS=Bos taurus OX=9913 GN=HPX PE=4 SV=1 | A0A452DI25 (+1) | Transport/Carrier/Binding | 52 kDa | 6.89 | 0 | 0.00% | 5 | 0.09% | 7 | 0.12% | 5 | 0.09% | 4 |
| Uncharacterized protein OS=Bos taurus OX=9913 GN=CFHR5 PE=4 SV=1 | A0A3Q1M3I2 (+1) | Complement | 75 kDa | 8.57 | 5 | 0.11% | 4 | 0.07% | 5 | 0.09% | 5 | 0.09% | 5 |
| Alpha-2-antiplasmin OS=Bos taurus OX=9913 GN=SERPINF2 PE=1 SV=2 | P28800 | Protease Inhibitor | 55 kDa | 5.71 | 6 | 0.13% | 4 | 0.07% | 7 | 0.12% | 0 | 0.00% | 2 |
| Sulfhydryl oxidase OS=Bos taurus OX=9913 GN=QSOX1 PE=3 SV=1 | A0A3Q1N9Y5 (+1) | Enzyme/Protease | 86 kDa | 9.14 | 0 | 0.00% | 4 | 0.07% | 3 | 0.05% | 3 | 0.05% | 12 |
| Cluster of Uncharacterized protein OS=Bos taurus OX=9913 PE=4 SV=1 (A0A3Q1MA45) | A0A3Q1MA45 [2] | Enzyme/Protease | 47 kDa | 5.5 | 4 | 0.09% | 5 | 0.09% | 4 | 0.07% | 0 | 0.00% | 4 |
| Cluster of Heat shock cognate 71 kDa protein OS=Bos taurus OX=9913 GN=HSPA8 PE=1 SV=1 (A0A3Q1LMS5 | A0A3Q1LMS5 [6] | Other | 72 kDa | 5.37 | 5 | 0.11% | 4 | 0.07% | 3 | 0.05% | 0 | 0.00% | 0 |
| Serpin family D member 1 OS=Bos taurus OX=9913 GN=SERPIND1 PE=3 SV=1 | F6R4P6 | Coagulation | 63 kDa | 6.23 | 7 | 0.15% | 3 | 0.05% | 0 | 0.00% | 4 | 0.07% | 5 |
| Junction plakoglobin OS=Bos taurus OX=9913 GN=JUP PE=1 SV=1 | A0A3Q1M1M7 (+1) | Cell-Cell Adhesion | 81 kDa | 5.95 | 0 | 0.00% | 18 | 0.31% | 0 | 0.00% | 0 | 0.00% | 0 |
| Inter-alpha-trypsin inhibitor heavy chain H4 OS=Bos taurus OX=9913 GN=ITIH4 PE=3 SV=1 | A0A3Q1LZ09 (+3) | Protease Inhibitor | 101 kDa | 5.92 | 3 | 0.06% | 3 | 0.05% | 5 | 0.09% | 3 | 0.05% | 0 |
| Beta-2-microglobulin OS=Bos taurus OX=9913 PE=3 SV=1 | A0A3Q1MU93 (+1) | Immunity/Immunity Related | 14 kDa | 9.1 | 0 | 0.00% | 3 | 0.05% | 0 | 0.00% | 4 | 0.07% | 4 |
| CD59 glycoprotein OS=Bos taurus OX=9913 GN=CD59 PE=2 SV=1 | Q32PA1 | Complement | 14 kDa | 7.64 | 5 | 0.11% | 4 | 0.07% | 3 | 0.05% | 4 | 0.07% | 3 |
| FCNB protein OS=Bos taurus OX=9913 GN=FCNB PE=2 SV=1 | A4FV99 | Immunity/Immunity Related | 43 kDa | 8.46 | 6 | 0.13% | 3 | 0.05% | 7 | 0.12% | 3 | 0.05% | 0 |
| Ig-like domain-containing protein OS=Bos taurus OX=9913 PE=4 SV=1 | A0A3Q1LJT1 | Immunoglobulin | 25 kDa | 4.73 | 0 | 0.00% | 3 | 0.05% | 4 | 0.07% | 0 | 0.00% | 5 |
| Cluster of Peptidyl-prolyl cis-trans isomerase A OS=Bos taurus OX=9913 GN=PPIA PE=1 SV=2 (P62935) | P62935 [3] | Enzyme/Protease | 18 kDa | 8.34 | 0 | 0.00% | 0 | 0.00% | 0 | 0.00% | 0 | 0.00% | 4 |
| Symplekin OS=Bos taurus OX=9913 GN=SYMPK PE=4 SV=2 | E1BJF6 | Other | 141 kDa | 5.7 | 0 | 0.00% | 0 | 0.00% | 3 | 0.05% | 0 | 0.00% | 0 |
| Filamin A OS=Bos taurus OX=9913 GN=FLNA PE=1 SV=1 | A0A3Q1LMA3 (+2) | Structural/Cytoskeleton | 271 kDa | 5.63 | 0 | 0.00% | 0 | 0.00% | 5 | 0.09% | 0 | 0.00% | 3 |
| Collagen alpha-1(X) chain OS=Bos taurus OX=9913 GN=COL10A1 PE=4 SV=3 | F1MZ30 (+1) | Extracellular Matrix | 66 kDa | 9.59 | 4 | 0.09% | 4 | 0.07% | 0 | 0.00% | 4 | 0.07% | 4 |
| NID1 protein OS=Bos taurus OX=9913 GN=NID1 PE=2 SV=1 | A6QNS6 | Cell-Cell Adhesion | 136 kDa | 5.24 | 3 | 0.06% | 3 | 0.05% | 3 | 0.05% | 0 | 0.00% | 0 |
| Cluster of BPTI/Kunitz inhibitor domain-containing protein OS=Bos taurus OX=9913 GN=LOC404103 PE=4 SV= | A0A3Q1M0F4 [4] | Protease Inhibitor | 16 kDa | 9.1 | 0 | 0.00% | 0 | 0.00% | 2 | 0.03% | 0 | 0.00% | 0 |
| Streptavidin OS=Streptomyces avidinii OX=1895 | P22629 | Other | 19 kDa | 6.1 | 0 | 0.00% | 0 | 0.00% | 0 | 0.00% | 0 | 0.00% | 0 |
| Fibulin-1 OS=Bos taurus OX=9913 GN=FBLN1 PE=1 SV=1 | A0A3Q1MG73 (+2) | Coagulation | 74 kDa | 4.9 | 2 | 0.04% | 3 | 0.05% | 0 | 0.00% | 0 | 0.00% | 0 |
| Cluster of Polymeric immunoglobulin receptor OS=Bos taurus OX=9913 GN=PIGR PE=4 SV=3 (F1MR22) | F1MR22 [2] | Immunoglobulin | 83 kDa | 6.86 | 0 | 0.00% | 0 | 0.00% | 5 | 0.09% | 0 | 0.00% | 0 |
| Cluster of Glyceraldehyde-3-phosphate dehydrogenase OS=Bos taurus OX=9913 GN=GAPDH PE=1 SV=4 (P1( | P10096 [2] | Enzyme/Protease | 36 kDa | 8.52 | 3 | 0.06% | 2 | 0.03% | 3 | 0.05% | 0 | 0.00% | 0 |
| Cluster of Ubiquitin-60S ribosomal protein L40 OS=Bos taurus OX=9913 GN=UBA52 PE=1 SV=2 (P63048) | P63048 [2] | Other | 15 kDa | 6.56 | 0 | 0.00% | 0 | 0.00% | 4 | 0.07% | 0 | 0.00% | 0 |
| Elongation factor 1-alpha 1 OS=Bos taurus OX=9913 GN=EEF1A1 PE=1 SV=1 | P68103 | Other | 50 kDa | 9.1 | 3 | 0.06% | 0 | 0.00% | 0 | 0.00% | 0 | 0.00% | 0 |
| Uncharacterized protein OS=Bos taurus OX=9913 GN=TNC PE=3 SV=1 | A0A3Q1LIW2 (+6) | Extracellular Matrix | 221 kDa | 4.94 | 0 | 0.00% | 3 | 0.05% | 0 | 0.00% | 2 | 0.04% | 0 |
| Chondroadherin OS=Bos taurus OX=9913 GN=CHAD PE=4 SV=2 | F1MYE4 | Cartilage Related | 41 kDa | 9.43 | 0 | 0.00% | 0 | 0.00% | 4 | 0.07% | 3 | 0.05% | 0 |
| Actin-depolymerizing factor OS=Bos taurus OX=9913 GN=GSN PE=3 SV=1 | A0A3Q1MIS1 | Structural/Cytoskeleton | 90 kDa | 5.98 | 0 | 0.00% | 7 | 0.12% | 0 | 0.00% | 0 | 0.00% | 3 |
| Uncharacterized protein OS=Bos taurus OX=9913 GN=LAMB2 PE=4 SV=3 | E1BDK6 | Extracellular Matrix | 196 kDa | 6.23 | 0 | 0.00% | 0 | 0.00% | 0 | 0.00% | 3 | 0.05% | 0 |
| Alpha-1B-glycoprotein OS=Bos taurus OX=9913 GN=A1BG PE=1 SV=1 | A0A3Q1MJT2 (+1) | Immunity/Immunity Related | 62 kDa | 5.76 | 0 | 0.00% | 0 | 0.00% | 0 | 0.00% | 0 | 0.00% | 0 |
| Thrombospondin 3 OS=Bos taurus OX=9913 GN=THBS3 PE=3 SV=1 | A0A3Q1MQX0 (+1) | Cell-Cell Adhesion | 104 kDa | 4.43 | 0 | 0.00% | 0 | 0.00% | 3 | 0.05% | 0 | 0.00% | 0 |
| Uncharacterized protein OS=Bos taurus OX=9913 PE=1 SV=1 | A0A3Q1LRW4 | Immunity/Immunity Related | 47 kDa | 8.2 | 0 | 0.00% | 0 | 0.00% | 0 | 0.00% | 0 | 0.00% | 4 |

| | | | | | | | | | | | | | |
|---|---|---|---|---|---|---|---|---|---|---|---|---|---|
| Serpin family G member 1 OS=Bos taurus OX=9913 GN=SERPING1 PE=3 SV=3 | E1BMJ0 (+1) | Complement | 52 kDa | 6.28 | 0 | 0.00% | 3 | 0.05% | 0 | 0.00% | 0 | 0.00% | 3 |
| Lecithin-cholesterol acyltransferase OS=Bos taurus OX=9913 GN=LCAT PE=2 SV=1 | Q2KIW4 | Enzyme/Protease | 50 kDa | 5.96 | 3 | 0.06% | 0 | 0.00% | 0 | 0.00% | 0 | 0.00% | 0 |
| Collectin-11 OS=Bos taurus OX=9913 GN=COLEC11 PE=4 SV=1 | A0A3Q1M944 (+2) | Immunity/Immunity Related | 29 kDa | 5.62 | 2 | 0.04% | 0 | 0.00% | 3 | 0.05% | 0 | 0.00% | 2 |
| Cluster of Ig-like domain-containing protein OS=Bos taurus OX=9913 PE=4 SV=2 (G5E604) | G5E604 [3] | Immunoglobulin | 13 kDa | 6.53 | 2 | 0.04% | 0 | 0.00% | 0 | 0.00% | 0 | 0.00% | 0 |
| Integrin beta OS=Bos taurus OX=9913 GN=ITGB1 PE=3 SV=1 | A0A452DIS2 (+1) | Cell-Cell Adhesion | 88 kDa | 5.29 | 0 | 0.00% | 0 | 0.00% | 0 | 0.00% | 0 | 0.00% | 2 |
| Amine oxidase OS=Bos taurus OX=9913 GN=AOC1 PE=2 SV=1 | Q3MHQ1 | Enzyme/Protease | 86 kDa | 6.86 | 0 | 0.00% | 0 | 0.00% | 0 | 0.00% | 0 | 0.00% | 5 |
| Uncharacterized protein OS=Bos taurus OX=9913 GN=LOC525947 PE=3 SV=1 | A0A3Q1MUV9 (+2) | Transport/Carrier/Binding | 73 kDa | 7.88 | 0 | 0.00% | 0 | 0.00% | 0 | 0.00% | 0 | 0.00% | 3 |
| Alpha-S2-casein OS=Bos taurus OX=9913 GN=CSN1S2 | P02663 | Other | 26 kDa | 8.34 | 0 | 0.00% | 0 | 0.00% | 0 | 0.00% | 0 | 0.00% | 0 |
| Anion exchange protein OS=Bos taurus OX=9913 GN=SLC4A1 PE=3 SV=1 | A0A3Q1MR36 (+3) | Transport/Carrier/Binding | 101 kDa | 5.21 | 2 | 0.04% | 0 | 0.00% | 2 | 0.03% | 0 | 0.00% | 0 |
| Lamin A/C OS=Bos taurus OX=9913 GN=LMNA PE=1 SV=1 | F1MYG5 | Other | 74 kDa | 6.94 | 0 | 0.00% | 3 | 0.05% | 0 | 0.00% | 0 | 0.00% | 0 |
| Biglycan OS=Bos taurus OX=9913 GN=BGN PE=1 SV=3 | P21809 | Extracellular Matrix | 42 kDa | 8.13 | 5 | 0.11% | 0 | 0.00% | 0 | 0.00% | 0 | 0.00% | 0 |
| Cluster of Periostin OS=Bos taurus OX=9913 GN=POSTN PE=1 SV=1 (A0A3Q1MF21) | A0A3Q1MF21 [3] | Extracellular Matrix | 90 kDa | 7.33 | 0 | 0.00% | 0 | 0.00% | 0 | 0.00% | 0 | 0.00% | 3 |
| Ig gamma chain C region OS=Oryctolagus cuniculus OX=9986 | P01870 | Immunoglobulin | 35 kDa | 8.62 | 0 | 0.00% | 4 | 0.07% | 0 | 0.00% | 0 | 0.00% | 0 |
| Alpha-S1-casein OS=Bos taurus OX=9913 GN=CSN1S1 PE=1 SV=1 | A0A3Q1MJE5 (+2) | Transport/Carrier/Binding | 23 kDa | 4.68 | 0 | 0.00% | 0 | 0.00% | 0 | 0.00% | 0 | 0.00% | 0 |

**Supplemental Table 4: Mass spectroscopy data for dendrimers incubated in FBS stock**

| Identified Proteins (310) | Accession Number | Biological Process | Molecular Weight | Theoreticl pI | FBS stock Total Spectrum Counts | FBS stock Percentage of Total Spectral Counts |
|---|---|---|---|---|---|---|
| Cluster of Albumin OS=Bos taurus OX=9913 GN=ALB PE=4 SV=1 (A0A140T897) | A0A140T897 [2] | Transport/Carrier/Binding | 69 kDa | 5.77 | 2484 | 22.66% |
| Alpha-1-antiproteinase OS=Bos taurus OX=9913 GN=SERPINA1 PE=1 SV=1 | P34955 | Protease Inhibitor | 46 kDa | 5.98 | 833 | 7.60% |
| Alpha-2-HS-glycoprotein OS=Bos taurus OX=9913 GN=AHSG PE=1 SV=2 | P12763 | Protease Inhibitor | 38 kDa | 5.1 | 772 | 7.04% |
| Cluster of Beta-1 metal-binding globulin OS=Bos taurus OX=9913 GN=TF PE=1 SV=2 (G3X6N3) | G3X6N3 [3] | Transport/Carrier/Binding | 78 kDa | 6.63 | 743 | 6.78% |
| Alpha-2-macroglobulin OS=Bos taurus OX=9913 GN=A2M PE=1 SV=2 | Q7SIH1 | Protease Inhibitor | 168 kDa | 5.68 | 537 | 4.90% |
| Complement C3 OS=Bos taurus OX=9913 GN=C3 PE=1 SV=2 | Q2UVX4 | Complement | 187 kDa | 6.37 | 353 | 3.22% |
| Apolipoprotein B OS=Bos taurus OX=9913 GN=APOB PE=1 SV=3 | E1BNR0 | Apolipoprotein | 516 kDa | 6.24 | 275 | 2.51% |
| Alpha-fetoprotein OS=Bos taurus OX=9913 GN=AFP PE=2 SV=1 | Q3SZ57 | Transport/Carrier/Binding | 69 kDa | 5.92 | 268 | 2.44% |
| Cluster of Gc-globulin OS=Bos taurus OX=9913 GN=GC PE=4 SV=2 (F1N5M2) | F1N5M2 [2] | Transport/Carrier/Binding | 53 kDa | 5.24 | 225 | 2.05% |
| Cluster of C4a anaphylatoxin OS=Bos taurus OX=9913 GN=LOC107131209 PE=4 SV=3 (F1MVK1) | F1MVK1 [3] | Protease Inhibitor | 192 kDa | 7.26 | 174 | 1.59% |
| Serpin A3-1 OS=Bos taurus OX=9913 GN=SERPINA3-1 PE=3 SV=1 | A0A452DJK6 (+1) | Protease Inhibitor | 52 kDa | 5.99 | 161 | 1.47% |
| Cluster of Plasminogen OS=Bos taurus OX=9913 GN=PLG PE=3 SV=2 (E1B726) | E1B726 [2] | Coagulation | 91 kDa | 7.71 | 153 | 1.40% |
| Cluster of Inter-alpha-trypsin inhibitor heavy chain H4 OS=Bos taurus OX=9913 GN=ITIH4 PE=3 SV=3 (F1MMD7) | F1MMD7 [3] | Protease Inhibitor | 102 kDa | 5.99 | 143 | 1.30% |
| ITIH2 protein OS=Bos taurus OX=9913 GN=ITIH2 PE=1 SV=1 | A5D7R6 | Protease Inhibitor | 106 kDa | 7.75 | 135 | 1.23% |
| Fetuin-B OS=Bos taurus OX=9913 GN=FETUB PE=1 SV=1 | Q58D62 | Protease Inhibitor | 43 kDa | 5.59 | 135 | 1.23% |
| Cluster of Apolipoprotein A-I OS=Bos taurus OX=9913 GN=APOA1 PE=1 SV=3 (P15497) | P15497 [2] | Apolipoprotein | 30 kDa | 5.39 | 124 | 1.13% |
| Cluster of Uncharacterized protein OS=Bos taurus OX=9913 GN=LOC506828 PE=3 SV=3 (F1MJK3) | F1MJK3 [3] | Protease Inhibitor | 163 kDa | 6.91 | 122 | 1.11% |
| Cluster of Antithrombin-III OS=Bos taurus OX=9913 GN=SERPINC1 PE=3 SV=1 (A0A3Q1NJR8) | A0A3Q1NJR8 [2] | Protease Inhibitor | 60 kDa | 8.85 | 120 | 1.09% |
| Cluster of Serpin A3-7 OS=Bos taurus OX=9913 GN=SERPINA3-7 PE=1 SV=1 (A0A0A0MP92) | A0A0A0MP92 [2] | Protease Inhibitor | 47 kDa | 5.61 | 117 | 1.07% |
| Alpha-2-antiplasmin OS=Bos taurus OX=9913 GN=SERPINF2 PE=1 SV=2 | P28800 | Protease Inhibitor | 55 kDa | 5.71 | 107 | 0.98% |
| Cluster of Hemopexin OS=Bos taurus OX=9913 GN=HPX PE=2 SV=1 (Q3SZV7) | Q3SZV7 [2] | Transport/Carrier/Binding | 52 kDa | 7.1 | 99 | 0.90% |
| Cluster of Bradykinin OS=Bos taurus OX=9913 GN=KNG1 PE=4 SV=1 (A0A140T8C8) | A0A140T8C8 [2] | Protease Inhibitor | 69 kDa | 6.09 | 93 | 0.85% |
| Inter-alpha-trypsin inhibitor heavy chain H3 OS=Bos taurus OX=9913 GN=ITIH3 PE=3 SV=1 | A0A3Q1LQ21 (+1) | Transport/Carrier/Binding | 99 kDa | 5.91 | 93 | 0.85% |
| Complement factor B OS=Bos taurus OX=9913 GN=CFB PE=1 SV=2 | P81187 | Complement | 85 kDa | 7.68 | 86 | 0.78% |
| Cluster of Complement factor H OS=Bos taurus OX=9913 GN=CFH PE=1 SV=3 (Q28085) | Q28085 [6] | Complement | 140 kDa | 6.33 | 85 | 0.78% |
| Prothrombin OS=Bos taurus OX=9913 GN=F2 PE=1 SV=2 | P00735 | Coagulation | 71 kDa | 5.4 | 85 | 0.78% |
| Hemoglobin fetal subunit beta OS=Bos taurus OX=9913 PE=1 SV=1 | P02081 | Transport/Carrier/Binding | 16 kDa | 6.51 | 83 | 0.76% |
| Cluster of Serpin A3-3 OS=Bos taurus OX=9913 GN=SERPINA3-3 PE=1 SV=2 (Q3ZEJ6) | Q3ZEJ6 [3] | Protease Inhibitor | 46 kDa | 5.73 | 82 | 0.75% |
| Angiotensinogen OS=Bos taurus OX=9913 GN=AGT PE=1 SV=2 | P01017 | Protease Inhibitor | 51 kDa | 6.49 | 67 | 0.61% |
| Alpha-1-acid glycoprotein OS=Bos taurus OX=9913 GN=ORM1 PE=2 SV=1 | Q3SZR3 (+1) | Transport/Carrier/Binding | 23 kDa | 5.67 | 62 | 0.57% |
| Serpin family D member 1 OS=Bos taurus OX=9913 GN=SERPIND1 PE=3 SV=1 | F6R4P6 | Protease Inhibitor | 63 kDa | 6.23 | 62 | 0.57% |
| Complement C5a anaphylatoxin OS=Bos taurus OX=9913 GN=C5 PE=1 SV=3 | F1MY85 | Complement | 189 kDa | 6.16 | 61 | 0.56% |
| Alpha-1B-glycoprotein OS=Bos taurus OX=9913 GN=A1BG PE=1 SV=1 | Q2KJF1 | Immunity/Immunity Related | 54 kDa | 5.37 | 60 | 0.55% |
| Uncharacterized protein OS=Bos taurus OX=9913 GN=LOC525947 PE=1 SV=1 | A0A3Q1NEQ0 | Transport/Carrier/Binding | 78 kDa | 6.8 | 58 | 0.53% |
| Apolipoprotein H OS=Bos taurus OX=9913 GN=APOH PE=4 SV=1 | A0A140T843 (+1) | Apolipoprotein | 38 kDa | 8.55 | 57 | 0.52% |
| Actin-depolymerizing factor OS=Bos taurus OX=9913 GN=GSN PE=1 SV=2 | F1N1I6 | Structural/Cytoskeleton | 92 kDa | 6.42 | 52 | 0.47% |
| Uncharacterized protein OS=Bos taurus OX=9913 PE=1 SV=1 | A0A3Q1M3L6 | Immunoglobulin | 40 kDa | 5.16 | 51 | 0.47% |
| Transthyretin OS=Bos taurus OX=9913 GN=TTR PE=1 SV=1 | O46375 | Transport/Carrier/Binding | 16 kDa | 5.91 | 51 | 0.47% |
| Hemoglobin subunit alpha OS=Bos taurus OX=9913 GN=HBA PE=1 SV=2 | P01966 | Transport/Carrier/Binding | 15 kDa | 8.19 | 49 | 0.45% |
| Afamin OS=Bos taurus OX=9913 GN=AFM PE=1 SV=1 | G3MYZ3 | Transport/Carrier/Binding | 70 kDa | 5.57 | 48 | 0.44% |
| Pigment epithelium-derived factor OS=Bos taurus OX=9913 GN=SERPINF1 PE=1 SV=1 | Q95121 | Coagulation | 46 kDa | 6.31 | 47 | 0.43% |
| Inter-alpha-trypsin inhibitor heavy chain H1 OS=Bos taurus OX=9913 GN=ITIH1 PE=1 SV=1 | Q0VCM5 | Protease Inhibitor | 101 kDa | 7.52 | 45 | 0.41% |
| Cluster of Fibronectin OS=Bos taurus OX=9913 GN=FN1 PE=1 SV=4 (P07589) | P07589 [3] | Structural/Cytoskeleton | 272 kDa | 5.28 | 42 | 0.38% |
| Cluster of Serpin family G member 1 OS=Bos taurus OX=9913 GN=SERPING1 PE=3 SV=3 (E1BMJ0) | E1BMJ0 [2] | Protease Inhibitor | 52 kDa | 6.28 | 41 | 0.37% |
| Alpha-1-microglobulin OS=Bos taurus OX=9913 GN=KIF12 PE=3 SV=3 | F1MMK9 | Other | 53 kDa | 8.87 | 41 | 0.37% |
| SERPIN domain-containing protein OS=Bos taurus OX=9913 GN=LOC784932 PE=1 SV=1 | A0A0A0MPA0 | Protease Inhibitor | 47 kDa | 5.44 | 39 | 0.36% |
| Cluster of Trypsin OS=Sus scrofa OX=9823 (P00761) | P00761 [2] | Enzyme/Protease | 24 kDa | 8.26 | 37 | 0.34% |
| Fibulin-1 OS=Bos taurus OX=9913 GN=FBLN1 PE=1 SV=1 | A0A3Q1MG73 | Coagulation | 74 kDa | 4.9 | 36 | 0.33% |
| Cluster of Actin, cytoplasmic 1 OS=Bos taurus OX=9913 GN=ACTB PE=1 SV=1 (P60712) | P60712 [4] | Structural/Cytoskeleton | 42 kDa | 5.29 | 35 | 0.32% |
| Cluster of Complement factor I OS=Bos taurus OX=9913 GN=CFI PE=1 SV=1 (A0A3Q1LGM4) | A0A3Q1LGM4 [4] | Complement | 65 kDa | 8.13 | 35 | 0.32% |
| Hemoglobin subunit beta OS=Bos taurus OX=9913 GN=HBB PE=1 SV=1 | P02070 | Transport/Carrier/Binding | 16 kDa | 7.02 | 32 | 0.29% |
| Vitronectin OS=Bos taurus OX=9913 GN=VTN PE=1 SV=1 | Q3ZBS7 | Cell-Cell Adhesion & Signaling | 54 kDa | 5.79 | 32 | 0.29% |
| Cluster of Plasma serine protease inhibitor OS=Bos taurus OX=9913 GN=SERPINA5 PE=3 SV=1 (A0A452DHZ3) | A0A452DHZ3 [3] | Coagulation | 44 kDa | 9.36 | 31 | 0.28% |
| Pantetheinase OS=Bos taurus OX=9913 GN=VNN1 PE=3 SV=1 | A0A140T891 (+1) | Enzyme/Protease | 57 kDa | 5.26 | 30 | 0.27% |
| Cluster of Thrombospondin-1 OS=Bos taurus OX=9913 GN=THBS1 PE=3 SV=1 (A0A3Q1MQV3) | A0A3Q1MQV3 [4] | Cell-Cell Adhesion & Signaling | 130 kDa | 4.69 | 28 | 0.26% |
| Apolipoprotein E OS=Bos taurus OX=9913 GN=APOE PE=3 SV=2 | A0A140T881 (+2) | Apolipoprotein | 37 kDa | 5.55 | 27 | 0.25% |
| Cluster of Clusterin OS=Bos taurus OX=9913 GN=CLU PE=1 SV=1 (P17697) | P17697 [2] | Apolipoprotein | 51 kDa | 5.72 | 26 | 0.24% |
| Leucine rich alpha-2-glycoprotein 1 OS=Bos taurus OX=9913 GN=LRG1 PE=1 SV=1 | F6RMV5 | Structural/Cytoskeleton | 39 kDa | 6.26 | 26 | 0.24% |

| | | | | | | |
|---|---|---|---|---|---|---|
| Cluster of Uncharacterized protein OS=Bos taurus OX=9913 PE=1 SV=1 (A0A3Q1M032) | A0A3Q1M032 [2] | Other | 40 kDa | 5.77 | 25 | 0.23% |
| Periostin OS=Bos taurus OX=9913 GN=POSTN PE=1 SV=1 | A0A3Q1NNK1 | Extracellular Matrix | 90 kDa | 7.91 | 25 | 0.23% |
| Fibrinogen alpha chain OS=Bos taurus OX=9913 GN=FGA PE=4 SV=1 | F6QND5 | Coagulation | 95 kDa | 5.69 | 24 | 0.22% |
| Cluster of Complement component C7 OS=Bos taurus OX=9913 GN=C7 PE=2 SV=1 (Q29RQ1) | Q29RQ1 [3] | Complement | 93 kDa | 6.9 | 24 | 0.22% |
| Serpin peptidase inhibitor, clade A (Alpha-1 antiproteinase, antitrypsin), member 7 OS=Bos taurus OX=9913 GN=SERPIN. | Q3SYR0 (+1) | Protease Inhibitor | 46 kDa | 5.51 | 23 | 0.21% |
| Uncharacterized protein OS=Bos taurus OX=9913 PE=1 SV=1 | G3N0V0 | Complement | 36 kDa | 8.05 | 21 | 0.19% |
| Cluster of Uncharacterized protein OS=Bos taurus OX=9913 PE=4 SV=3 (F1N160) | F1N160 [3] | Immunity/Immunity Related | 34 kDa | 8.86 | 21 | 0.19% |
| Plasma retinol-binding protein OS=Bos taurus OX=9913 GN=RBP4 PE=3 SV=1 | A0A3Q1MSW9 (+2) | Transport/Carrier/Binding | 29 kDa | 5.66 | 21 | 0.19% |
| Lumican OS=Bos taurus OX=9913 GN=LUM PE=1 SV=1 | Q05443 | Extracellular Matrix | 39 kDa | 5.93 | 21 | 0.19% |
| Cluster of Chitinase OS=Bos taurus OX=9913 GN=CHIA PE=3 SV=2 (F1MH27) | F1MH27 [2] | Enzyme/Protease | 52 kDa | 5.36 | 21 | 0.19% |
| Serpin A3-8 OS=Bos taurus OX=9913 GN=SERPINA3-8 PE=2 SV=1 | A6QPQ2 | Protease Inhibitor | 47 kDa | 5.49 | 19 | 0.17% |
| Cluster of Complement component C9 OS=Bos taurus OX=9913 GN=C9 PE=2 SV=1 (Q3MHN2) | Q3MHN2 [2] | Complement | 62 kDa | 5.57 | 19 | 0.17% |
| Cartilage oligomeric matrix protein OS=Bos taurus OX=9913 GN=COMP PE=1 SV=2 | P35445 | Extracellular Matrix | 82 kDa | 4.36 | 18 | 0.16% |
| SERPINA10 protein OS=Bos taurus OX=9913 GN=SERPINA10 PE=2 SV=1 | A5PJ69 | Protease Inhibitor | 52 kDa | 6.05 | 18 | 0.16% |
| Plasma kallikrein OS=Bos taurus OX=9913 GN=KLKB1 PE=2 SV=1 | Q2KJ63 | Complement | 71 kDa | 8.62 | 18 | 0.16% |
| C4b-binding protein alpha chain OS=Bos taurus OX=9913 GN=C4BPA PE=2 SV=1 | Q28065 | Complement | 69 kDa | 5.66 | 16 | 0.15% |
| Corticosteroid-binding globulin OS=Bos taurus OX=9913 GN=SERPINA6 PE=3 SV=1 | E1BF81 | Protease Inhibitor | 45 kDa | 5.54 | 16 | 0.15% |
| Cadherin-5 OS=Bos taurus OX=9913 GN=CDH5 PE=4 SV=2 | G3MZL1 (+1) | Cell-Cell Adhesion & Signaling | 87 kDa | 5.3 | 16 | 0.15% |
| HGF activator OS=Bos taurus OX=9913 GN=HGFAC PE=4 SV=3 | E1BCW0 | Coagulation | 70 kDa | 7.66 | 16 | 0.15% |
| Cluster of Tetranectin OS=Bos taurus OX=9913 GN=CLEC3B PE=4 SV=1 (A0A452DIU4) | A0A452DIU4 [4] | Transport/Carrier/Binding | 31 kDa | 6.52 | 15 | 0.14% |
| Adiponectin OS=Bos taurus OX=9913 GN=ADIPOQ PE=4 SV=1 | A0A3Q1M564 (+1) | Cell-Cell Adhesion & Signaling | 35 kDa | 5.36 | 15 | 0.14% |
| Cluster of Uncharacterized protein OS=Bos taurus OX=9913 PE=1 SV=3 (F1MZ96) | F1MZ96 [2] | Immunity/Immunity Related | 27 kDa | 5.94 | 14 | 0.13% |
| Cystatin-C OS=Bos taurus OX=9913 GN=CST3 PE=1 SV=2 | P01035 | Protease Inhibitor | 16 kDa | 9.03 | 14 | 0.13% |
| Collagen type VI alpha 1 chain OS=Bos taurus OX=9913 GN=COL6A1 PE=1 SV=1 | E1BI98 | Extracellular Matrix | 109 kDa | 5.18 | 14 | 0.13% |
| CPN2 protein OS=Bos taurus OX=9913 GN=CPN2 PE=2 SV=1 | A6QP30 | Enzyme/Protease | 60 kDa | 5.66 | 14 | 0.13% |
| Galectin-3-binding protein OS=Bos taurus OX=9913 GN=LGALS3BP PE=1 SV=1 | A7E3W2 | Immunity/Immunity Related | 62 kDa | 5.25 | 14 | 0.13% |
| Apolipoprotein A-IV OS=Bos taurus OX=9913 GN=APOA4 PE=3 SV=2 | F1N3Q7 | Apolipoprotein | 52 kDa | 5.5 | 13 | 0.12% |
| Alpha-amylase OS=Bos taurus OX=9913 GN=AMY2B PE=3 SV=1 | F1MJQ3 | Enzyme/Protease | 57 kDa | 5.87 | 13 | 0.12% |
| Coagulation factor V OS=Bos taurus OX=9913 GN=F5 PE=3 SV=3 | F1N0I3 | Coagulation | 240 kDa | 5.74 | 13 | 0.12% |
| Cluster of Fibrinogen beta chain OS=Bos taurus OX=9913 GN=FGB PE=4 SV=1 (A0A3Q1MG04) | A0A3Q1MG04 [3] | Coagulation | 57 kDa | 8.33 | 12 | 0.11% |
| Collagen alpha-2(I) chain OS=Bos taurus OX=9913 GN=COL1A2 PE=1 SV=1 | A0A3Q1LZN8 (+2) | Extracellular Matrix | 120 kDa | 9.3 | 12 | 0.11% |
| C-X-C motif chemokine OS=Bos taurus OX=9913 GN=PPBP PE=1 SV=2 | F1MD83 | Immunity/Immunity Related | 32 kDa | 9.41 | 12 | 0.11% |
| Vitamin K-dependent protein S OS=Bos taurus OX=9913 GN=PROS1 PE=4 SV=1 | A0A3Q1MMH1 (+1) | Coagulation | 71 kDa | 5.42 | 11 | 0.10% |
| Apolipoprotein A-II OS=Bos taurus OX=9913 GN=APOA2 PE=1 SV=2 | P81644 | Apolipoprotein | 11 kDa | 5.34 | 11 | 0.10% |
| Cluster of Thrombospondin-4 OS=Bos taurus OX=9913 GN=THBS4 PE=3 SV=1 (A0A452DJ62) | A0A452DJ62 [2] | Cell-Cell Adhesion & Signaling | 106 kDa | 4.41 | 11 | 0.10% |
| Cation-independent mannose-6-phosphate receptor OS=Bos taurus OX=9913 GN=IGF2R PE=4 SV=3 | F1MIE6 | Transport/Carrier/Binding | 275 kDa | 5.62 | 10 | 0.09% |
| CTRB1 protein OS=Bos taurus OX=9913 GN=CTRB1 PE=2 SV=1 | A5PJB8 | Enzyme/Protease | 28 kDa | 5.17 | 10 | 0.09% |
| Gluconolactonase OS=Bos taurus OX=9913 GN=RGN PE=3 SV=1 | A0A3Q1MLX2 (+1) | Enzyme/Protease | 34 kDa | 5.54 | 10 | 0.09% |
| Fibrinogen gamma-B chain OS=Bos taurus OX=9913 GN=FGG PE=4 SV=1 | F1MGU7 (+1) | Coagulation | 50 kDa | 5.38 | 9 | 0.08% |
| Cluster of Christmas factor OS=Bos taurus OX=9913 GN=F9 PE=4 SV=3 (F1MBC5) | F1MBC5 [3] | Enzyme/Protease | 50 kDa | 5.9 | 9 | 0.08% |
| SHBG protein OS=Bos taurus OX=9913 GN=SHBG PE=2 SV=1 | A5PKC2 | Transport/Carrier/Binding | 43 kDa | 5.43 | 9 | 0.08% |
| Complement component 8 subunit beta OS=Bos taurus OX=9913 GN=C8B PE=3 SV=3 | F1N102 | Complement | 67 kDa | 8.15 | 8 | 0.07% |
| Complement C8 alpha chain OS=Bos taurus OX=9913 GN=C8A PE=3 SV=1 | F1MX87 | Complement | 66 kDa | 6.24 | 8 | 0.07% |
| Uncharacterized protein OS=Bos taurus OX=9913 PE=4 SV=2 | G3N0S9 | Complement | 21 kDa | 5.37 | 8 | 0.07% |
| Apolipoprotein M OS=Bos taurus OX=9913 GN=APOM PE=3 SV=1 | A0A3Q1MB70 (+1) | Apolipoprotein | 25 kDa | 6.89 | 8 | 0.07% |
| Collagen alpha-1(I) chain OS=Bos taurus OX=9913 GN=COL1A1 PE=1 SV=3 | P02453 | Extracellular Matrix | 139 kDa | 9.28 | 8 | 0.07% |
| Insulin-like growth factor-binding protein 2 OS=Bos taurus OX=9913 GN=IGFBP2 PE=1 SV=2 | P13384 | Protease Inhibitor | 34 kDa | 6.93 | 8 | 0.07% |
| Protein HP-20 homolog OS=Bos taurus OX=9913 PE=2 SV=1 | Q2KIT0 | Structural/Cytoskeleton | 21 kDa | 8.86 | 7 | 0.06% |
| Apolipoprotein D OS=Bos taurus OX=9913 GN=APOD PE=3 SV=3 | F1MS32 (+1) | Apolipoprotein | 24 kDa | 5.07 | 7 | 0.06% |
| Insulin-like growth factor binding protein acid labile subunit OS=Bos taurus OX=9913 GN=IGFALS PE=2 SV=1 | Q09TE3 | Structural/Cytoskeleton | 66 kDa | 6.73 | 7 | 0.06% |
| Collectin-43 OS=Bos taurus OX=9913 GN=CL43 PE=1 SV=2 | P42916 | Immunity/Immunity Related | 34 kDa | 5.04 | 7 | 0.06% |
| Elongation factor 1-alpha 1 OS=Bos taurus OX=9913 GN=EEF1A1 PE=1 SV=1 | P68103 | Other | 50 kDa | 9.1 | 7 | 0.06% |
| Extracellular matrix protein 1 OS=Bos taurus OX=9913 GN=ECM1 PE=4 SV=1 | A0A3Q1M5Q6 | Extracellular Matrix | 63 kDa | 6.6 | 6 | 0.05% |
| Phospholipid transfer protein OS=Bos taurus OX=9913 GN=PLTP PE=2 SV=1 | Q58DL9 | Immunity/Immunity Related | 56 kDa | 6.32 | 6 | 0.05% |
| Apolipoprotein C-III OS=Bos taurus OX=9913 GN=APOC3 PE=1 SV=2 | P19035 | Apolipoprotein | 11 kDa | 4.73 | 6 | 0.05% |
| Coagulation factor X OS=Bos taurus OX=9913 GN=F10 PE=4 SV=1 | A0A452DIP8 (+1) | Coagulation | 67 kDa | 5.96 | 6 | 0.05% |
| Adenosylhomocysteinase OS=Bos taurus OX=9913 GN=AHCY PE=1 SV=1 | A0A3Q1LW27 (+1) | Enzyme/Protease | 47 kDa | 5.89 | 6 | 0.05% |
| Coagulation factor XIII B chain OS=Bos taurus OX=9913 GN=F13B PE=4 SV=1 | A0A3Q1MWQ1 (+1) | Coagulation | 68 kDa | 6 | 6 | 0.05% |
| Hepatocyte growth factor-like protein OS=Bos taurus OX=9913 GN=MST1 PE=3 SV=2 | E1BDW7 (+1) | Enzyme/Protease | 80 kDa | 8.42 | 6 | 0.05% |
| Cluster of Ceruloplasmin OS=Bos taurus OX=9913 GN=CP PE=1 SV=1 (A0A3Q1NJB1) | A0A3Q1NJB1 [2] | Transport/Carrier/Binding | 121 kDa | 5.67 | 5 | 0.05% |
| Cluster of Histidine-rich glycoprotein OS=Bos taurus OX=9913 GN=HRG PE=4 SV=3 (F1MKS5) | F1MKS5 [2] | Protease Inhibitor | 61 kDa | 7.14 | 5 | 0.05% |
| Protein HP-25 homolog 2 OS=Bos taurus OX=9913 PE=2 SV=1 | Q2KIU3 | Structural/Cytoskeleton | 23 kDa | 5.29 | 5 | 0.05% |
| Cluster of Ig-like domain-containing protein OS=Bos taurus OX=9913 PE=4 SV=1 (A0A3Q1ML26) | A0A3Q1ML26 [6] | Immunoglobulin | 23 kDa | 9.08 | 5 | 0.05% |
| Amine oxidase OS=Bos taurus OX=9913 GN=LOC100138645 PE=3 SV=1 | A0A452DHX8 | Other | 87 kDa | 5.56 | 5 | 0.05% |
| Carboxypeptidase B2 OS=Bos taurus OX=9913 GN=CPB2 PE=3 SV=1 | A0A3Q1N1A7 (+1) | Enzyme/Protease | 45 kDa | 8.69 | 5 | 0.05% |
| ApoN protein OS=Bos taurus OX=9913 GN=APON PE=2 SV=1 | Q2KIH2 | Apolipoprotein | 29 kDa | 5.36 | 5 | 0.05% |
| Cluster of von Willebrand factor OS=Bos taurus OX=9913 GN=VWF PE=4 SV=1 (A0A3Q1LLU1) | A0A3Q1LLU1 [3] | Coagulation | 308 kDa | 5.35 | 5 | 0.05% |

| | | | | | | |
|---|---|---|---|---|---|---|
| Collagen alpha-1(III) chain OS=Bos taurus OX=9913 GN=COL3A1 PE=2 SV=1 | Q08E14 | Extracellular Matrix | 138 kDa | 5.96 | 5 | 0.05% |
| Glyceraldehyde-3-phosphate dehydrogenase OS=Bos taurus OX=9913 GN=GAPDH PE=1 SV=4 | P10096 | Enzyme/Protease | 36 kDa | 8.52 | 5 | 0.05% |
| Mannose-binding protein C OS=Bos taurus OX=9913 GN=MBL PE=2 SV=1 | O02659 | Complement | 26 kDa | 5.11 | 5 | 0.05% |
| Myoglobin OS=Bos taurus OX=9913 GN=MB PE=1 SV=3 | P02192 | Transport/Carrier/Binding | 17 kDa | 6.97 | 5 | 0.05% |
| Primary amine oxidase, liver isozyme OS=Bos taurus OX=9913 PE=1 SV=1 | Q29437 | Enzyme/Protease | 85 kDa | 5.58 | 4 | 0.04% |
| Uncharacterized protein OS=Bos taurus OX=9913 GN=LOC528040 PE=4 SV=3 | E1B805 | Other | 186 kDa | 6.54 | 4 | 0.04% |
| Cluster of CD5 molecule like OS=Bos taurus OX=9913 GN=CD5L PE=2 SV=1 (A6QNW7) | A6QNW7 [3] | Immunity/Immunity Related | 50 kDa | 5.24 | 4 | 0.04% |
| Beta-2-microglobulin OS=Bos taurus OX=9913 PE=3 SV=1 | A0A3Q1MU93 (+1) | Immunity/Immunity Related | 14 kDa | 9.1 | 4 | 0.04% |
| Ig-like domain-containing protein OS=Bos taurus OX=9913 PE=4 SV=2 | G3MXG6 | Immunoglobulin | 15 kDa | 8.56 | 4 | 0.04% |
| Ig-like domain-containing protein OS=Bos taurus OX=9913 PE=4 SV=1 | A0A3Q1LSF0 (+1) | Immunoglobulin | 15 kDa | 9.82 | 4 | 0.04% |
| Complement factor D OS=Bos taurus OX=9913 GN=CFD PE=2 SV=1 | Q3T0A3 | Complement | 28 kDa | 6.85 | 4 | 0.04% |
| Complement factor properdin OS=Bos taurus OX=9913 GN=CFP PE=4 SV=1 | A0A3Q1MHU8 (+1) | Complement | 50 kDa | 8.33 | 4 | 0.04% |
| Vitamin K-dependent protein C OS=Bos taurus OX=9913 GN=PROC PE=4 SV=2 | A0A140T851 (+2) | Coagulation | 51 kDa | 6.06 | 4 | 0.04% |
| Carboxypeptidase N catalytic chain OS=Bos taurus OX=9913 GN=CPN1 PE=3 SV=1 | G5E5V0 (+1) | Enzyme/Protease | 53 kDa | 8.76 | 4 | 0.04% |
| Cluster of 2-phospho-D-glycerate hydro-lyase OS=Bos taurus OX=9913 GN=ENO1 PE=3 SV=1 (A0A3Q1MXQ0) | A0A3Q1MXQ0 [5] | Enzyme/Protease | 54 kDa | 9.12 | 4 | 0.04% |
| Hyaluronan-binding protein 2 OS=Bos taurus OX=9913 GN=HABP2 PE=2 SV=1 | Q5E9Z2 | Extracellular Matrix | 62 kDa | 5.94 | 4 | 0.04% |
| Phosphatidylinositol-glycan-specific phospholipase D OS=Bos taurus OX=9913 GN=GPLD1 PE=1 SV=1 | P80109 | Enzyme/Protease | 93 kDa | 6.06 | 4 | 0.04% |
| Sulfhydryl oxidase OS=Bos taurus OX=9913 GN=QSOX1 PE=3 SV=3 | F1MM32 | Enzyme/Protease | 67 kDa | 9.29 | 4 | 0.04% |
| Tubulin beta-5 chain OS=Bos taurus OX=9913 GN=TUBB5 PE=2 SV=1 | Q2KJD0 | Structural/Cytoskeleton | 50 kDa | 4.78 | 4 | 0.04% |
| Protein HP-25 homolog 1 OS=Bos taurus OX=9913 PE=1 SV=1 | Q2KIX7 | Structural/Cytoskeleton | 23 kDa | 5.71 | 3 | 0.03% |
| EGF containing fibulin extracellular matrix protein 1 OS=Bos taurus OX=9913 GN=EFEMP1 PE=1 SV=1 | A0A3Q1MIU1 (+2) | Extracellular Matrix | 50 kDa | 4.83 | 3 | 0.03% |
| Monocyte differentiation antigen CD14 OS=Bos taurus OX=9913 GN=CD14 PE=2 SV=1 | A6QNL0 (+1) | Immunity/Immunity Related | 40 kDa | 5.24 | 3 | 0.03% |
| Regakine-1 OS=Bos taurus OX=9913 PE=1 SV=2 | P82943 | Immunity/Immunity Related | 10 kDa | 8.81 | 3 | 0.03% |
| Uncharacterized protein OS=Bos taurus OX=9913 GN=C8G PE=4 SV=1 | A0A3Q1NH68 | Other | 35 kDa | 10.52 | 3 | 0.03% |
| Transforming growth factor-beta-induced protein ig-h3 OS=Bos taurus OX=9913 GN=TGFBI PE=1 SV=2 | P55906 | Cell-Cell Adhesion & Signaling | 74 kDa | 6.69 | 3 | 0.03% |
| Coagulation factor XII OS=Bos taurus OX=9913 GN=F12 PE=4 SV=2 | F1MTT3 (+1) | Coagulation | 66 kDa | 7.61 | 3 | 0.03% |
| Cysteine-rich secretory protein 2 OS=Bos taurus OX=9913 GN=CRISP3 PE=2 SV=1 | Q3ZCL0 | Immunity/Immunity Related | 27 kDa | 8.55 | 3 | 0.03% |
| Ferritin OS=Bos taurus OX=9913 PE=3 SV=1 | A0A3Q1MM35 (+1) | Other | 20 kDa | 5.87 | 3 | 0.03% |
| Glutathione S-transferase A2 OS=Bos taurus OX=9913 GN=GSTA2 PE=2 SV=4 | O18879 | Enzyme/Protease | 26 kDa | 8.67 | 3 | 0.03% |
| Glutathione S-transferase OS=Bos taurus OX=9913 GN=GSTA1 PE=3 SV=1 | A0A3Q1MEC8 | Enzyme/Protease | 32 kDa | 7.01 | 3 | 0.03% |
| Ig-like domain-containing protein OS=Bos taurus OX=9913 PE=4 SV=2 | G3MY71 (+1) | Immunoglobulin | 23 kDa | 9.42 | 3 | 0.03% |
| Insulin-like growth factor II OS=Bos taurus OX=9913 GN=IGF2 PE=3 SV=1 | A0A3Q1LM00 (+3) | Other | 21 kDa | 9.27 | 3 | 0.03% |
| Peptidyl-prolyl cis-trans isomerase A OS=Bos taurus OX=9913 GN=PPIA PE=1 SV=2 | P62935 | Enzyme/Protease | 18 kDa | 8.34 | 3 | 0.03% |
| Phosphatidylethanolamine-binding protein 1 OS=Bos taurus OX=9913 GN=PEBP1 PE=1 SV=2 | P13696 | Protease Inhibitor | 21 kDa | 7.39 | 3 | 0.03% |
| Profilin-1 OS=Bos taurus OX=9913 GN=PFN1 PE=1 SV=2 | P02584 | Structural/Cytoskeleton | 15 kDa | 8.5 | 3 | 0.03% |
| VASN protein OS=Bos taurus OX=9913 GN=VASN PE=2 SV=1 | A4IFA5 | Cell-Cell Adhesion & Signaling | 72 kDa | 8.63 | 3 | 0.03% |
| Complement component C6 OS=Bos taurus OX=9913 GN=C6 PE=3 SV=2 | F1MM86 | Complement | 105 kDa | 6.74 | 2 | 0.02% |
| Zinc-alpha-2-glycoprotein OS=Bos taurus OX=9913 GN=AZGP1 PE=3 SV=1 | A0A452DK44 (+1) | Immunity/Immunity Related | 34 kDa | 5.11 | 2 | 0.02% |
| Argininosuccinate synthase OS=Bos taurus OX=9913 GN=ASS1 PE=1 SV=1 | A0A3Q1MM52 (+1) | Enzyme/Protease | 48 kDa | 6.81 | 2 | 0.02% |
| CD109 molecule OS=Bos taurus OX=9913 GN=CD109 PE=3 SV=1 | A0A3Q1LI93 (+6) | Protease Inhibitor | 163 kDa | 5.5 | 2 | 0.02% |
| Cluster of 15-oxoprostaglandin 13-reductase OS=Bos taurus OX=9913 GN=PTGR1 PE=3 SV=1 (F1N2W0) | F1N2W0 [3] | Enzyme/Protease | 36 kDa | 7.65 | 2 | 0.02% |
| Coagulation factor XI OS=Bos taurus OX=9913 GN=F11 PE=2 SV=1 | Q5NTB3 | Coagulation | 70 kDa | 8.32 | 2 | 0.02% |
| Collagen alpha-1(II) chain OS=Bos taurus OX=9913 GN=COL2A1 PE=1 SV=4 | P02459 | Extracellular Matrix | 142 kDa | 8.38 | 2 | 0.02% |
| Mimecan OS=Bos taurus OX=9913 GN=OGN PE=2 SV=1 | A5D9E8 (+1) | Extracellular Matrix | 34 kDa | 5.21 | 2 | 0.02% |
| Osteomodulin OS=Bos taurus OX=9913 GN=OMD PE=4 SV=1 | G3X6Y4 (+1) | Extracellular Matrix | 49 kDa | 5.21 | 2 | 0.02% |
| Osteonectin OS=Bos taurus OX=9913 GN=SPARC PE=3 SV=1 | A0A3Q1N541 | Extracellular Matrix | 44 kDa | 5.05 | 2 | 0.02% |
| Thyroglobulin OS=Bos taurus OX=9913 GN=TG PE=3 SV=1 | A0A3Q1LKN2 (+5) | Other | 275 kDa | 5.42 | 2 | 0.02% |
| Triosephosphate isomerase OS=Bos taurus OX=9913 GN=TPI1 PE=3 SV=1 | A0A452DIX3 (+1) | Enzyme/Protease | 31 kDa | 5.89 | 2 | 0.02% |
| Tubulin alpha chain OS=Bos taurus OX=9913 GN=TUBA4A PE=3 SV=1 | A0A3Q1M1Z2 (+2) | Structural/Cytoskeleton | 46 kDa | 5.91 | 2 | 0.02% |
